# Whole-genome sequencing of 1,171 elderly admixed individuals from the largest Latin American metropolis (São Paulo, Brazil)

**DOI:** 10.1101/2020.09.15.298026

**Authors:** Michel S. Naslavsky, Marilia O. Scliar, Guilherme L. Yamamoto, Jaqueline Yu Ting Wang, Stepanka Zverinova, Tatiana Karp, Kelly Nunes, José Ricardo Magliocco Ceroni, Diego Lima de Carvalho, Carlos Eduardo da Silva Simões, Daniel Bozoklian, Ricardo Nonaka, Nayane dos Santos Brito Silva, Andreia da Silva Souza, Heloísa de Souza Andrade, Marília Rodrigues Silva Passos, Camila Ferreira Bannwart Castro, Celso T. Mendes-Junior, Rafael L. V. Mercuri, Thiago L. A. Miller, Jose Leonel Buzzo, Fernanda O. Rego, Nathalia M Araújo, Wagner CS Magalhães, Regina Célia Mingroni-Netto, Victor Borda, Heinner Guio, Mauricio L Barreto, Maria Fernanda Lima-Costa, Bernardo L Horta, Eduardo Tarazona-Santos, Diogo Meyer, Pedro A. F. Galante, Victor Guryev, Erick C. Castelli, Yeda A. O. Duarte, Maria Rita Passos-Bueno, Mayana Zatz

**Author notes:** Authors contributed equally (Michel S. Naslavsky, Marilia O. Scliar, Guilherme L. Yamamoto). Corresponding authors, Rua do Matão, 277/211, ZIP 05508090, São Paulo - SP, Brazil, Rua do Matão, Tv. 13, 106, ZIP 05508090, São Paulo - SP, Brazil.

## Abstract

As whole-genome sequencing (WGS) becomes the gold standard tool for studying population genomics and medical applications, data on diverse non-European and admixed individuals are still scarce. Here, we present a high-coverage WGS dataset of 1,171 highly admixed elderly Brazilians from a census-based cohort, providing over 76 million variants, of which ~2 million are absent from large public databases. WGS enabled identifying ~2,000 novel mobile element insertions, nearly 5Mb of genomic segments absent from human genome reference, and over 140 novel alleles from HLA genes. We reclassified and curated nearly four hundred variant's pathogenicity assertions in genes associated with dominantly inherited Mendelian disorders and calculated the incidence for selected recessive disorders, demonstrating the clinical usefulness of the present study. Finally, we observed that whole-genome and HLA imputation could be significantly improved compared to available datasets since rare variation represents the largest proportion of input from WGS. These results demonstrate that even smaller sample sizes of underrepresented populations bring relevant data for genomic studies, especially when exploring analyses allowed only by WGS.

## INTRODUCTION

Whole-genome sequencing (WGS) of a large number of individuals can reveal rare variants in known disease genes^1–4^, improve identification of novel genes and pathways associated with phenotypes^5^ and identify genomic regions not represented on reference genomes^6^. Most importantly, ancestry diversity is critical to elucidate differences in disease’s genomic architecture and improve signals detected by previous studies, since non-European and admixed populations harbor specific variants^7–9^, which are still vastly underrepresented in genomic studies^10^. The lack of diversity leads to a significant bias on the primary resource for precision medicine and consequently less accurate tests on non-European descent individuals, potentially increasing health disparities^10–13^.

Knowledge about allelic frequencies from multiple populations is also crucial when prioritizing candidate clinical variants. For rare Mendelian disorders, the frequency in any given population cannot be higher than expected for disease incidence. Moreover, the penetrance of variants may vary across backgrounds^14,15^. For variants associated with monogenic late-onset disorders, unaffected elderly individuals serve as a proper control group to improve diagnosis accuracy. This rationale was previously explored by us using whole-exome sequencing of elderly Brazilians^16^, and by others using a European-descent whole-genome dataset of Australian elderly^17^.

Here we present the first high-coverage WGS of a Latin American census-based cohort composed of 1,171 unrelated elderly from São Paulo, Brazil’s largest metropolis, which includes immigrant descendants from different continents and individuals from various Brazilian states^18^. These individuals aged 60 or older have been comprehensively phenotyped by the longitudinal Health, Well-Being, and Aging (SABE - *Saúde, Bem-estar e Envelhecimento*) study. By carrying out WGS on this population-based cohort, we identified genomic variation absent from public databases, including single nucleotide substitutions, insertion/deletion variants (indels), chromosomal haplotypes, accurate HLA variant calls, mobile element insertions, and non-reference sequences (NRS). Additionally, we explored pathogenicity assertions in disease-related genes of clinical relevance and GWAS performance for selected phenotypes. We also created new reference imputation panels for the whole-genome and HLA alleles, which improved imputation accuracy. Lastly, we provide variants and respective allelic frequencies in a public resource, ABraOM (http://abraom.ib.usp.br).

## COHORT DESCRIPTION

SABE is a longitudinal study initiated in 2000, with a follow-up occurring every 5 years (see Supplementary Information and Supplementary Fig. 1 for details on study design). After quality control, 1,171 unrelated individuals composed the WGS dataset, with an average age of 71.86 (±7.94) years and 1.74 female to male ratio (Supplementary Table 1). Data collection involves at-home interviews and functional measurements, summarized in Supplementary Table 2.

High-coverage WGS data (average 38X) was generated and analyzed (Supplementary Fig. 2, Supplementary Table 3). Nearly 76 million single nucleotide variants (SNVs) and indels were identified with their predicted consequences, including over 22 thousand potential loss of function (pLOF) variants annotated by LOFTEE^3^ (Supplementary Table 4). After filtering out low confidence variants (Methods, Supplementary Fig. 3), we obtained a dataset of over 61 million variants, among which approximately 2 million are not described in gnomAD, dbSNP, or 1000 Genomes (Extended Data Fig. 1).

The average global ancestries for SABE are 0.726 ± 0.263 European, 0.178 ± 0.209 African, 0.067 ± 0.066 Native American, and 0.028 ± 0.162 East Asian (Fig.1A, Supplementary Table 5). There is considerable variation in individual ancestries, ranging from a single ancestry to admixture involving two or more ancestries (~75% of the cohort). Individuals with East Asian ancestry have virtually 100% of this parental component, consistent with the historical information as first generation of Japanese immigrants (Figure 1A and 1B). The proportions of ancestry differ among self-reported ethnoracial groups (p <0.001; Extended Data Fig. 2), which partially accounts for the ancestry variation (r^2^=0.63; p-value < 2.2e-16).

**Figure 1.**
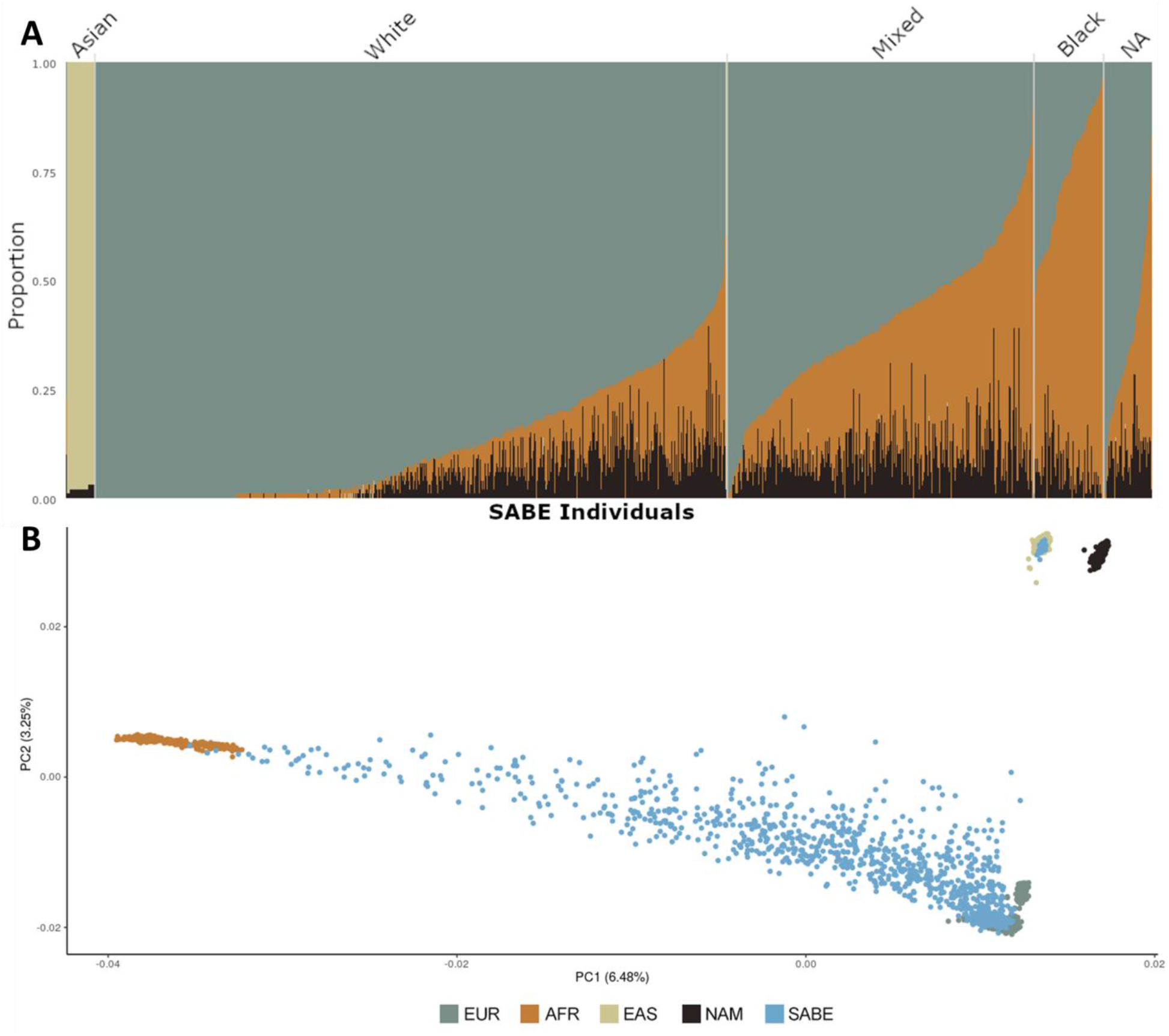
Global ancestry of SABE cohort. **A.** Individual ancestry bar plots of SABE cohort (N = 1,168) using Europeans (EUR), Africans (AFR), East Asians (EAS), and Native Americans (NAM) as parental populations and distributed by self-reported ethnoracial groups (Supplementary Figure 5). NA = Not available. **B** Principal component analysis of SABE individuals and parental populations. Analyses were performed with 372,527 SNPs (after overlapping- and LD-pruning).

## CLINICALLY RELEVANT FINDINGS

Although SABE participants are not affected by severe monogenic disorders, they might carry pathogenic variants related to recessively inherited disorders, mild phenotypes, or with incomplete penetrance. Moreover, it is known that many pathogenic assertions are misclassified^19^, and cohorts with individual genotypes and phenotypes can aid reclassification.

We analyzed ‘Pathogenic’ or ‘Likely Pathogenic’ (P/LP) ClinVar asserted variants carried by SABE individuals across 4,250 genes associated with monogenic disorders (Online Mendelian Inheritance in Man - OMIM disease genes, Supplementary Table 6) and manually curated using ACMG guidelines^20^ (Supplementary Fig. 4). In total, out of 394 variants asserted as either P/LP in genes annotated to have at least one phenotype with a dominant inheritance, curation resulted in the reclassification of pathogenicity (31% of variants), by inheritance mechanism (52%) or penetrance (13%), with only 3% of variants associated with a matching detectable phenotype (Extended Data Tab. 1).

Manual curation promotes the downgrading of pathogenic assertions when diverse ancestries are added to databases. Reclassification of variants is improved when based on standardized criteria and reports of reduced penetrance^19,21^. Moreover, variants' penetrance may differ according to different genetic or environmental backgrounds^15^, observable in well-established monogenic mutations that segregate in families^22^ that can be modified by rare variants^23^ or a polygenic profile^24^. This explains why population-specific genomic architecture reduces GWAS replication^25^ and affects distribution of polygenic risk scores^13^. Therefore, pathogenicity assertions must be interpreted based on specific population datasets. Also, regarding P/LP asserted variants in the 59 ACMG actionable genes list (Supplementary Table 11), 14 were found in 1.2% of individuals^26^ comparable to the Australian elderly cohort^17^.

Common pathogenic variants in genes associated with selected recessively inherited Mendelian disorders were manually curated using locus-specific databases and ACMG. Common and rare P/LP variants in *CFTR*, *HBB*, *GJB2*, *MEFV*, and *HFE* were accounted for incidence estimates (Supplementary Table 12). We showed that cystic fibrosis and hemoglobinopathies have similar expected incidences when compared to gnomAD. Other diseases appear more frequently in Brazilians (*GJB2*-related deafness and *MEFV* Familial Mediterranean fever) after calculating the expected offspring number of homozygotes and compound heterozygotes.

These disparities observed for *GJB2* and *MEFV* between Brazilians and global gnomAD, but similar to gnomAD Latinos and PAGE Study samples of Cubans, Puerto Ricans, and Central Americans are probably due to the Iberian, Mediterranean, and Middle Eastern contributions^27–29^ in Brazil

Finally, regarding potential loss of function variants (pLOFs) within the OMIM disease genes, we identified 3,704 non-benign variants (Supplementary Fig. 5), most absent from ClinVar with frequencies comparable to gnomAD. The few but very discrepant frequencies are mostly false positives due to calling or annotation from either dataset (Supplementary Fig. 6-7).

## MOBILE ELEMENTS INSERTIONS (MEIs)

We investigated structural variations caused by insertions of mobile elements (MEIs), which constitute a rich and underexplored source of genetic variation. MEIs here identified are insertions to the human reference genome (GRCh38) occurring in at least one out of 1,171 SABE genomes. First, we found a set of 7,490 nonredundant MEIs, including 5,971, 1,131, 375, and 13 events of *Alu*, L1, SVA (SINE-R, VNTR, and *Alu* composite), and Human Endogenous Retrovirus K (HERV), respectively (Fig. 2A, variants deposited in http://abraom.ib.usp.br). As expected^30^, *Alu*, and L1 insertions are the prevalent events (94.7%). Next, we classified these MEIs into (i) *Shared* (*i.e.*, MEIs present in two or more unrelated SABE individuals and also in individuals from gnomAD); (ii) *SABE-privative* events (present in two or more SABE genomes, but absent in other genomes from Database of Genomic Variation - DGV^31^, which include gnomAD data); and (iii) *Singletons* (present in only one SABE individual and absent from DGV. *Shared* is the most frequent class, corresponding to 5,571 (74.3%) MEIs (Fig. 2B). *SABE-privative* MEIs constitute 1,501 (20.1%) events (Fig. 2B) and comprise approximately 0.97 kbp potentially polymorphic and still unreported events in other databases. We also found 418 insertions classified as *Singletons* (5.6%; Fig. 2B), which are either somatic or lineage-specific germinative MEIs. On average, each individual carries 869 MEIs (Fig. 2C), among which the vast majority (97.0%) are *Alu* (758 events, on average) and L1 (85 events, on average), Fig. 2C.

**Figure 2.**
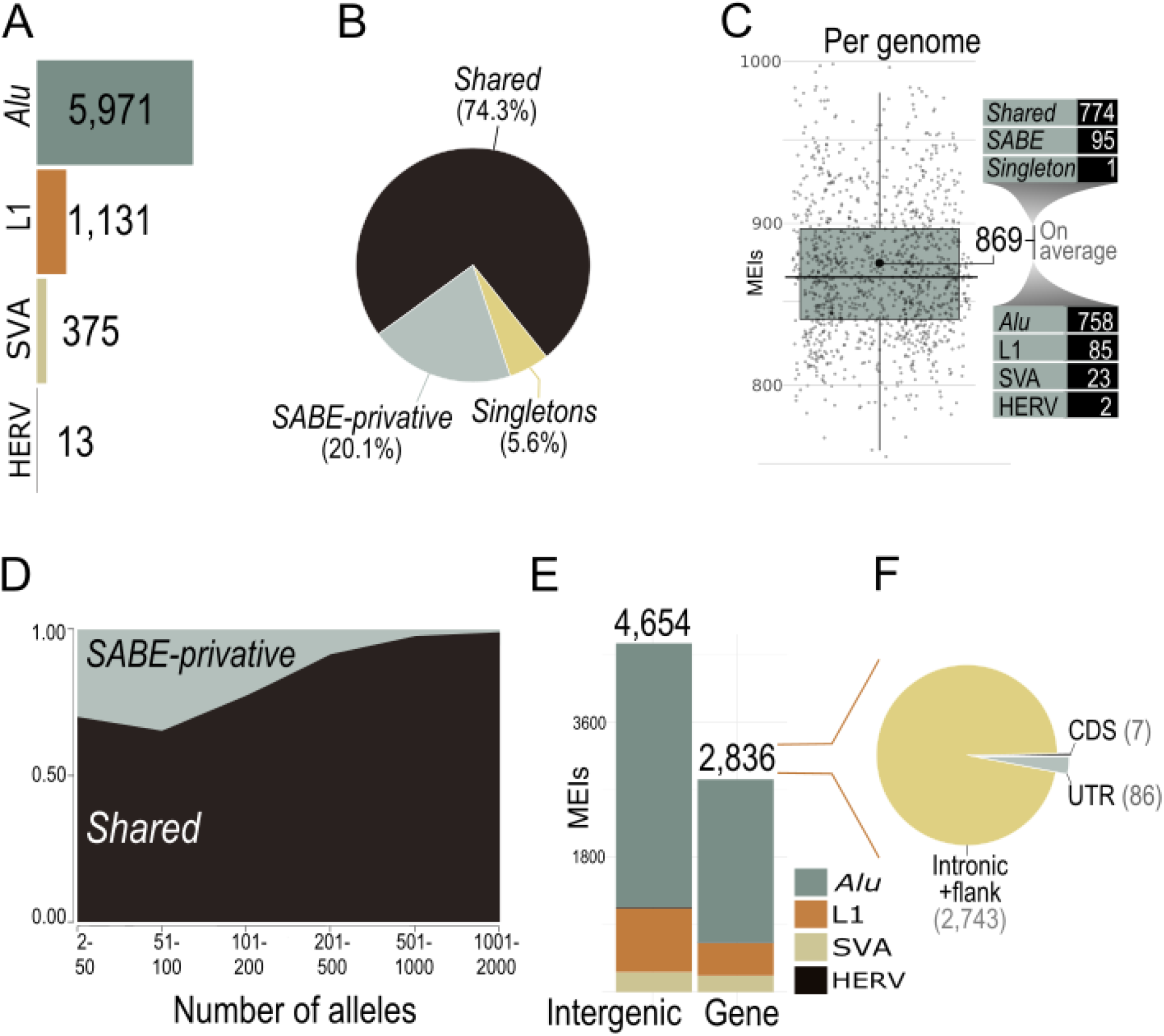
A landscape of mobile element insertions (MEIs) into SABE genomes. **A**. Total of MEIs in SABE genomes. As expected, Alu and L1 elements are predominant elements. **B.** Proportion MEIs in *Shared* (present in DGV genomes), in two or more genomes from SABE cohort (*SABE-privative*) and present in only one SABE genome (*Singletons*) **C.** Number of MEIs per individual. **D.** Distribution of allele frequencies of *Shared* and *SABE-privative* MEIs. **E.** Number of MEIs into genes and in intergenic regions. **F.** Number of MEIs in the coding region (CDS), untranslated regions (UTR), or intronic and flank (2 kbp near genes).

As expected, most MEIs per individual are *Shared* (774 (89.0%); Fig. 2C). Furthermore, individuals from our cohort carry 10.9% of events classified as *SABE-privative* (Fig. 2C), which presented a lower allele frequency in comparison to the class of common events (Fig. 2D; p-value = 1.4e-0.7; Mann-Whitney test). Even though we expected *Shared* MEIs to have a higher allele frequency, 103 (7.3%) of *SABE-privative* events presented an unexpected high allele frequency (>20%). Further validations are required to confirm if these MEIs are enriched events in our cohort (and absent in other populations) or calling artifacts.

Next, we examined the insertion profile of MEIs regarding their genomic locations. We observed: i) a positive correlation between the number of MEIs and the chromosome length (p-value = 2.74x-6; rho = 0.95; Spearman's rank correlation), Extended Data Fig. 3; ii) that L1 and *Alu* insertions are skewed to AT-rich regions, while HERVs are biased to GC-rich regions (Extended Data Fig. 4); iii) an enrichment of MEIs into intergenic regions (Fig. 2F; p-value < 0.00001; chi-square = 72.608; d.f. = 1). Out of the 2,836 MEIs within genic regions, intronic regions have significantly more (2,743) MEIs than untranslated (UTRs: 86) and protein-coding (CDS: 7) regions (p-value < 0.00001; chi-square = 62.3; d.f. = 1), indicating selection against insertions in coding (CDS) or regulatory (UTR) regions.

## NON-REFERENCE SEQUENCES (NRS)

WGS data from diverse human populations can contribute with genomic insertions that are not part of the current reference genome, so-called non-reference segments^6,32^. These mostly uncharacterized sequences contain gene exons and full genes, and may modulate susceptibility and prevalence of different diseases. We characterized these ‘missing’ segments by performing *de novo* assembly of high-quality reads that do not map to current reference.

The total lengths of NRS per individual ranged between 11.3-23.4Mbps, with an average of 15.4Mbps (Extended Data Fig. 5A). The nonredundant non-reference segments library of the SABE dataset contains 192,183 sequences (67.4Mbps), from which 428 NRS (0.43Mbps) were observed in all individuals (Extended Data Fig. 5B). Although most NRS (92.5%, totaling 56.4Mbps) are shorter than 500bps, we observed 40 contigs larger than 10 kbps, up to a maximum length of 34.5Kbps (Extended Data Fig. 5C).

Comparison with NRS from the Chinese HAN population^32^, African Pangenome^6^, Genome of the Netherlands^33^, and NCBI nonredundant database revealed that a sizable fraction of 28,264 NR-contigs (totaling 9Mbps) is unique to the SABE dataset. Simultaneously, as much as 15Mbps of NR-contigs are shared with the HAN and African Pangenome data (Fig. 3A).

**Figure 3.**
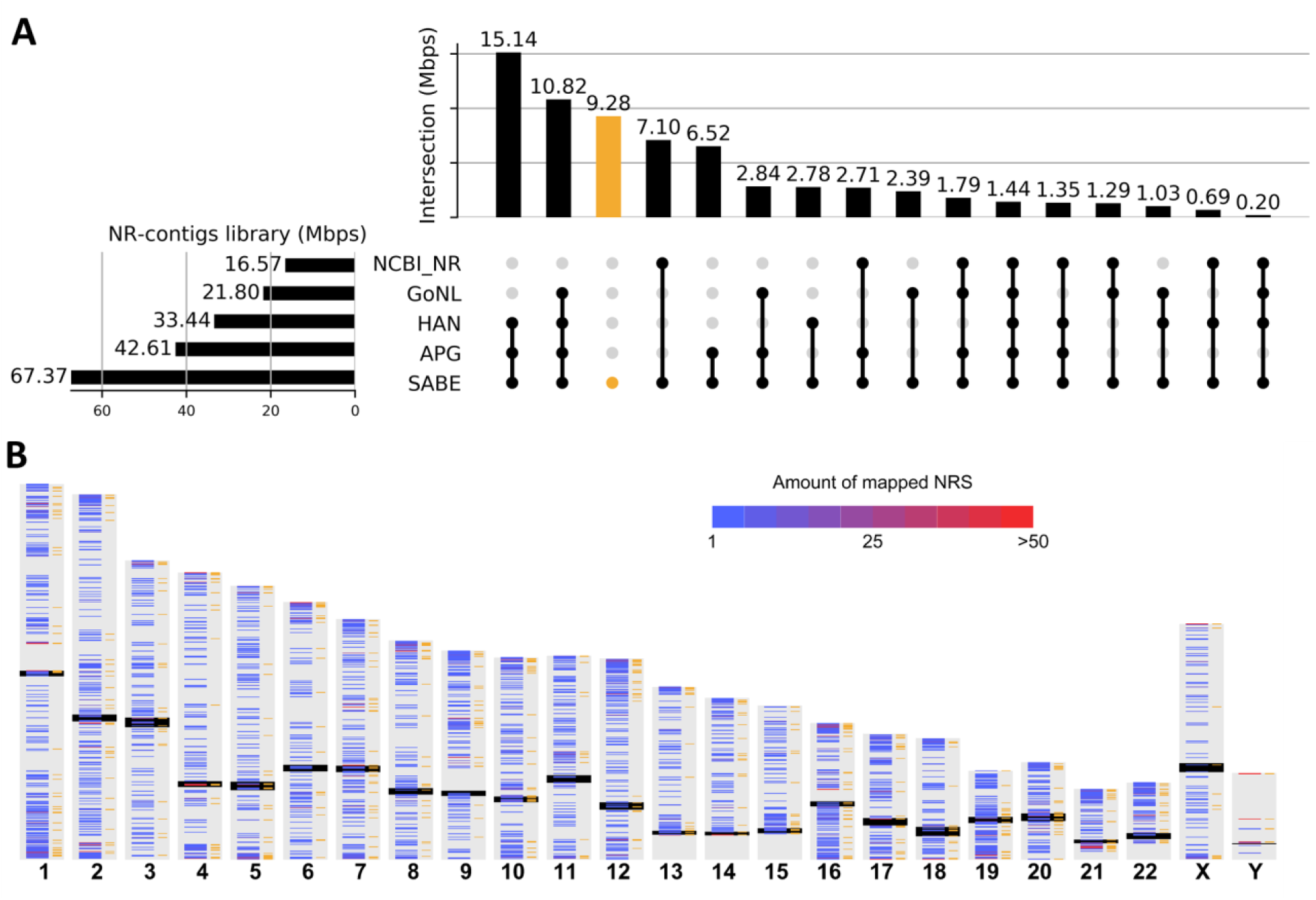
Non-reference genome sequences (NRS) in the SABE dataset. **A.** UpSet plot showing the presence of the SABE NRS in other public databases: NCBI nonredundant database (NCBI_NR), Genome of the Netherlands (GoNL), NAH Chinese (HAN) and African (APG) pan-genomes. **B.** Distribution of NRS across chromosomes. The black bars mark centromeres, bands on the left of each chromosome show density of NRS contigs, orange bands on the right side of each chromosome indicate positions of SABE-private NRS.

In total, we were able to localize 78,831 contigs (28.2Mbps) to the most recent reference assembly GRCh38, from which 12,617 of localized contigs (4.9Mbp) are unique to our dataset (Fig. 3B). The reported population frequency and genomic location of these non-reference segments will assist future functional studies that characterize their contribution to protein isoforms, gene regulation, and their potential link to human diseases.

## AN IMPROVED LATIN AMERICAN IMPUTATION PANEL

Previous studies have shown that using a reference panel composed of individuals with a similar genetic background to the target sample improves imputation accuracy, especially for rare variants^34^. We created a new imputation panel by merging SABE and the public 1000 Genomes Project Phase 3 dataset (1KGP3)^35^, hereafter called the SABE+1KGP3 reference panel. Data from chromosomes 15, 17, 20, and 22 were used to test the usefulness of the SABE+1KGP3 reference panel compared to the 1KGP3 alone. We imputed a dataset of Omni 2.5M Illumina array genotyped on 6,487 Brazilians from the EPIGEN initiative, which is composed of three different cohorts across the country (Salvador, Bambuí, and Pelotas), that vary in admixture levels and demographic histories^36^. When using the SABE+1KGP3 reference panel, we imputed the largest number of variants, ~20% of which were added exclusively by the SABE dataset (Fig. 4A). There was a gain of ~8% of high confidence imputed variants (info score > 0.8) by the SABE+1KGP3 reference panel compared to 1KGP3 alone (Fig. 4B), driven mainly by very rare variants (Fig. 4B), which also mainly contributed in improving imputation accuracy measured by r^2^ increase (Fig. 4C). The SABE+1KGP3 reference panel improved imputation independent of the target cohort and its level of admixture, suggesting that our panel can improve imputation for other Latin American populations. This improvement was also observed regardless of the chromosome tested (Supplementary Fig. 8-22; Supplementary Tables 13-17).

**Figure 4.**
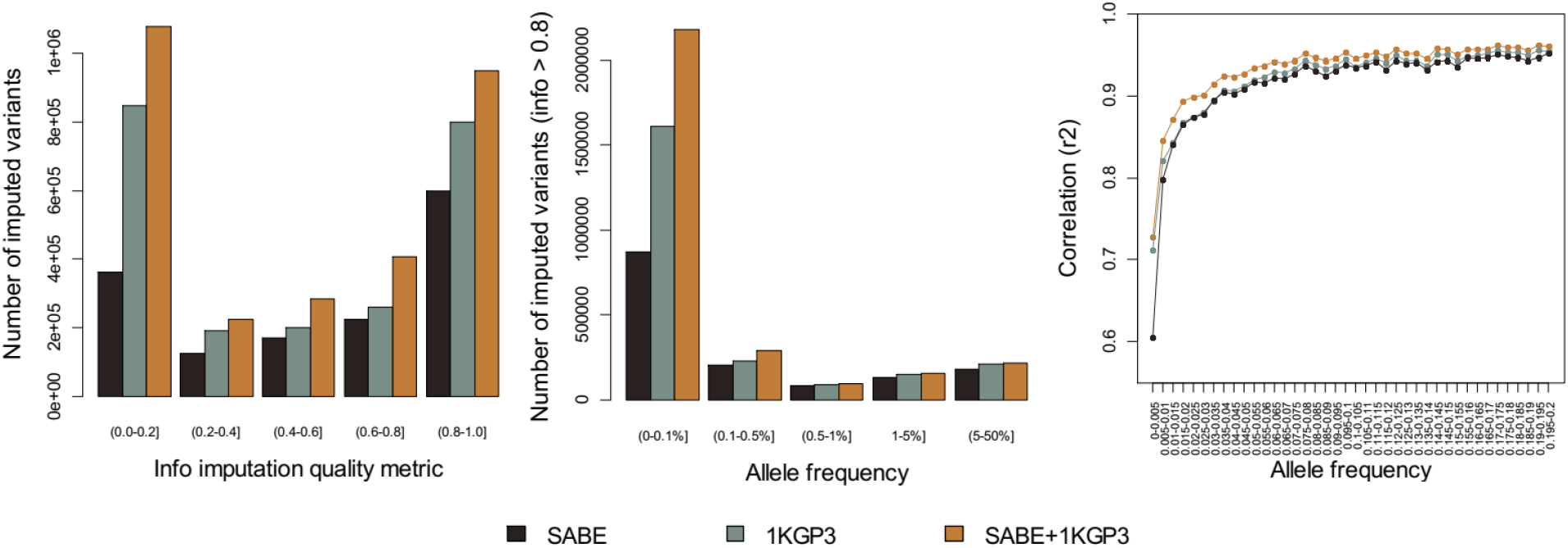
Comparison of imputation performance of SABE, 1KGP3, and SABE+1KGP3 reference panels using the Omni 2.5M array data for 6,487 Brazilians from EPIGEN as target panel (chromosome 15). **A.** The total number of imputed variants across different classes of info score quality metric. **B.** The total number of imputed variants with info score ≥ 0.8 across the allele frequency spectrum. **C.** Improvement in imputation accuracy as a function of minor allele frequency (MAF) for the target dataset after imputation (MAF from 0 to 0.2, bin sizes of 0.005). Similar results were reached for the other chromosomes tested and for each cohort (Supplementary Fig. 8-22; Supplementary Tables 13-17).

## DIVERSITY OF HLA GENES

We previously developed *hla-mapper*^37^ to optimize mappings for HLA genes, providing high-confidence genotype and haplotype calls for this unusually polymorphic region^38^, with complex structure involving duplications. We applied *hla-mapper* in the SABE dataset, detecting 2.4X more variants in the HLA class I genes than with the computational workflow for genotype calling used in the entire genome. We identified an abundance of new rare variants (Fig. 5B) and haplotypes (Extended Data Fig. 6), defining 143 novel HLA alleles, mostly rare.

**Figure 5.**
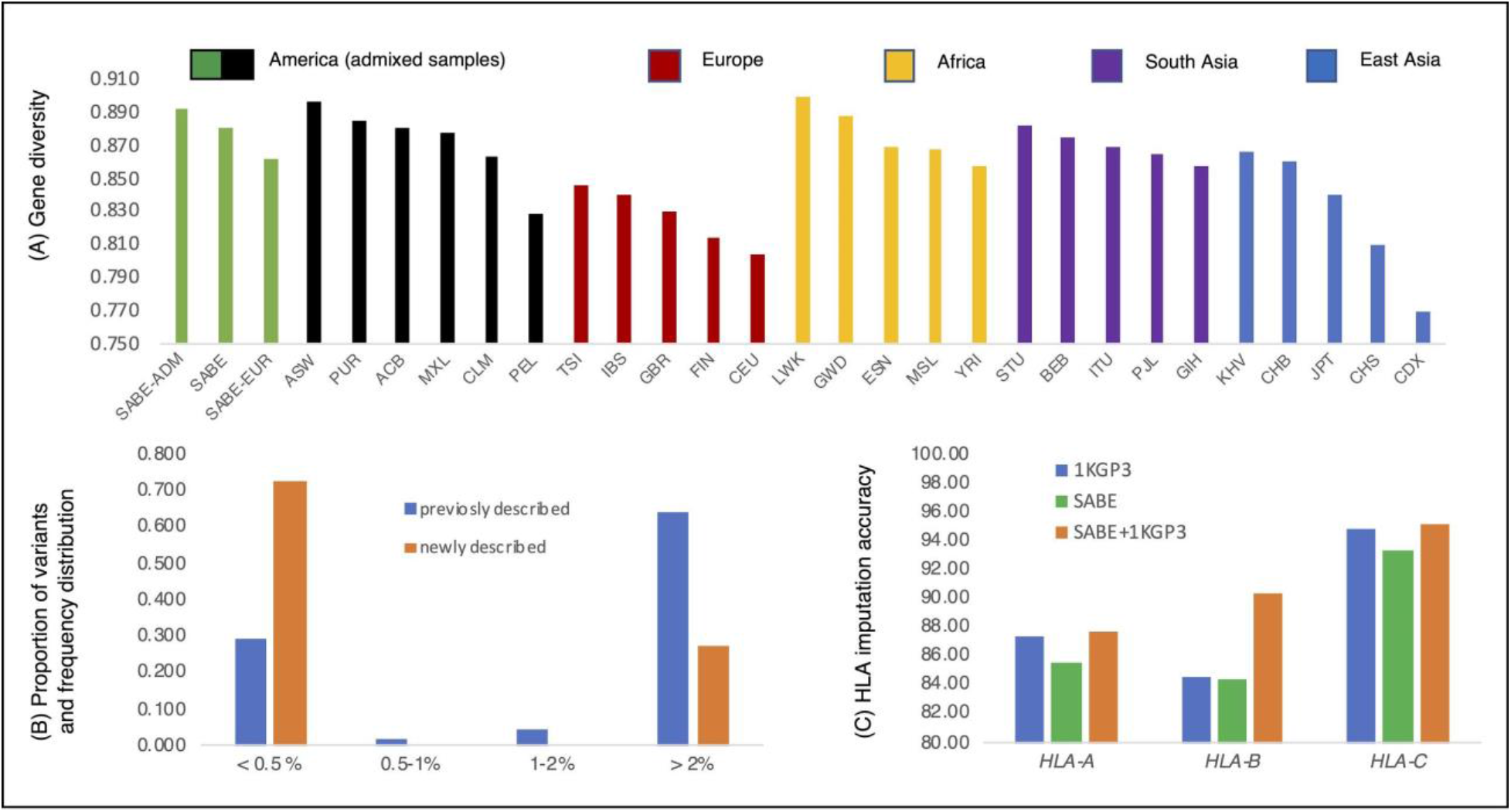
HLA polymorphism in the SABE cohort. SABE and 1KGP3 samples were processed with the same HLA workflow, as described in the Supplementary Information. **A.** Average gene diversity across SABE and the 1KGP3 populations considering haplotypes of all SNPs, i.e., the 2065 SNPs from six HLA class I genes, *HLA-A, HLA-B, HLA-C, HLA-E, HLA-F*, and *HLA-G*. SABE: all samples from SABA dataset; SABE-ADM: samples with at least 30% of both European and African global ancestry; SABE-EUR: samples with 100% European global ancestry. **B.** The proportion of SABE SNPs found at different minor allele frequency classes. **C.** HLA imputation accuracy when using the 1KGP3 (blue), SABE (green), and combining both (orange). Imputation was performed on 146 highly admixed Brazilians previously genotyped on Axiom Human Origins array and HLA genotyping by sequence-based typing methods.

While only 1% of the SABE individuals carry sequences that code for previously undescribed HLA proteins for at least one HLA class I locus, 33% have at least one new sequence comprising introns, exons, and UTRs. Moreover, 2.9% of variants detected in the HLA class I loci are novel with respect to dbSNP, concentrated in introns and regulatory sequences. The list of HLA variants and their frequencies are available in the ABraOM database (http://abraom.ib.usp.br).

To contextualize our findings, we compared polymorphism for the full sequence of *HLA-A*, *HLA-B*, *HLA-C*, *HLA-E*, *HLA-F*, and *HLA-G* in the SABE to 26 populations from the 1000 Genomes Project (1KG3P), processed through the *hla-mapper* pipeline. A highly admixed subset of SABE individuals (with at least 30% of both European and African global ancestry, n=207, SABE-ADM, Fig. 5A) presented the third-highest worldwide gene diversity, and second-highest allele richness and mean number of observed haplotypes. The subset of individuals with 100% European global ancestry (n=152, SABE-EUR, Figure 5A) had the lowest diversity among subsets we explored within SABE, although still higher than that of individual European populations from 1KGP3. These results highlight not only the contribution of non-European admixture to HLA polymorphism in Brazilians, but also the presence of European ancestries (such as Iberic and Mediterranean) that are likely to be underrepresented in major databases.

Finally, we used SABE as part of a reference panel to impute HLA alleles in a sample of 146 highly admixed Brazilians from another study^39^. As for the whole-genome imputation, the SABE+1KG3P combined reference panel provided higher accuracy than 1KGP3 panel alone (Figure 5C), particularly for *HLA-B* (an increase of 5.87%).

## GWAS

We performed genome-wide association analyses for BMI, LDL, triglycerides, positive history of cancer, cognitive decline, diabetes, frailty, and hypertension (Supplementary Table 19). We identified 14 hits (p-value ≤ 10-9) associated with cancer, BMI, LDL, and triglycerides (Supplementary Tables 20-23, Supplementary Figs. 24-27). Of those, 12 hits are within or near genes that were not previously associated with the respective phenotype^40^. Except for two hits associated with BMI, all significant alleles are rare in our cohort and very rare among Europeans (frequency < 0.005, Supplementary Tables 20-23), and two of them are also very rare among Africans. Comparing the performance of this WGS-based association with an SNP-set mimicking an array-based (Illumina Omni2.5M) GWAS (Supplementary Tables 20-23, Supplementary Figs. 24-31), we observed only two hits (p-value≤5×10-8) associated with BMI (from the total of six hits) and two hits associated with LDL (from the total of five hits).

## DISCUSSION

São Paulo is the largest city in Latin America, with over 12 million individuals, and captures the Brazilian population's main structure. Since WGS will become the standard genomic tool for research purposes and the future of precision medicine, providing a reference for admixed populations is critical. Genomic datasets such as gnomAD and TOPMed have recently included Latin American samples, but this is the first study to include more than 1,000 high-coverage WGS in any Latin American census-based cohort. Moreover, Brazil is not represented in these databases, although it is the only Latin American country colonized by Portugal and the destination of the largest contingency of individuals brought by the slave trade from the East, Central, and West regions of Africa^41^, and homeland of hundreds of Native American groups. During the 19th and 20th centuries, São Paulo was the destination of other Europeans (Italian, German, Dutch, Polish, Spanish), Middle Eastern (Syrians and Lebanese), and East Asian (Japanese) immigrants^18^.

Even though the SABE sample size is modest compared to other initiatives, we have identified over 76 million short variants (SNVs and indels), of which ~2 million are absent from major public databases. We highlight that those elderly individuals unaffected by rare genetic disorders are useful controls and support pathogenicity classification. Regarding structural variation, we found a large set of approximately 2,000 novel mobile element insertions and nearly 5Mb of genomic segments absent from human genome reference (version GRCh38). Additionally, over 140 novel HLA alleles were inferred in our sample. Whole-genome and HLA imputation were improved by the dataset when combined with 1KG3P, pointing that sample size can be, to some extent, compensated by diversity and representativeness. All results emphasize how WGS of admixed populations contribute as resourceful assets for population genomic studies and medical applications, as well as for improving the human reference genome.

## METHODS

### Samples

SABE is a census-based longitudinal study of elderly individuals that reside in the city of São Paulo, Brazil Details on sampling and study design can be found in Supplementary Information and Supplementary Fig. 1. All subjects in the genomic dataset have agreed on participating in this study on written consent forms approved by CEP/CONEP (Brazilian local and national ethical committee boards).

### Sequencing and quality control

Whole-genome sequencing was performed at Human Longevity Inc. following protocols previously described^1^. Library preparation was carried out using the TruSeq Nano DNA HT kit, and whole-genome sequencing was targeted at 30X and performed in Illumina HiSeqX sequencers using a 150 base paired-end single index read format. Reads were mapped to human reference GRCh38 using ISIS analysis software^1^. The sex of the samples was checked against proportions of read pairs concordantly mapped to the X chromosome and male-specific part of Y chromosomes (MSY) related to those mapped to autosomes. As expected, females showed around 55,000 CPM X chromosomal reads and below 200 CPM, while genomic data from males showed these values being around 27,500 CPM and above 550 CPM, respectively.

Following GATK's Best Practices for germline short variant discovery (single nucleotide substitutions and insertion/deletions) and using GATK software (3.7 release)^2^, we first generated individual GVCF (HaplotypeCaller) and then combined the GVCFs of all individuals (CombineGVCFs) to jointly call variants (GenotypeGVCFs) and perform Variant Quality Score Recalibration (VQSR-AS). Further, we used an in-house script to split the multiallelic variants into multiple lines and BCFtools^3^ to standardize variants by left alignment. Annovar^4^ and an in-house script were used to cross-reference the variants with dbSNP, 1000 Genomes Project, and gnomAD. The VEP-plugin LOFTEE (v0.3-beta, https://github.com/konradjk/loftee) was used to identify putative loss of function (pLOF) variants in at least one transcript irrespectively of confidence labeling.

We have previously developed an in-house two-step algorithm, CEGH-Filter, to evaluate the quality of called variants and genotypes^5^, by directly flagging genotypes based on the depth of coverage and allele balance using hard cutoffs. Variants are flagged based on proportions of flagged genotypes, to provide insight in site-context batch effects (Supplementary Fig. 3). All analyses involving SNVs and indels resulted from filtering out GATK VQSR-AS non-PASS variants and lower confidence flags from the in-house CEGH-Filter (Supplementary Information). A summarized table of computational steps, software, versions, packages, and datasets used throughout this article can be found in Supplementary Table 3.

Initial related analysis using KING^6^ identified 28 pairs of relatives (sibships and duos), and only one individual from the pair was selected as proband by the following order of criteria: having brain MRI, oldest age, and being male. We used PC-Relate implemented in the GENESIS software^7^ and the same dataset used for Admixture (see topic below) to confirm that no first degree relatives remained in the sample. verifyBAMID^8^ identified one sample with over 3% of contamination, leading to its exclusion. A final dataset of 1,171 unrelated participants was used in downstream analyses (Supplementary Figure 1). Samples reached a minimum mean depth of coverage of 31.3X up to 64.8X, with an average depth of coverage of 38.65X and a median of 36.6X (Supplementary Fig. 2).

### Ancestry analyses

We used ADMIXTURE v.1.3.0^9^ to perform global ancestry inference through supervised analysis (K = 4). African (N=504), European (N=503), and East Asian (N=400) samples from 1KGP3, and Native Americans (N=221) from recently published datasets^10^, were used as parental populations (Supplementary Table 5). The Native American samples were genotyped on the Illumina Omni 2.5M array; thus the genetic variants of the 1KGP3 and SABE samples (dataset of PASS (GATK) and vSR (CEGH Filter, Supplementary Fig. 3) variants with genotypes flagged by CEGH-Filter as FD or FB set as missing) were filtered to overlap with this array, totaling 1,842,125 SNPs. LD-pruning on this subset of markers was performed with PLINK v.1.9^11^, with an r^2^ threshold of 0.1 within a sliding window of 50Kb and a shift step of 10Kb, resulting in 372,527 SNPs. We used the same dataset to perform PCA analysis with SNPRelate^12^. The PCs obtained were further used for ancestry adjustment in GWAS analysis.

### Clinical analyses

To evaluate the occurrence and clinical significance of pathogenic variants in genes associated to Mendelian disorders, a comprehensive panel containing 4,250 OMIM disease genes (Supplementary Table 6) was retrieved and used for filtering SNVs and indels annotated with ClinVar pathogenic assertions (Pathogenic, Likely Pathogenic and Conflicting containing Pathogenic) and/or pLOFs identified by LOFTEE^13^. Classification of modes of inheritance was based initially by OMIM references, and upon manual curation with ClinGen (https://clinicalgenome.org/) and PanelApp (https://panelapp.genomicsengland.co.uk/). Manual curation was performed using ACMG recommendations^14^, with current literature and evaluation of the most recent phenotypes collected in SABE follow-up, when available. Summary of steps and workflows can be found in Supplementary Fig. 4-5.

### Mobile Elements Insertions

Mobile Elements Insertions (MEIs) were detected using Mobile Element Locator Tool^15^ (MELT; ver. 2.1.4). Specifically, MEIs (Alu, LINE-1, HERVs, and SVA) absent from the reference genome (GRCh38) were called with the MELT-SPLIT program and reference MEIs were genotyped using the MELT-Deletion program using the recommended standard calling procedures (https://melt.igs.umaryland.edu/manual.php). Next, additional filters were used to obtain a high-quality call and genotyping of MEIs. We filtered out i) candidates not classified as "PASS" by MELT; ii) candidates inserted in a low complexity genomic region; iii) candidates presenting more than the expected number of discordant read pairs at the insertion site. For *SABE-privative* and *singletons* events, we also applied additional filters. We selected only MEIs with MELT ASSESS score equal five, with a defined Target Site Duplication (TSD) domain and with minimal support (>2) split reads defining the insertion point. The assignment of LINE-1, *Alu*, and HERVs events to families and subfamilies was also performed using MELT. SVAs insertions were not subclassified in families.

MEIs (Alu, LINE-1, HERVs, and SVA) discovered among SABE samples were compared to MEIs present in the Database of Genomic Variation (DGV^16^), which includes Genome Aggregation Database (gnomAD) WGS samples. SABE events found in DGV were classified as *Shared* MEIs. Only the same mobile element (e.g., *Alu-Alu*, L1-L1, HERV-HERV, or SVA-SVA) in the same genomic region was considered to be the same event, considering a ± 20 bp window of positional tolerance. Different classes of mobile elements falling in the same position are considered separate events. This overlap tolerance was based on the following possibilities: if there was a single ancestral event in the parental population followed by lineage-specific rearrangements, or calling discrepancies, or if there were independent events; regarding functional consequences and context interpretation, the overlapping events could be treated similarly. Manual examinations of the MEIs coordinate differences between our and public data revealed that the differences could be the result of variation in the TSD length or alignment adjustments.

To classify the genomic locations of MEI identified in the SABE genomes into genic (CDS, UTR, Intronic+flank) or intergenic, we matched the event coordinates against the GENCODE database. GENCODE (version 32) was used to define the set of transcribed regions. Exonic (CDS and UTR) and intronic regions (including 2k bp up and downstream the transcription start/end site) were defined as genic regions; all other genomic locations were defined as intergenic. In-house scripts were used to match MEIs coordinated to these regions aforementioned. In order to investigate the GC or AT composition of mobile elements insertion region, first, we randomly selected 10,000 windows of length 100 bp from the human reference genome (GRCh38) and calculated their GC content (control). Second, we made the same for all mobile element insertion regions, discriminating by *Alu*, L1, SVA and HERV. Finally, we tested with Kolmogorov-Smirnov test (KS test) the random windows distribution (control) against distribution of mobile element insertion point.

### Non-reference nonredundant DNA segments library

Unmapped (to GRCh38) paired reads from each individual were filtered for low-quality reads (average base quality below 20) and assembled using Megahit de novo assembler^23^. Non-reference sequence contigs (NRS) from the 1,171 individuals were cross-assembled again with Megahit, and sequences longer than 200bp were retained as nonredundant DNA segments. We aligned nonredundant segments against GRCh38 (including alternative haplotypes and decoy segments), using minimap2^24^, and we filtered out sequences with an identity of 95% or higher. We checked for bacterial and viral contaminations by blasting NRS against NCBI nonredundant database^25^.

To determine the presence/absence of NRS in each individual, we aligned unmapped reads from each individual to GRCh38 extended with NRS, using bowtie2^26^. We discarded NRS for which none of the individuals showed read coverage in the range of 7.5x – 100x as potential contaminants or misassembled contigs. For coverage calculation, we considered only reads with mapping quality above 20.

Three sources of data were used for determining genomic positions of NRS. i) For contigs where the only part of it mapped to GRCh38, and the remaining portion (at least 200bp) did not, the mapping coordinate of the former was used for anchoring the non-reference part of the contig to chromosomal location. ii) Discordantly aligned read pairs (when mapped against GRCh38+NRS) in which one read is aligned to NRS and its pair mate aligned to a chromosomal location. iii) We used publicly available 10x Chromium linked-reads data^27^ from 26 Human Genome Diversity Project individuals (HGDP)^28^ and nine Human Genome Structural Variation Consortium individuals (HGSVC)^29^ to find overlap between barcodes mapped to NRS and chromosomal regions. Using bowtie2, we aligned 10X Chromium Genomes Linked Reads data to extended GRCh38+NRS reference and extracted barcodes for reads uniquely mapped to NRS. The best target location for each NRS was defined as a location with the highest cross-sample number of linked reads with matching barcodes (per 1kb window). NRS was considered as reliably localized if the best target location was discovered by at least two chromium barcodes (in the same or different individuals). Mapping positions of NRS anchored to mitochondrial DNA or decoy sequences were not reported. In cases when multiple mapping information was available, the preference was given to coordinates obtained by partial mapping or discordant paired reads giving more precise genomic coordinates.

### Whole-genome imputation

To create the SABE reference panel, we used only variants flagged as PASS (GATK) and vSR (CEGH filter), we set genotypes flagged as FD or FB (by CEGH-filter) as missing and removed variants with > 5% of missing genotypes. We used SHAPEIT2^30^ to infer the chromosome phase using the extractPIRs tool, which incorporates the phase information contained in sequencing reads, improving phasing quality, particularly at rare variants^31^. We used the public 1000 Genomes Project Phase 3 haplotypes (1KGP3), version 27022019, including phased biallelic variants for 5,248 unrelated samples, that were directly aligned against GRCh38^32^. The SABE+1KGP3 reference panel was obtained by the merge of the SABE and 1KGP3 reference panels using the IMPUTE2 program^33^

To evaluate imputation performance, we used the EPIGEN-2.5M dataset comprising 6,487 Brazilians from three population-based cohorts from Brazil genotyped on the Illumina Omni 2.5M array^34^: (i) 1,309 children from Salvador with 51% of African, 43% of European, and 6% of Native American ancestry; (ii) 1,442 elderly from Bambuí with 16% of African, 76% of European, and 8% of Native American ancestry; and (iii) 3,736 young adults from Pelotas with 14% of African, 79% of European, and 7% of Native American ancestry. We used CrossMap^35^ to convert genome coordinates from hg19 to GRCh38 assembly, and removed SNPs with more than 5% missing.

We checked the consistency of the SNP’s strand of the target and each reference panel with SHAPEIT2 using the human genome reference sequence GRCh38, and we used PLINK software^36^ to flip the strands in case of inconsistencies. We phased the target EPIGEN-2.5M data set using (1) the SABE haplotypes as phasing references, for the imputation with the SABE reference panel; (2) the 1KGP3 haplotypes as phasing references, for the imputation with the 1KGP3 reference panel; and (3) the 1KGP3 haplotypes as phasing references, for the imputation with the SABE+1KGP3 reference panel.

We used IMPUTE2 to perform the imputation for chromosomes 15, 17, 20, and 22, on chromosome chunks of 7 Mb, with an additional 250 kb of buffer on both sides (these were used for imputation inference but omitted from the results) and set the effective size parameter (Ne) to 20,000. We used IMPUTE2 info score as a metric of imputation quality, in which a value of 0 indicates that there is complete uncertainty about the imputed genotypes, and 1 indicates certainty about the genotypes.

To test imputation accuracy, we used the squared correlation (r^2^) obtained by internal cross-validation performed by IMPUTE2. To this, IMPUTE2 masks the genotypes in the target panel, one by one, imputes the masked genotypes, and then compares the original genotypes with the imputed genotypes for each masked variant.

### HLA variants and haplotypes processing

WGS reads from the SABE cohort were processed as described earlier. For the 1000 Genomes dataset, we obtained high coverage BAM files using the ASPERA protocol. We processed these BAM files using *hla-mapper* version 4^17^ (www.castelli-lab.net/apps/hla-mapper), as described elsewhere^18,19^.

We used GATK HaplotypeCaller version 4.1.7 to call genotypes in the genome confidence model (GVCF), concatenating all samples together in a VCF file using GenotypeGVCFs. We processed each HLA locus separately. For variant refinement and selection, we used the vcfx checkpl, checkad, and evidence algorithms to introduce missing alleles in genotypes with low likelihood and annotate each variant with a series of quantitative parameters^19^ (www.castelli-lab.net/apps/vcfx). Each variant that has not been approved by the vcfx evidence algorithm was evaluated manually. The hla-mapper/GATK/vcfx workflow allowed the detection of 2,257 high-quality variants considering 6 HLA class I loci, *HLA-A*, *HLA-B*, *HLA-C*, *HLA-G*, *HLA-E*, and *HLA-F*, against only 910 (40%) when using the regular workflow applied to the entire genome. We also calculated gene diversity, allele richness, and the mean number of different haplotypes across the 1000Genomes populations and SABE using a local Perl script, resampling 50 samples in 5,000 batches. This dataset was used in the analysis presented here.

For haplotype inference, we combined both physical phasing using GATK ReadBackedPhasing (RBP) and probabilistic models, as described in the supplementary material. After, we exported the phased data to complete sequences (exons+introns) and CDS sequences (only exons), comparing them with the ones described in the IPD-IMGT/HLA Database version 3.4.0^20^. Allele, genotype, and haplotype frequencies were calculated by direct counting. Please refer to the supplementary material for other details regarding the HLA workflow.

### HLA imputation

Multi-ethnic imputation models for each of the class I classical HLA genes (*HLA-A, −B* and *C*) were fitted using as reference panel: (a) SABE (1171 sample); (b) 1KGP3 (2503 samples); and (c) SABE + 1KGP3 (3674 samples). The imputation models were built on HIBAG v.1.4^21^, based on overlapping SNPs with the Axiom Human Origins array (Affymetrix), with HLA allelic resolution at the protein level (HLA - 2 fields), 100 classifiers, and other default settings (Supplementary Table 18). To assess the accuracy of the models, imputation was performed on a sample of 146 highly admixed Brazilian individuals (43% AFR, 41% EUR, and 16% NAM) previously genotyped on Axiom Human Origins array and had HLA genotyped by standard methods (see details in Nunes et al., 2016^22^). To verify the accuracy of the imputation in each locus, the number of chromosomes with the correct HLA call was quantified over the total number of imputed chromosomes. The empirical cumulative distribution (ECD) was performed to access the posterior probability distribution associated with the different reference panels (Supplementary Fig. 23)

## GWAS

We selected the following phenotypes to perform genome-wide association analyses: (i) body mass index (BMI) calculated as weight (kg) divided by squared height (meters); (ii) low-density lipoprotein (LDL, mg/dL); (iii) log(10) transformed triglycerides (mg/dL); (iv) cancer (self-reported and checked in healthcare registry); (v) cognitive decline (decline: mini-MMSE < 13, no decline: mini-MMSE >=13); (vi) diabetes (self-reported); (vii) frailty (≥1 component); and (viii) hypertension (self-reported and cross-checked with medication). Distributions are shown in Supplementary Table 19. We used plink2 (www.cog-genomics.org/plink/2.0/)^11^ to run linear regressions for quantitative traits and logistic regressions for categorical variables. We excluded variants with minor allele frequency < 0.005 for the analyses. Regressions were adjusted for age, sex, classes of income, schooling, and 10 first PCs obtained with PCrelate^37^, and we used the --covar-variance-standardize flag to standardize covariates. We only used biallelic variants, flagged as PASS (by GATK), and we set genotypes flagged as FD or FB (by CEGH-filter) as missing. The final input was obtained by removing variants with MAF > 5% with more than 5% of missing data, and variants with MAF <5% with more than 1% of missing data. We also performed the analyses for an array-filtered input, obtained by including only the variants present in the Illumina (San Diego, CA, US) Omni2.5M array. Genomic inflation for all traits was below 1.05. A significance threshold of 10^−9^ and 5×10^−8^ was used for the whole genome and array-filtered data, respectively. We used the qqman package for R^38^ to build Manhattan plots and qq plots, and we estimated the linkage disequilibrium statistics using the software Haploview^39^.

## ACKNOWLEDGMENTS

We acknowledge SABE participants for their long-term contribution and Prof. Maria Lúcia Lebrão (in memoriam) for her conceptualization and conduction of SABE. Funding was provided by FAPESP grants and fellowships (CEPID 2013/08028-1, SABE 2014/50649-6, INCT 2014//50931-3, 2013/17084-2, 2017/19223-0, 2012/24731-1, 2018/15579-8, 2015/25020-0, 2020/02413-4) and Conselho Nacional de Desenvolvimento Científico e Tecnológico (CNPq INCT 465355/2014-5). We thank the Human Longevity Inc. team that supported and conducted whole-genome sequencing. We acknowledge the staff from HUG-CELL and USP Public Health School.

## AUTHOR CONTRIBUTIONS

MSN, MOS and GLY conducted short variant analyses, managed ABraOM updates and wrote the main manuscript. GLY, JYTW and MOS performed and adapted the main bioinformatics pipeline. MSN, GLY and JRMC curated clinical variants. KN and DM performed ancestry analyses. SZ, TK and VG performed NRS analyses. DC, CES, DB and RN developed and maintained ABraOM. MOS, NMA, WCSM, and ETS developed and performed WGS imputation pipeline. NSBS, ASS, MRSP, CFBC, HSA, CTMJ, and ECC developed the HLA pipeline and conducted the HLA variant and haplotype calls. RM, JLB, FOR, TLAM, and PAFG conducted Insertion of Mobile Elements analyses. RCMN, VB, HG, ETS, BLH, MLB and MFLC provided additional datasets used in imputation. MOS and MSN performed GWAS analyses. ETS, DM, VG, PAFG and ECC wrote sections of the main manuscript. MSN, MOS, GLY and MZ conceived the study. YAOD, MRPB and MZ provided SABE cohort samples and genomic data. All authors reviewed the manuscript.

