## Supplementary Information for "Whole-genome sequencing of 1,171 elderly admixed individuals from the largest Latin American metropolis (São Paulo, Brazil)"

*Authors contributed equally

**Extended Data Tab. 1.** Counts of variants per category after manual curation of 394 pathogenic variants in genes with dominant mode of inheritance ........................................................... 13

**Extended Data Fig. 1**. **Single nucleotide variants (SNVs) and insertion/deletion (indel) counts in SABE WGS dataset.** Spanning deletions were excluded from the annotation in the current study. The remaining variants were classified based on combined flags of GATK Filter and CEGH Filter. Variants with GATK Pass flags were counted as GATK+, and variants with CEGH vSR, SR, and WK flags were counted as CEGH+, a combination of both were considered high confidence (GATK+/CEGH+), in the sharp color branch or else placed in the faded branch. Variants were classified as Novel (outer branches) if they are absent in all reference databases used (dbSNP v150, gnomAD v2.1.1 genomes, gnomAD v2.1.1 exomes, ESP6500 and 1000 Genomes), or if found in at least one database were included in inner branches. Yellow boxes represent counts of indels longer than 1bp, and blue boxes represent counts of SNVs and 1bp-indels. pLOF counts in red boxes[1]. ..... 11
**Extended Data Fig. 2. Ancestry distributions and self-reported ethno-racial groups.** Upper section: Boxplots of the proportions of genetic ancestry per self-reported ethno-racial groups. Bottom table: Counts of individuals per self-reported ethno-racial group and corresponding average ancestries; Number of individuals withing different ranges of ancestry proportions........................ 12
**Extended Data Fig. 3. Number of MEIs per chromosome length.** We observed a positive correlation between the number of MEIs and the chromosome length (p-value = 2.74x-6; rho = 0.95, spearman's rank correlation)........................................................................................ 14
**Extended Data Fig. 4. Base pair composition of mobile elements insertion point.** We randomly selected 10,000 windows of length 100 bp from the human genome version 38 and calculated their GC content. Then, we made the same for all Mobile Elements Insertion points, discriminating by Alu, L1, SVA, and HERV. Finally, we tested with Kolmogorov-Smirnov test (KS test) the random windows distribution against those of MEIs. L1 and Alu insertions are skewed to AT-rich regions, while HERVs are biased to GC-rich regions. ........................................................................... 15
**Extended Data Fig. 5. Analysis of non-reference contigs. A.** The frequency of non-reference contigs (NR-contigs) in the SABE population. There are 372 NR- contigs found in all samples in the population. **B.** Violin plot showing the distribution of the total NR-contigs length in megabase pairs (Mbps) for the individuals. **C.** Length distribution of the NR-contigs with the vertical axis representing the sum of the contig lengths. From a total of 67Mbps of NR-contigs, 56Mbps are less than 500 base pairs long. There are 40 NR-contigs longer than 10kbps........................................ 16
**Extended Data Fig. 6. HLA polymorphism in the SABE cohort. A.** The average number of different HLA haplotypes observed in 10,000 resamplings of 50 individuals, considering genes HLA-A, HLA-B, HLA-C, HLA-E, HLA-F, and HLA-G. SABE: all samples from Brazil; SABE-ADM: samples with at least 30% of European and African ancestry; SABE-EUR: samples with 100% European ancestry. **B.** Distribution of known and new HLA haplotypes in the SABE+1KGP3 combined dataset compared to the IPD-IMGT/HLA database. We detected 682 new haplotypes (53.2% of all observed haplotypes), 21% exclusively in SABE (green). The cumulative frequency of these new haplotypes is 4.43%. **C.** Frequency of individuals in SABE with a new HLA allele (33%, light blue), and where these new alleles were detected. Most of the novelty was observed for non-classical HLA class I alleles (gray). **D.** HLA variant annotation and the proportion of new variants detected in the SABE cohort according to dbSNP..................................................................... 17

**Supplementary Fig. 1**. **SABE cohorts longitudinal design and datasets deposited at ABraOM. A.** The first census-based cohort (A00) participants were enrolled in 2000, with 60 years of age and older, and followed up ever since in waves of phenotypic and biological samples recollections. A new cohort (B, C, D) was included every 5-6 years, with individuals aging 60-65 years old at enrollment. Whole-genome sequencing (WGS) was performed for most subjects (n=1200) of cohorts A, B, and C enrolled in the wave of 2010, of which 1171 are unrelated. **B.** Nearly half of SABE 2010 participants (N=609) were previously whole-exome sequenced, and this dataset of variants and allele frequencies was deposited at ABraOM. The current study refers to WGS of 1171 unrelated individuals, of which 574 were in the previously published dataset[4]........................................................................ 20
**Supplementary Figure 2. Depth of coverage of 1,171 WGS samples from SABE. A.** Distribution of the average depth of coverage per individual. **B.** Histograms of horizontal coverage per vertical coverage thresholds. ........................................................................................................ 22
**Supplementary Fig. 3. CEGH-Filter algorithm.** Steps, criteria, genotype flags, and variant flags. 23
**Supplementary Fig. 4. Filtering strategies for the identification of variants of potential clinical relevance and indication of downstream results.** Among high-confidence variants, we have identified a total of 5,142 variants within 4,250 OMIM Disease genes that were found to have pathogenic, likely pathogenic, or conflicting containing at least one pathogenic ClinVar assertions, or classified as potential loss of function (pLOF). Downstream analyses pointed that: over 10% are absent from population databases, but most are singleton or low frequency (Supplementary Table 7-8); most genes contain up to five variants (Supplementary Table 9); most genes are annotated to associate with recessive modes of inheritance (Supplementary Table 10); manual curation of variants initially classified as pathogenic or likely pathogenic in genes of dominant inheritance can be either reclassified or fall indeed in recessively inherited conditions or else are required to be in trans with a more deleterious variant (Extended Data Tab. 1); SABE cohort recapitulates variant-based incidence of recessive disorders that are more prevalent in European or African populations (Supplementary Table 12)............................................................................................................................ 29
**Supplementary Fig. 5. Filtering strategies for identification of variants of potential loss of function.** Among high-confidence variants, we have identified 5,142 variants within 4,250 OMIM Disease genes, 4,000 of which were classified as potential loss of function (pLOF). A subset of any pLOF non-Benign variants corresponds to 3,704 variants with any ClinVar non Benign assertion (640 variants) plus 3,064 variants that lack any assertions. Substitutions, 1bp-indels, and indels >1bp were analyzed for the distribution of individual loads (Supplementary Fig. 6A, 6B, and 6C) and allele frequencies compared to gnomAD (Supplementary Fig. 7A). Further filtering to remove indels >1bp, common variants and LC-flagged pLOFs yielded 1,854 variants, which individual load distributions were also analyzed (Supplementary Fig. 6D, 6E, and 6F) as well as frequency comparison to gnomAD (Supplementary Fig. 7B). ............................................................................. 32
**Supplementary Fig. 6. Distribution of individual loads of potential loss of function (pLOF) variants.** Left panels: a subset of any non-Benign pLOF (ClinVar non-Benign assertions plus variants that lack any assertions) variants. Histogram of individual loads of pLOF variants in **(A)** heterozygosity, **(B)** homozygosity, and **(C)** variants with pathogenic assertions on ClinVar (including Pathogenic, Likely Pathogenic and Conflicting containing one or more pathogenic entries). Right panels: a subset of pLOF variants that are single nucleotide substitutions or 1bp-indels below 5% SABE cohort frequency and flagged as LOFTEE HC. Histogram of individual loads of pLOF variants in **(D)** heterozygosity, **(E)** homozygosity, and **(F)** variants with pathogenic assertions on ClinVar... 33

**Supplementary Fig. 7. Comparison of allele frequencies between pLOFs found in SABE and gnomAD (v2.2), within genes of pLI>=0.7 and pLI<0.7**. **A.** Subset of non-Benign variants provided a comparison of pLOF frequencies of up to 100%. **B.** Rare single nucleotide variants flagged as HC on LOFTEE. **C.** Five examples of deviation were manually verified in gnomAD and explained by context leading to calls or annotations. ........................................................................ 34
**Supplementary Fig. 8.** Comparison of imputation performance of SABE, 1KGP3, and SABE+1KGP3 reference panels using the Omni 2.5M array data for 6,487 Brazilians from **EPIGEN for chromosome 17** as target panel. **A.** The total number of imputed variants across different classes of the info score quality metric. **B.** The total number of imputed variants with info score ≥ 0.8 across the allele frequency spectrum. **C.** Improvement in imputation accuracy as a function of MAF for the target dataset after imputation (MAF from 0 to 0.2, bin sizes of 0.005)........................................ 37
**Supplementary Fig. 9.** Comparison of imputation performance of SABE, 1KGP3, and SABE+1KGP3 reference panels using the Omni 2.5M array data for 6,487 Brazilians from **EPIGEN for chromosome 20** as target panel. **A.** The total number of imputed variants across different classes of the info score quality metric. **B.** The total number of imputed variants with info score ≥ 0.8 across the allele frequency spectrum. **C.** Improvement in imputation accuracy as a function of MAF for the target dataset after imputation (MAF from 0 to 0.2, bin sizes of 0.005)........................................ 37
**Supplementary Fig. 10.** Comparison of imputation performance of SABE, 1KGP3, and SABE+1KGP3 reference panels using the Omni 2.5M array data for 6,487 Brazilians from **EPIGEN for chromosome 22** as target panel. **A.** The total number of imputed variants across different classes of the info score quality metric. **B.** The total number of imputed variants with info score ≥ 0.8 across the allele frequency spectrum. **C.** Improvement in imputation accuracy as a function of MAF for the target dataset after imputation (MAF from 0 to 0.2, bin sizes of 0.005)........................................ 38
**Supplementary Fig. 11.** Comparison of imputation performance of SABE, 1KGP3, and SABE+1KGP3 reference panels using the Omni 2.5M array data for 6,487 Brazilians from **BAMBUÍ for chromosome 15** as target panel. **A.** The total number of imputed variants across different classes of the info score quality metric. **B.** The total number of imputed variants with info score ≥ 0.8 across the allele frequency spectrum. **C.** Improvement in imputation accuracy as a function of MAF for the target dataset after imputation (MAF from 0 to 0.2, bin sizes of 0.005)........................................ 38
**Supplementary Fig. 12**. Comparison of imputation performance of SABE, 1KGP3, and SABE+1KGP3 reference panels using the Omni 2.5M array data for 6,487 Brazilians from **BAMBUÍ for chromosome 17** as target panel. **A.** The total number of imputed variants across different classes of the info score quality metric. **B.** The total number of imputed variants with info score ≥ 0.8 across the allele frequency spectrum. **C.** Improvement in imputation accuracy as a function of MAF for the target dataset after imputation (MAF from 0 to 0.2, bin sizes of 0.005)........................................ 39
**Supplementary Fig. 13.** Comparison of imputation performance of SABE, 1KGP3, and SABE+1KGP3 reference panels using the Omni 2.5M array data for 6,487 Brazilians from **BAMBUÍ for chromosome 20** as target panel. **A.** The total number of imputed variants across different classes of the info score quality metric. **B.** The total number of imputed variants with info score ≥ 0.8 across the allele frequency spectrum. **C.** Improvement in imputation accuracy as a function of MAF for the target dataset after imputation (MAF from 0 to 0.2, bin sizes of 0.005)........................................ 39
**Supplementary Fig. 14.** Comparison of imputation performance of SABE, 1KGP3, and SABE+1KGP3 reference panels using the Omni 2.5M array data for 6,487 Brazilians from **BAMBUÍ for chromosome 22** as target panel. **A.** The total number of imputed variants across different classes of the info score quality metric. **B.** The total number of imputed variants with info score ≥ 0.8 across the allele frequency spectrum. **C.** Improvement in imputation accuracy as a function of MAF for the target dataset after imputation (MAF from 0 to 0.2, bin sizes of 0.005)........................................ 40

**Supplementary Fig. 15.** Comparison of imputation performance of SABE, 1KGP3, and SABE+1KGP3 reference panels using the Omni 2.5M array data for 6,487 Brazilians from **SALVADOR for chromosome 15** as target panel. **A.** The total number of imputed variants across different classes of the info score quality metric. **B.** The total number of imputed variants with info score ≥ 0.8 across the allele frequency spectrum. **C.** Improvement in imputation accuracy as a function of MAF for the target dataset after imputation (MAF from 0 to 0.2, bin sizes of 0.005). .... 40
**Supplementary Fig. 16**. Comparison of imputation performance of SABE, 1KGP3, and SABE+1KGP3 reference panels using the Omni 2.5M array data for 6,487 Brazilians from **SALVADOR for chromosome 17** as target panel. **A.** The total number of imputed variants across different classes of the info score quality metric. **B.** The total number of imputed variants with info score ≥ 0.8 across the allele frequency spectrum. **C.** Improvement in imputation accuracy as a function of MAF for the target dataset after imputation (MAF from 0 to 0.2, bin sizes of 0.005). .... 41
**Supplementary Fig. 17.** Comparison of imputation performance of SABE, 1KGP3, and SABE+1KGP3 reference panels using the Omni 2.5M array data for 6,487 Brazilians from **SALVADOR for chromosome 20** as target panel. **A.** The total number of imputed variants across different classes of the info score quality metric. **B.** The total number of imputed variants with info score ≥ 0.8 across the allele frequency spectrum. **C.** Improvement in imputation accuracy as a function of MAF for the target dataset after imputation (MAF from 0 to 0.2, bin sizes of 0.005). .... 41
**Supplementary Fig. 18**. Comparison of imputation performance of SABE, 1KGP3, and SABE+1KGP3 reference panels using the Omni 2.5M array data for 6,487 Brazilians from **SALVADOR for chromosome 22** as target panel. **A.** The total number of imputed variants across different classes of the info score quality metric. **B.** The total number of imputed variants with info score ≥ 0.8 across the allele frequency spectrum. **C.** Improvement in imputation accuracy as a function of MAF for the target dataset after imputation (MAF from 0 to 0.2, bin sizes of 0.005). .... 42
**Supplementary Fig. 19**. Comparison of imputation performance of SABE, 1KGP3, and SABE+1KGP3 reference panels using the Omni 2.5M array data for 6,487 Brazilians from **PELOTAS for chromosome 15** as target panel. **A.** The total number of imputed variants across different classes of the info score quality metric. **B.** The total number of imputed variants with info score ≥ 0.8 across the allele frequency spectrum. **C.** Improvement in imputation accuracy as a function of MAF for the target dataset after imputation (MAF from 0 to 0.2, bin sizes of 0.005). .... 42
**Supplementary Fig. 20**. Comparison of imputation performance of SABE, 1KGP3, and SABE+1KGP3 reference panels using the Omni 2.5M array data for 6,487 Brazilians from **PELOTAS for chromosome 17** as target panel. **A.** The total number of imputed variants across different classes of the info score quality metric. **B.** The total number of imputed variants with info score ≥ 0.8 across the allele frequency spectrum. **C.** Improvement in imputation accuracy as a function of MAF for the target dataset after imputation (MAF from 0 to 0.2, bin sizes of 0.005). .... 43
**Supplementary Fig. 21.** Comparison of imputation performance of SABE, 1KGP3, and SABE+1KGP3 reference panels using the Omni 2.5M array data for 6,487 Brazilians from **PELOTAS for chromosome 20** as target panel. **A.** The total number of imputed variants across different classes of the info score quality metric. **B.** The total number of imputed variants with info score ≥ 0.8 across the allele frequency spectrum. **C.** Improvement in imputation accuracy as a function of MAF for the target dataset after imputation (MAF from 0 to 0.2, bin sizes of 0.005). .... 43
**Supplementary Fig. 22.** Comparison of imputation performance of SABE, 1KGP3, and SABE+1KGP3 reference panels using the Omni 2.5M array data for 6,487 Brazilians from **PELOTAS for chromosome 22** as target panel. **A.** The total number of imputed variants across different classes of the info score quality metric. **B.** The total number of imputed variants with info score ≥ 0.8 across the allele frequency spectrum. **C.** Improvement in imputation accuracy as a function of MAF for the target dataset after imputation (MAF from 0 to 0.2, bin sizes of 0.005). .... 44

### EXTENDED DATA

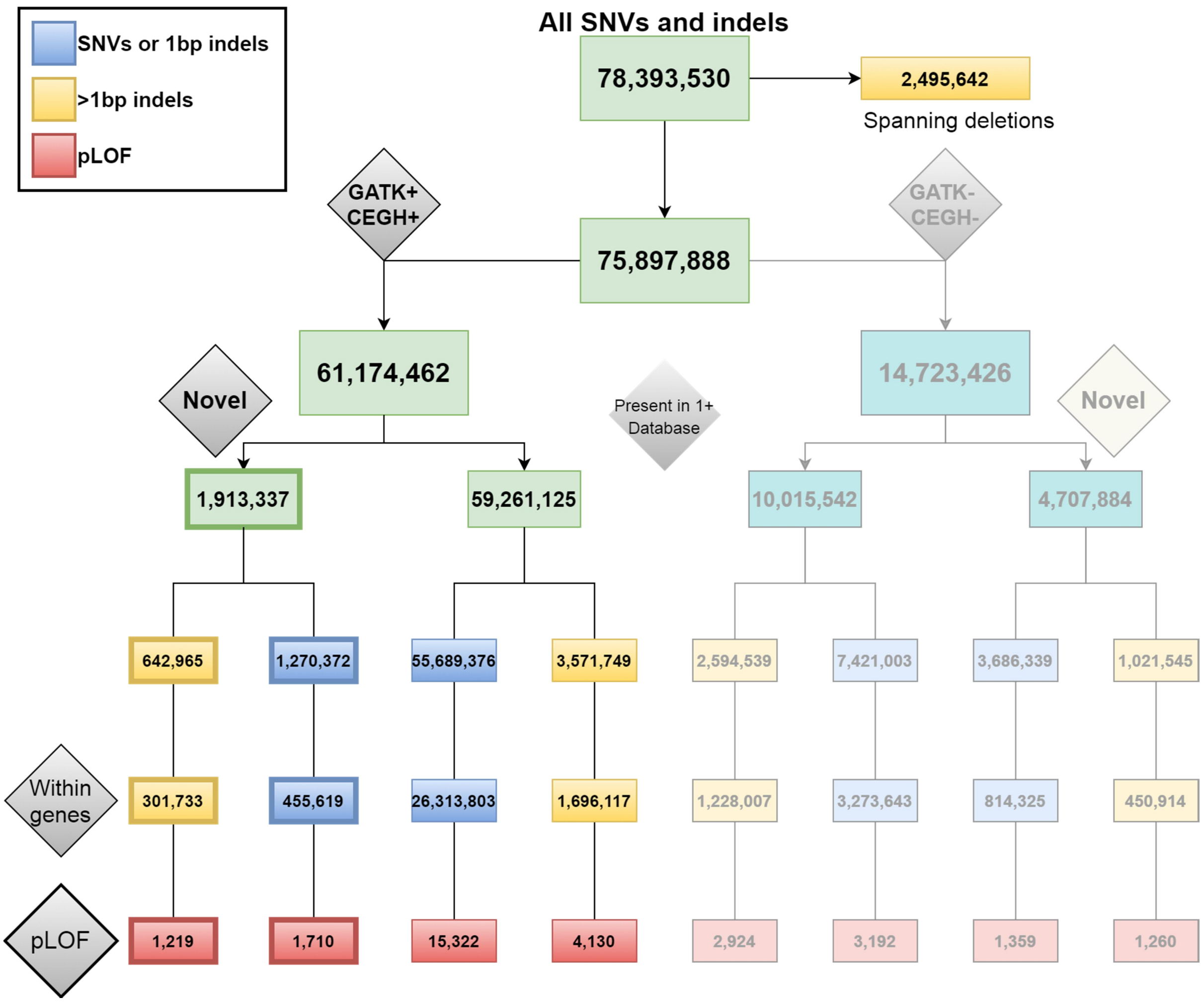

**Extended Data Fig. 1. Single nucleotide variants (SNVs) and insertion/deletion (indel) counts in SABE WGS dataset.** Spanning deletions were excluded from the annotation in the current study. The remaining variants were classified based on combined flags of GATK Filter and CEGH Filter. Variants with GATK Pass flags were counted as GATK+, and variants with CEGH vSR, SR, and WK flags were counted as CEGH+, a combination of both were considered high confidence (GATK+/CEGH+), in the sharp color branch or else placed in the faded branch. Variants were classified as Novel (outer branches) if they are absent in all reference databases used (dbSNP v150, gnomAD v2.1.1 genomes, gnomAD v2.1.1 exomes, ESP6500 and 1000 Genomes), or if found in at least one database were included in inner branches. Yellow boxes represent counts of indels longer than 1bp, and blue boxes represent counts of SNVs and 1bp-indels. pLOF counts in red boxes[1].

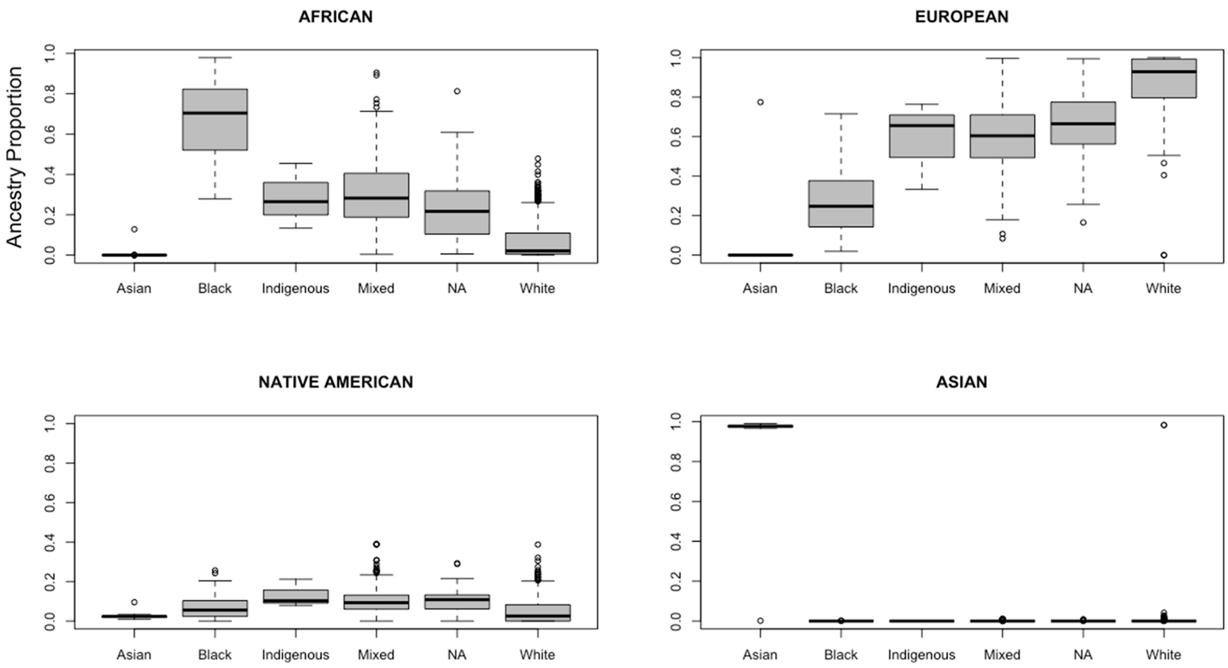

| | | Average ancestry | | | |
|---|---|---|---|---|---|
| **Self-reported ethno-racial group** | **N** | **AFR** | **EUR** | **NAM** | **EAS** |
| Asian | 32 | 0.004 | 0.024 | 0.025 | 0.947 |
| Black | 75 | 0.670 | 0.261 | 0.068 | 0.000 |
| Indigenous | 3 | 0.285 | 0.584 | 0.131 | 0.000 |
| Mixed | 330 | 0.298 | 0.600 | 0.102 | 0.000 |
| NA = No answer/Not available | 51 | 0.247 | 0.649 | 0.103 | 0.001 |
| White | 680 | 0.069 | 0.879 | 0.049 | 0.003 |
| | | **Number of individuals** | | | |
| | **Proportion** | **AFR** | **EUR** | **NAM** | **EAS** |
| Individuals with ancestry proportion above or equal to | 0.3 | 262 | 1072 | 8 | 33 |
| | 0.5 | 96 | 964 | 0 | 33 |
| | 0.7 | 46 | 711 | 0 | 33 |
| | 0.9 | 8 | 394 | 0 | 33 |

**Extended Data Fig. 2. Ancestry distributions and self-reported ethno-racial groups.** Upper section: Boxplots of the proportions of genetic ancestry per self-reported ethno-racial groups. Bottom table: Counts of individuals per self-reported ethno-racial group and corresponding average ancestries; Number of individuals withing different ranges of ancestry proportions.

**Extended Data Tab. 1.** Counts of variants per category after manual curation of 394 pathogenic variants in genes with dominant mode of inheritance

| Category after curation | Counts | % |
|---|---|---|
| Compatible finding | 2 | 0.51 |
| Subclinical/Mild phenotypes | 6 | 1.52 |
| Somatic mosaicism | 1 | 0.25 |
| Somatic mosaicism/Subclinical/Mild phenotype | 2 | 0.51 |
| Conditional (PGx) | 2 | 0.51 |
| Incomplete penetrance/Subclinical/Mild phenotypes | 10 | 2.54 |
| Recessive allele/Incomplete penetrance/Subclinical | 1 | 0.25 |
| Incomplete penetrance | 39 | 9.90 |
| Recessive allele/Incomplete penetrance | 2 | 0.51 |
| Recessive allele | 60 | 15.23 |
| Recessive gene | 146 | 37.06 |
| Reclassified | 123 | 31.22 |
| **Total** | 394 | 100.00 |

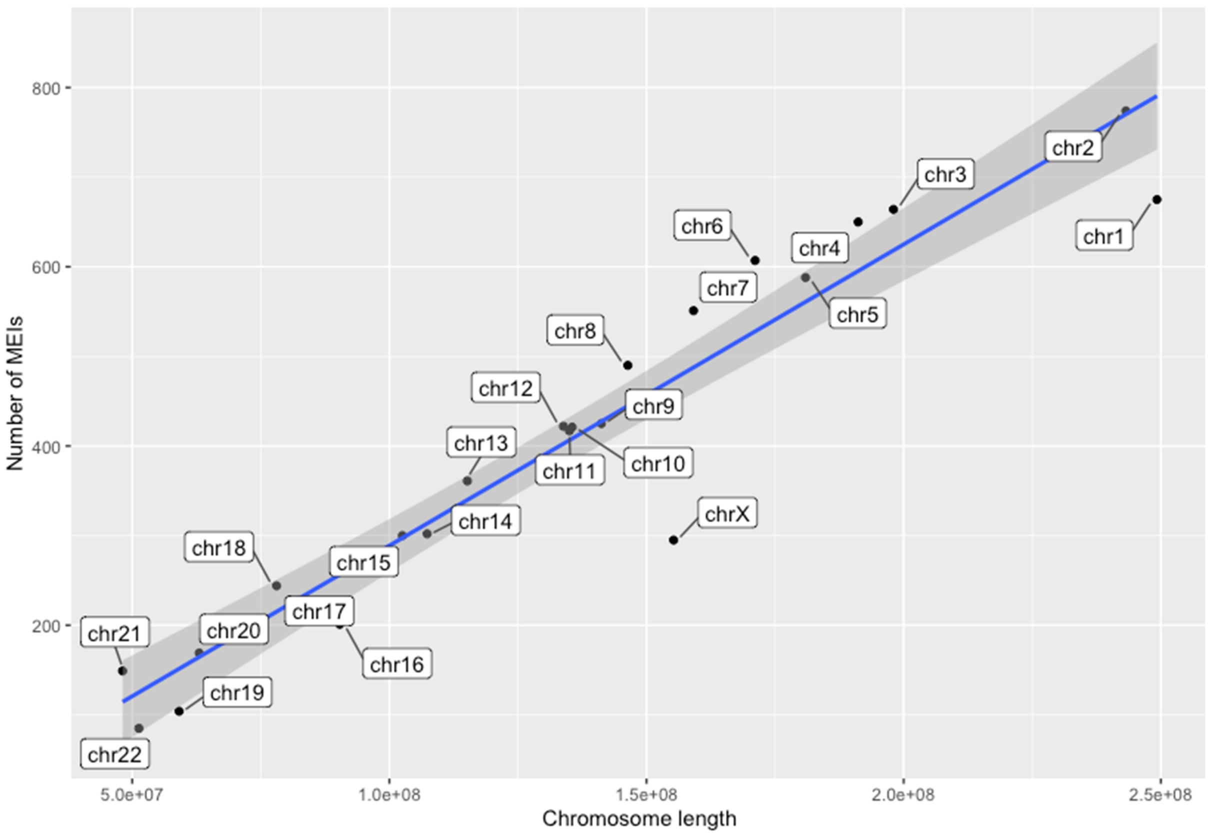

**Extended Data Fig. 3. Number of MEIs per chromosome length.** We observed a positive correlation between the number of MEIs and the chromosome length (p-value = 2.74x-6; rho = 0.95, spearman's rank correlation).

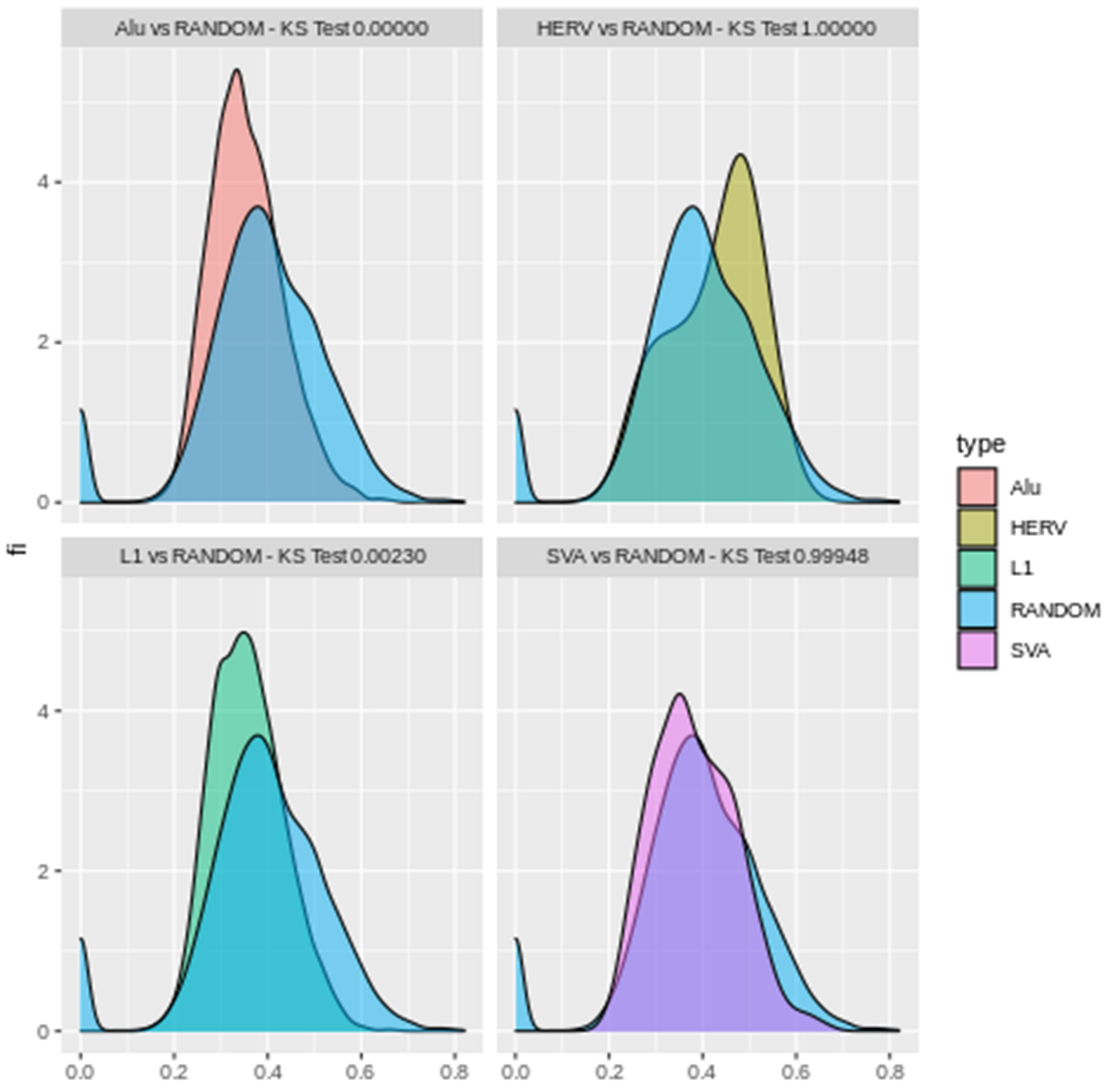

**Extended Data Fig. 4. Base pair composition of mobile elements insertion point.** We randomly selected 10,000 windows of length 100 bp from the human genome version 38 and calculated their GC content. Then, we made the same for all Mobile Elements Insertion points, discriminating by *Alu*, L1, SVA, and HERV. Finally, we tested with Kolmogorov-Smirnov test (KS test) the random windows distribution against those of MEIs. L1 and *Alu* insertions are skewed to AT-rich regions, while HERVs are biased to GC-rich regions.

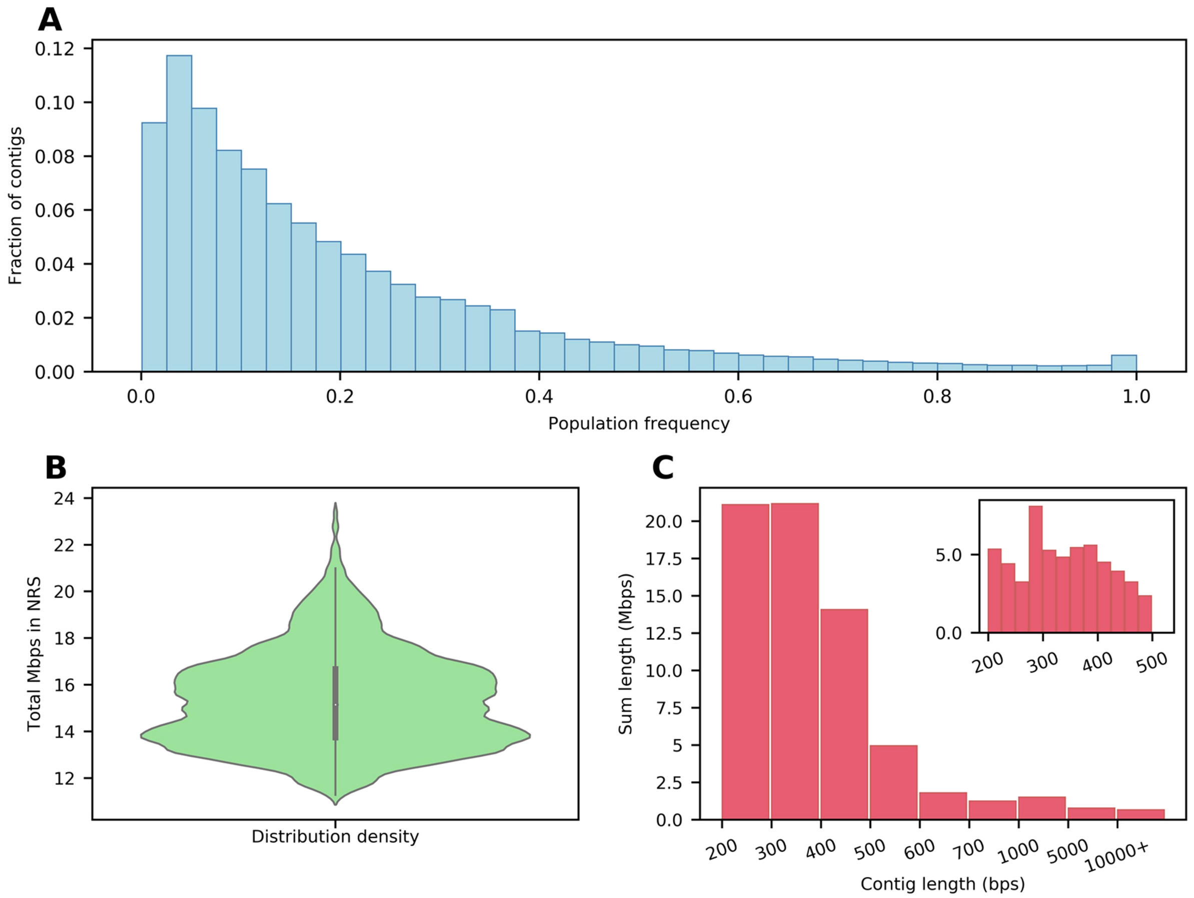

**Extended Data Fig. 5. Analysis of non-reference contigs. A.** The frequency of non-reference contigs (NR-contigs) in the SABE population. There are 372 NR- contigs found in all samples in the population. **B.** Violin plot showing the distribution of the total NR-contigs length in megabase pairs (Mbps) for the individuals. **C.** Length distribution of the NR-contigs with the vertical axis representing the sum of the contig lengths. From a total of 67Mbps of NR-contigs, 56Mbps are less than 500 base pairs long. There are 40 NR-contigs longer than 10kbps.

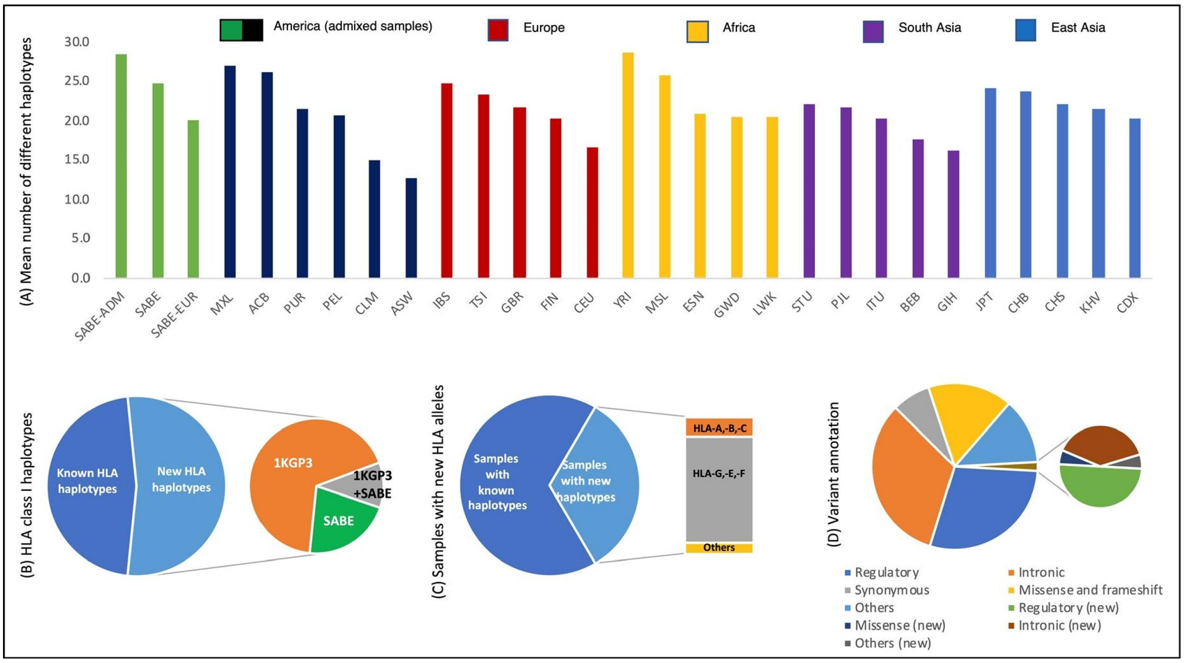

**Extended Data Fig. 6. HLA polymorphism in the SABE cohort. A.** The average number of different HLA haplotypes observed in 10,000 resamplings of 50 individuals, considering genes *HLA-A, HLA-B, HLA-C, HLA-E, HLA-F*, and *HLA-G*. SABE: all samples from Brazil; SABE-ADM: samples with at least 30% of European and African ancestry; SABE-EUR: samples with 100% European ancestry. **B.** Distribution of known and new HLA haplotypes in the SABE+1KGP3 combined dataset compared to the IPD-IMGT/HLA database. We detected 682 new haplotypes (53.2% of all observed haplotypes), 21% exclusively in SABE (green). The cumulative frequency of these new haplotypes is 4.43%. **C.** Frequency of individuals in SABE with a new HLA allele (33%, light blue), and where these new alleles were detected. Most of the novelty was observed for non-classical HLA class I alleles (gray). **D.** HLA variant annotation and the proportion of new variants detected in the SABE cohort according to dbSNP.

### SUPPLEMENTARY INFORMATION

### 1. Cohort description

The Health, Well-being, and Aging (SABE) Study is a large effort to investigate health-related conditions of the elderly in Latin America and the Caribbean, initiated in 2000 with a follow-up design and at the time coordinated by the Pan American Health Organization. The Brazilian branch is based on the Public Health School at the University of São Paulo and enrolled elderly from the city of São Paulo, the largest in the Southern hemisphere. Subjects were invited based on probabilistic sampling from the census stratified from 60 years of age and older at the time of collection, with an oversample at the initial cohort of individuals with 75 and older. Every five years, recollection was performed with the inclusion of new cohorts (B, C, D) to reintroduce elderly subjects aging 60-64 (Supplementary Figure 1A)[2]. Supplementary Table 1 presents the age and sex distribution of SABE cohorts.

**Supplementary Table 1**. Age and sex distribution of SABE cohorts

| Entry SABE cohort | Wave 2010 (n) | Unrelated individuals with WGS (n) | Age at collection (years ± s.d.) | Females (%) | Males (%) |
|---|---|---|---|---|---|
| A | 746 | 640 | 78.89±6.82 | 416 (65) | 224 (35) |
| B | 240 | 220 | 65.52±1.29 | 135 (61.4) | 85 (38.6) |
| C | 349 | 311 | 61.86±1.36 | 193 (62.1) | 118 (37.9) |
| Total | 1335 | 1171 | 71.86±7.94 | 744 (63.5) | 427 (36.5) |

SABE participants were asked for specific consent on taking part in genomic studies from the year 2010 and beyond. All subjects in the genomic dataset have agreed on participating in this study on written consent forms approved by CEP/CONEP (Brazilian local and national ethical committee boards) under the following protocols: COEP FSP USP OF.COEP/23/10, CONEP 2044/2014, CEP HIAE 1263-10.

Our group in the HUG-CELL center was responsible for creating SABE DNA collection and sequencing its subjects to evaluate their genomes' features. In 2017, 609 individuals of three SABE cohorts (A, B, and C) were whole-exome sequenced, and variants and respective allelic frequencies deposited in ABraOM (http://abraom.ib.usp.br), a resource that has been widely used by the scientific community and by molecular diagnosis laboratories as controls (Supplementary Figure 1B). Later, whole-genome sequencing of near all samples from the 2010 wave was performed (Supplementary Figure 1, Supplementary Table 2).

From a total of 1,335 SABE participants enrolled in 2010, samples from 1,200 met quality criteria and were submitted to whole-genome sequencing at Human Longevity Inc. using the protocols previously described [3]. Relatedness was assessed by KING, and when identifying siblings and duos, one individual was maintained. The final number of unrelated individuals was 1,171 (Supplementary Figure 1, Supplementary Table 2).

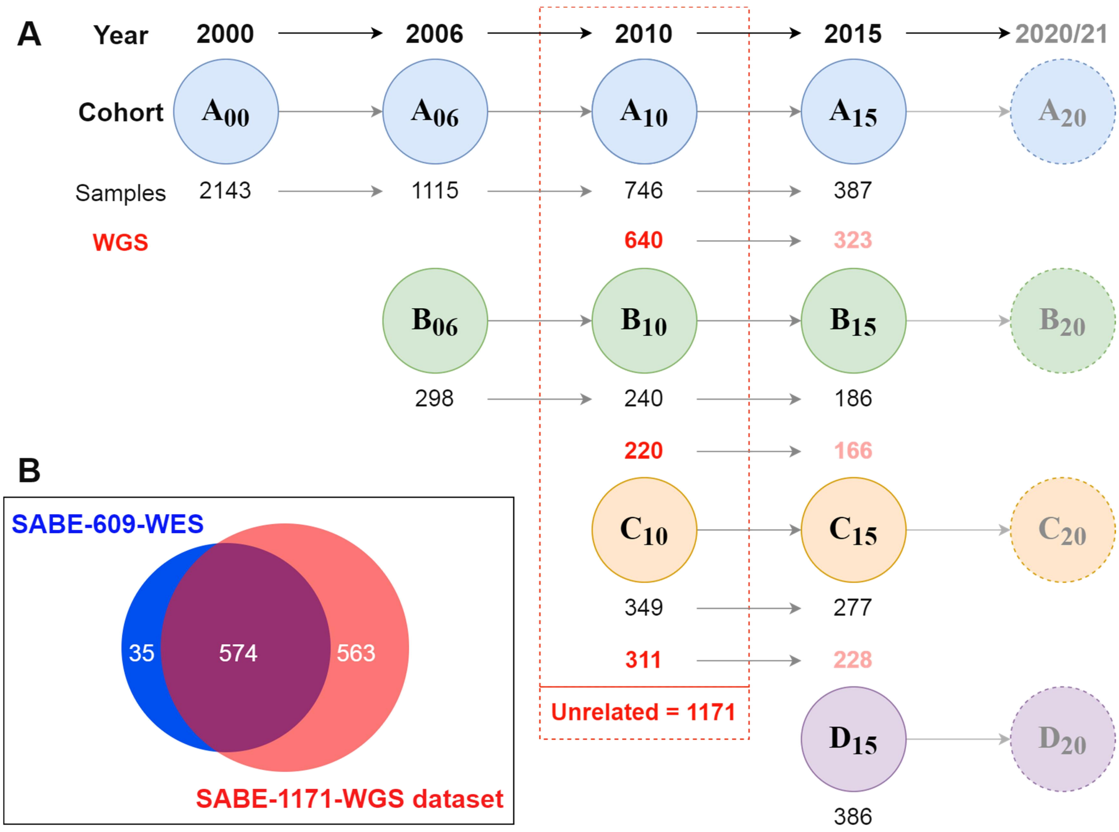

**Supplementary Fig. 1**. **SABE cohorts longitudinal design and datasets deposited at ABraOM. A.** The first census-based cohort (A00) participants were enrolled in 2000, with 60 years of age and older, and followed up ever since in waves of phenotypic and biological samples recollections. A new cohort (B, C, D) was included every 5-6 years, with individuals aging 60-65 years old at enrollment. Whole-genome sequencing (WGS) was performed for most subjects (n=1200) of cohorts A, B, and C enrolled in the wave of 2010, of which 1171 are unrelated. **B.** Nearly half of SABE 2010 participants (N=609) were previously whole-exome sequenced, and this dataset of variants and allele frequencies was deposited at ABraOM. The current study refers to WGS of 1171 unrelated individuals, of which 574 were in the previously published dataset[4].

Since baseline, several health-related phenotypes were collected at their households, including the self-reported history of prevalent disorders, medications, measured anthropometric values, and functional tests relevant to elderly individuals. Questionnaires are comprehensive and were expanded and optimized every step, with about 3,500 variables, most nested within specific interrogations (treatment details on disorders). A total of 496 individuals were successfully recruited to perform additional data collections, including magnetic resonance (3T) of the brain at Albert Einstein Hospital (Supplementary Table 2).

**Supplementary Table 2**. List of SABE study collected phenotypes per cohort-year

| Cohorts | Wave of collection | Measurements |
|---|---|---|
| SABE Cohort A | Baseline (2000-01) | Questionnaire;<br>Anthropometry: weight, height, waist circumference and hips;<br>Balance, mobility and flexibility. Cognitive test: "Mini" MMSE. |
| SABE Cohorts A+B | Follow up (2006) | Additional to above:<br>MMSE;<br>Blood pressure;<br>Blood glucose. |
| SABE Cohorts A+B+C | Follow up (2010-12) | Additional to above:<br>Wide range of haematological/biochemical blood tests;<br>Serum frozen at -80ºC;<br>HIV screening;<br>Urinalysis (uri-color check);<br>Immune response;<br>Accelerometer (trace movement). |
| SABE Cohorts A+B+C | Genetics + MRI + Functional (2010-14) | DNA extraction of all collected in 2010-12;<br>Whole-genome sequencing for 1,171 subjects of SABE cohorts A+B+C<br>496 individuals were recruited to Albert Einstein Hospital to perform: Brain MRI of n~452 (up to 5 acquisitions);<br>Pin pegboard of n~480;<br>Hand-grip strength n~480;<br>Ediburgh handedness inventory n~488;<br>Cognitive tests: 3MS and MMSE n~494; |

### 2. CEGH-Filter and variant analyses

We performed a standard pipeline cited in the main Methods. All software and versions can be found in Supplementary Table 3.

**Supplementary Table 3.** List of tools and versions applied in the bioinformatics pipeline.
Large table displayed only on Supplementary Tables worksheet.

In the final SABE dataset (1,171), WGS depth of coverage was assessed by GATK-DepthOfCoverage with a mapping quality threshold of 10 or greater[5]. Individual averages ranged from 31.3X to 64.8X, with a mean of individual averages of 38.65X and a median of 36.64X (Supplementary Fig. 2A). Horizontal coverage per vertical coverage thresholds yielded the complete dataset of 1,171 individual samples having 60% of bases with >30X and 91% of bases with >10X. A total of 1,098 individuals (93.7% of the sample) reach 70% of bases with >30X. (Supplementary Fig. 2B).

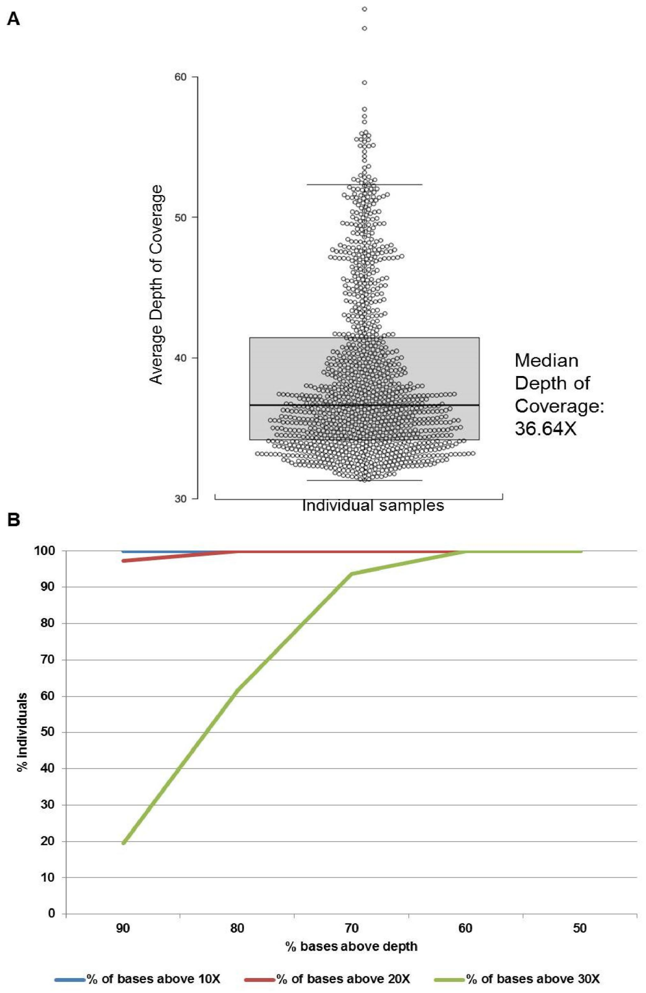

**Supplementary Figure 2. Depth of coverage of 1,171 WGS samples from SABE. A.** Distribution of the average depth of coverage per individual. **B.** Histograms of horizontal coverage per vertical coverage thresholds.

An in-house algorithm asserted genotype and variant qualities in addition to GATK flagging. CEGH-Filter (Supplementary Fig. 3) is a genotype walker algorithm that directly flags genotypes based on-site hard cutoff depth of coverage (DP>=10) and allele balance on a posteriori genotype calls (genotypes called heterozygous allelic proportion between inclusive 0.3 and 0.7; homozygous inclusive 0.1). After flagging all genotypes, each variant is flagged based on proportions of 'pass' genotypes carrying alternate alleles (heterozygotes 0/1 or alternative homozygotes 1/1) considering all genotypes at the site. Hard cutoffs on well-genotyped proportions of 90%, 50%, and one genotype to 10% will flag variants with 'Very Strong - vSR', 'Strong - SR' or 'Weak - WK' assertions, respectively. If no alternative allele carrying genotypes survive flagging, the variant is flagged either with 'Filtered out due to depth - FDP' or 'Filtered out due to allele balance - FAB' with corresponding proportions of each observation pending on 50% (FDP inclusive). If at least one genotype survives, but the quality of genotypes at the site does not fit quality criteria (cutting at 1,000 alleles or 500 genotypes), a 'Low cohort call' flag is added to vSR, SR, or WK. Allele frequencies are calculated before and after genotype flagging.

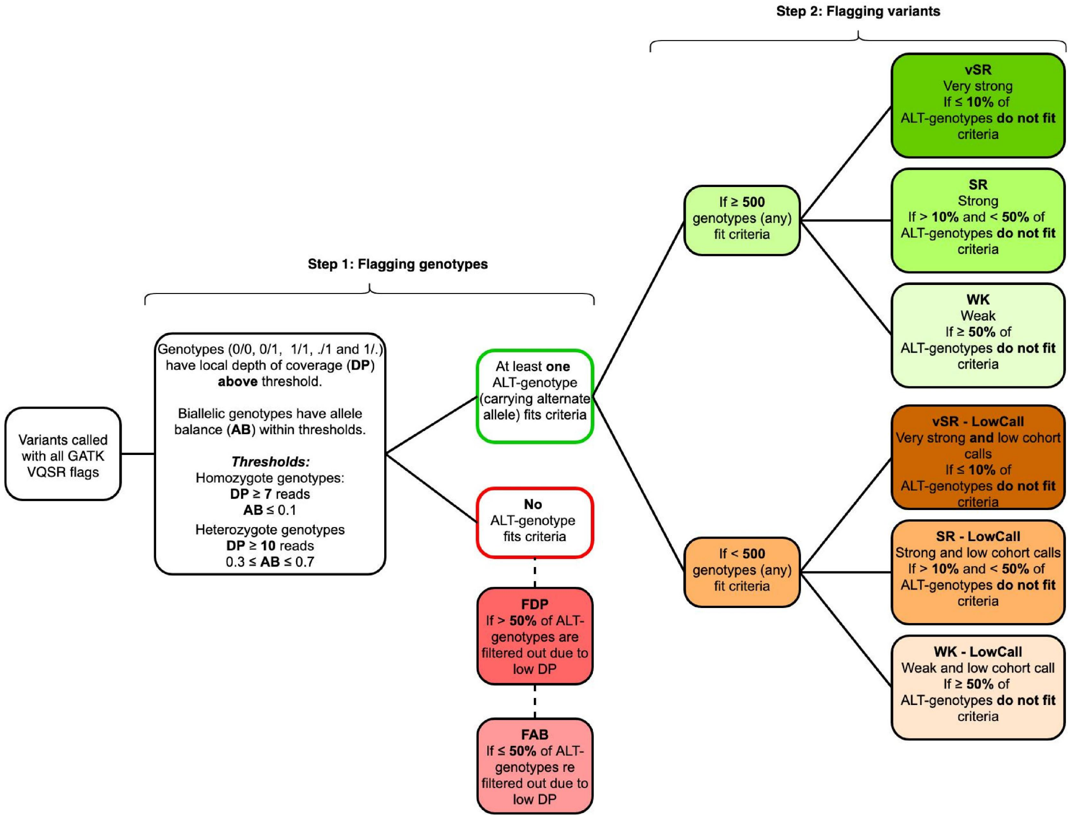

**Supplementary Fig. 3. CEGH-Filter algorithm.** Steps, criteria, genotype flags, and variant flags.

Some genotypes are flagged by CEGH-Filter but not considered in the final counts to generate allele frequencies. Multiallelic informative genotypes (0/. and ./1), including SNVs, indels, and spanning deletions, are not included in allele frequency calculations due to dependency on other variants at the same site. Also, non-pseudoautosomal regions (non-PAR) loci along the X chromosome that harbor genotypes called as heterozygous state in male individuals are not accounted for, since they are expected to be hemizygous in those sites. These unexpected genotypes may be explained as false positives due to call errors or contamination, or else as true positives (duplications, aneuploidies, or mosaicism).

Although we consider further investigating those cases, at this point, we opted to exclude both multiallelic genotypes and non-PAR heterozygous in males from the counts and allelic frequency calculations, without, however, excluding the variant row per se. Therefore, some variants will appear to have zero allele frequency, even in non-FDP or non-FAB sites.

For overall counts (Extended Data Fig. 1), the high confidence dataset considered only GATK 'PASS' and CEGH 'vSR', 'SR', and 'WK'. Therefore, among the shadowed branch of counts, there are variants likely to be true positives, but rather fall within sites containing lower confidence calls. pLOF variants classified by LOFTEE[1] were considered irrespective of confidence label (HC or LC), since LC contains variants with at least one filter failed but can be a true positive. Additional evidence for nonsense-mediated decay or non-canonical splice sites can enrich the classification (https://github.com/konradjk/loftee). In the clinical analyses dataset, we have considered any CEGH-filter flag except for 'FDP' and 'FAB', since manual curation took place in the final step.

Annotation of variants per predicted function yielded the expected higher number of intergenic (53.7%) and intronic (35.3%), whereas coding variants represent less than 1% with 635 thousand variants (Supplementary Table 4).

**Supplementary Table 4**. Variant counts per predicted function.

| Variants Predicted Function | Counts | % Group | % All |
|---|---|---|---|
| **Coding and splicing within genes** | **635689** | | **0.81** |
| Nonsynonymous SNV | 329744 | 51.87 | 0.42 |
| Synonymous SNV | 221436 | 34.83 | 0.28 |
| Splicing | 28526 | 4.49 | 0.04 |
| Exonic;splicing | 25752 | 4.05 | 0.03 |
| Nonframeshift deletion | 8218 | 1.29 | 0.01 |
| Frameshift deletion | 7196 | 1.13 | 0.01 |
| Stopgain | 6677 | 1.05 | 0.01 |
| Nonframeshift insertion | 4144 | 0.65 | 0.01 |
| Frameshift insertion | 3638 | 0.57 | 0.00 |
| Stoploss | 356 | 0.06 | 0.00 |
| Splicing;splicing | 2 | 0.00 | 0.00 |
| **Noncoding within genes** | **29064505** | | **37.08** |
| Intronic | 27186280 | 93.54 | 34.68 |
| UTR3 | 647982 | 2.23 | 0.83 |
| Downstream | 531982 | 1.83 | 0.68 |
| Upstream | 519497 | 1.79 | 0.66 |
| UTR5 | 157192 | 0.54 | 0.20 |
| Upstream;downstream | 21185 | 0.07 | 0.03 |
| UTR5;UTR3 | 379 | 0.00 | 0.00 |
| Intronic;intronic | 8 | 0.00 | 0.00 |
| **Noncoding genes** | **4829729** | | **6.16** |
| ncRNA_intronic | 4544805 | 94.10 | 5.80 |
| ncRNA_exonic | 275495 | 5.70 | 0.35 |
| ncRNA_exonic;splicing | 4734 | 0.10 | 0.01 |
| ncRNA_splicing | 4669 | 0.10 | 0.01 |
| ncRNA_UTR5 | 22 | 0.00 | 0.00 |
| ncRNA_intronic;ncrna_intronic | 4 | 0.00 | 0.00 |
| **Intergenic** | **41363727** | | **52.76** |
| Intergenic | 41363715 | 100.00 | 52.76 |
| Intergenic;intergenic | 12 | 0.00 | 0.00 |
| **Other** | **2499880** | | **3.19** |
| Spanning deletion | 2495642 | 99.83 | 3.18 |
| NA + unknown | 4238 | 0.17 | 0.01 |
| **All** | **78393530** | | |

### 3. Ancestry analyses

In ancestry inference analyses, data from 30 populations were used as parental references, among which 16 were Native American (Supplementary Table 5).

**Supplementary Table 5.** Parental population used for ancestry inferences.

| Population | Region/Parental Population | N | Reference |
|---|---|---|---|
| Luhya in Webuye, Kenya | Africa | 99 | Auton et al., 2015[6] |
| Yoruba in Ibadan, Nigeria | Africa | 108 | |
| Gambian in Western Divisions in the Gambia | Africa | 113 | |
| Mende in Sierra Leone | Africa | 85 | |
| Esan in Nigeria | Africa | 99 | |
| Han Chinese in Beijing, China | East Asia | 103 | |
| Southern Han Chinese | East Asia | 105 | |
| Chinese Dai in Xishuangbanna, China | East Asia | 93 | |
| Kinh in Ho Chi Minh City, Vietnam | East Asia | 99 | |
| Finnish in Finland | Europe | 99 | |
| British in England and Scotland | Europe | 91 | |
| Toscani in Italia | Europe | 107 | |
| Iberian Population in Spain | Europe | 107 | |
| Utah Residents (CEPH) with Northern and Western European Ancestry | Europe | 99 | |
| Aimara in Peru | Native American | 11 | Borda et al., 2020[7] |
| Ashaninka in Peru | Native American | 33 | |
| Awajun in Peru | Native American | 22 | |
| Candoshi in Peru | Native American | 16 | |
| Chopccas in Peru | Native American | 7 | |
| Lamas in Peru | Native American | 17 | |
| Matses in Peru | Native American | 11 | |
| Matsiguenka in Peru | Native American | 3 | |
| Mache in Peru | Native American | 9 | |
| Nahua in Peru | Native American | 2 | |
| Qeros in Peru | Native American | 12 | |
| Quechua in Peru | Native American | 1 | |
| Shimaa in Peru | Native American | 23 | |
| Shipibo in Peru | Native American | 14 | |
| Tallanes in Peru | Native American | 30 | |
| Uros in Peru | Native American | 12 | |

### 4. Clinical findings

#### 4.1. Strategy

As initial classification criteria, we flagged all variants harbored by 4,250 OMIM disease genes (Supplementary Table 6) with ClinVar pathogenic assertions (Pathogenic, Likely Pathogenic, or Pathogenic/Likely Pathogenic) or predicted as promoting any loss of function consequence by LOFTEE algorithm. A total of 5,142 variants met the criteria (4,096 SNVs and 1,046 >1bp indels) (Pathogenicity analyses summarized in Supplementary Figure 4).

**Supplementary Table 6.** OMIM Disease genes.
Large table displayed only on Supplementary Tables worksheet.

#### 4.2. Frequencies of variants with potential clinical relevance

Although 10.6% of these variants are absent from population databases (gnomAD, dbSNP, and 1000 genomes), and most of which are indels, the remaining are mainly rare single-nucleotide substitutions (frequencies ≤ 0.001) (Supplementary Table 7).

**Supplementary Table 7.** Distribution of variants identified in SABE 1171 cohorts with potential clinical relevance in OMIM Disease genes absent and present in database, per frequency.
Large table displayed only on Supplementary Tables worksheet.

Few exceptions reach over 0.1 Among these high-frequency variants, we highlight the five asserted as pathogenic (Supplementary Table 8): (a) rs429358 *APOE* p.C130R (NM_000041) with a frequency of 0.13, which should be considered in phase with rs7412 p.R176C to the well-known functional haplotypes (ε2, ε3, and ε4) associated with late-onset Alzheimer's and type III hyperlipoproteinemia; **C.** rs17261572 and rs1566734, which were asserted as pathogenic before large allelic frequency datasets and community-based consensual criteria such as recommendations provided by American College of Human Genetics and Genomics (ACMG) were available[8]; and (c) variants classified as risk factors (lower penetrance by definition) in sporadic breast cancer multifactorial susceptibility (rs2046210) or in glycine metabolism on a digenic model of inheritance (rs35329108). Therefore, pathogenic assertions should be considered with caution.

**Supplementary Table 8**.Pathogenic variants in OMIM Disease genes with frequencies above 10% in SABE dataset.
Large table displayed only on Supplementary Tables worksheet.

### 4.3. Context of variants

These 5,142 variants fall within 1,949 unique genes and 168 intergenic regions or overlapping genes, 98% of which harbor no more than 10 variants (Supplementary Table 9). Most genes are annotated as either recessive or non-monogenic modes of inheritance, but a considerable amount of genes (749) are described as having either dominant or both dominant and recessive inheritance (Supplementary Table 10).

**Supplementary Table 9**. Number of genes and regions that harbor one or more variant with potential clinical relevance

| Variants per gene/region | Number of genes/regions | % |
|---|---|---|
| 1 | 992 | 46.86 |
| 2 | 493 | 23.29 |
| 3 | 257 | 12.14 |
| 4 | 135 | 6.38 |
| 5 | 81 | 3.83 |
| (5-10] | 126 | 5.95 |
| (10-20] | 26 | 1.23 |
| >20 | 7 | 0.33 |
| **Total** | 2117 | 100 |

**Supplementary Table 10**. Number of genes that harbor variants with potential clinical relevance per inheritance mode

| Inheritance mode (OMIM) | Number of genes |
|---|---|
| AD/XLD only | 437 |
| AR/XLR only | 1170 |
| AD and AR, XLD and XLR | 312 |
| Somatic mutation | 31 |
| Multifactorial | 12 |
| Mitochondrial | 6 |
| Digenic (Dominant or Recessive) | 20 |
| Somatic mosaicism | 1 |
| **Total** | 1989 |

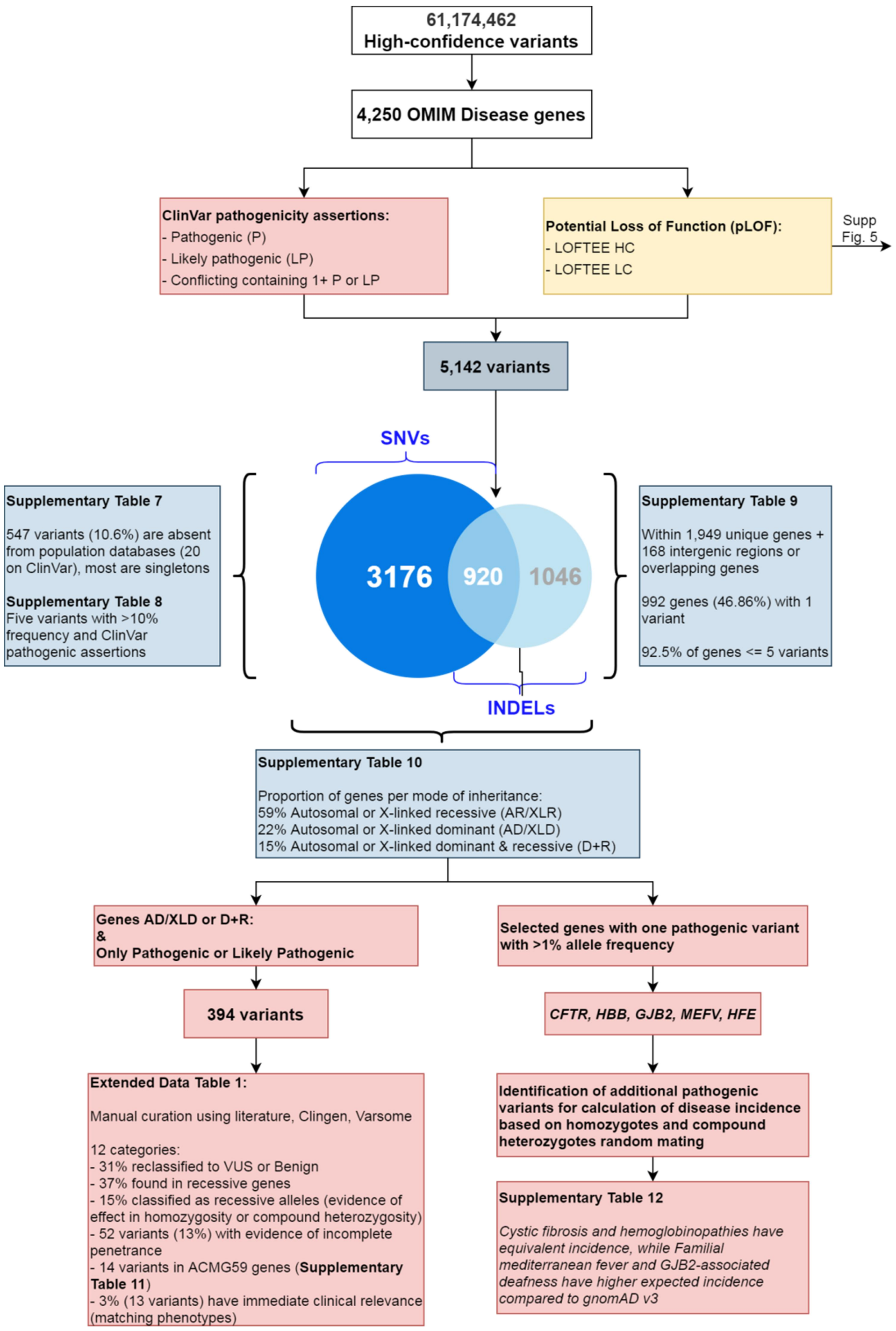

**Supplementary Fig. 4. Filtering strategies for the identification of variants of potential clinical relevance and indication of downstream results.** Among high-confidence variants, we have identified a total of 5,142 variants within 4,250 OMIM Disease genes that were found to have pathogenic, likely pathogenic, or conflicting containing at least one pathogenic ClinVar assertions, or classified as potential loss of function (pLOF). Downstream analyses pointed that: over 10% are absent from population databases, but most are singleton or low frequency (Supplementary Table 7-8); most genes contain up to five variants (Supplementary Table 9); most genes are annotated to associate with recessive modes of inheritance (Supplementary Table 10); manual curation of variants initially classified as pathogenic or likely pathogenic in genes of dominant inheritance can be either reclassified or fall indeed in recessively inherited conditions or else are required to be in trans with a more deleterious variant (Extended Data Tab. 1); SABE cohort recapitulates variant-based incidence of recessive disorders that are more prevalent in European or African populations (Supplementary Table 12).

#### 4.4. Manual curation of variant pathogenicity on genes associated with dominant mode of inheritance

In order to identify individuals carrying variants with potential clinical implications, including the reassessment of related phenotypes to support the analyses, we have filtered a total of 394 variants asserted as either 'Pathogenic' or 'Likely Pathogenic' (P/LP) in genes annotated to have a dominant mode of inheritance only, and in genes with more than one mode of inheritance, including dominant or monoallelic. Manual curation aiming reclassification of pathogenicity using ACMG criteria was performed and included family-based information described in the available literature, evidence details on the original assertions, and allele frequency. A total of 123 variants (31%) were reclassified as non-pathogenic assertions (benign, likely benign or unknown significance), and the remaining 271 kept as P/LPpathogenic or likely pathogenic. Among the latter, literature validation and matching phenotypes, when available, enabled further characterization of variants to either a reported reduced penetrance, non-dominant mode (of the specific allele or gene), or associated to clinical features that are not severe enough to cause mortality before the average age of subjects (Extended Data Tab. 1).

#### 4.5. Manual curation of variant pathogenicity on ACMG 59 actionable genes

We also analyzed P/LP variants in 59 actionable genes following ACMG recommendation[9] and found 14 variants distributed in heterozygosity in different individuals all in the heterozygous state (Supplementary Table 11), among which *BRCA2* and *RYR1* harbor four variants each. Ten variants were classified using the above-mentioned protocol as pathogenic with reported incomplete penetrance; three were described as pathogenic only when in *trans* with another pathogenic variant (recessive alleles), and one potential phenotypic match (outcome compatible with finding) in *LDLR*.

**Supplementary Table 11.** Pathogenic findings on ACMG-59 genes.
Large table displayed only on Supplementary Tables worksheet.

#### 4.6. Assessment of variant-based incidence pathogenicity on selected genes associated with recessive mode of inheritance

To roughly estimate the incidence using counts of heterozygotes from SABE and gnomAD global and population-specific datasets, we selected five genes associated with prevalent monogenic clinical phenotypes: cystic fibrosis (*CFTR*), hemoglobinopathies (*HBB*), deafness (*GJB2*), familial Mediterranean fever (*MEFV*), and hemochromatosis (*HFE*) (Supplementary Table 12).

**Supplementary Table 12.** Variant-based incidence of selected genes associated with recessively inherited disorders.
Large table displayed only on Supplementary Tables worksheet.

#### 4.7. Distribution of potential loss of function variants within OMIM Disease genes

There are 4,000 potential loss of function (pLOF) variants in SABE that fall within OMIM Disease genes, of which 3,704 are 'non-Benign', which excludes ClinVar benign, likely benign, or conflicting assertions that lack pathogenic entries (Supplementary Fig. 5). We have found a normal distribution of individual loads of pLOF variants in heterozygous state (Supplementary Fig. 6A) and homozygous state (Supplementary Fig. 6B), and a Poisson distribution of variants with one or more pathogenic assertions (Supplementary Fig. 6C), with medians of 55, 25, and 1 variants per person, respectively. A comparison of allele frequencies of these variants between SABE and gnomAD revealed a high correlation regardless of pLI contexts (Supplementary Fig. 7A).

Assuming a higher proportion of false positives among indels, mainly longer ones, we have filtered only pLOF variants produced by single nucleotide variants (substitutions or 1bp long indels), with allele frequency of 5% or lower on SABE and flagged as high-confidence by LOFTEE. These strict filtering criteria yielded a total of 1,853 pLOF variants, which were Poisson distributed with a median of 5 in heterozygosity per individual (Supplementary Fig. 6D), a median of 0 in homozygosity (although 20% have one or two, Supplementary Fig. 6E) and a median of 0 with one or more pathogenic assertion (39% have one to four, Supplementary Fig. 6F). A high correlation with gnomAD frequencies is maintained in this subset (Supplementary Fig. 7B). Further detailed analysis of pLOF variants within genes of pLI $\geq 0.7$ with higher differences in allele frequencies showed that they were either long indels multiallelic or homopolymers flagged as low confidence or in low-quality sites in gnomAD. Also, annotation of variants in intergenic regions may be wrongly attributed and lead to spurious flagging (Supplementary Fig. 7C). Therefore, regardless of the dataset quality cutoff, we have found non-deviant frequencies as compared with gnomAD.

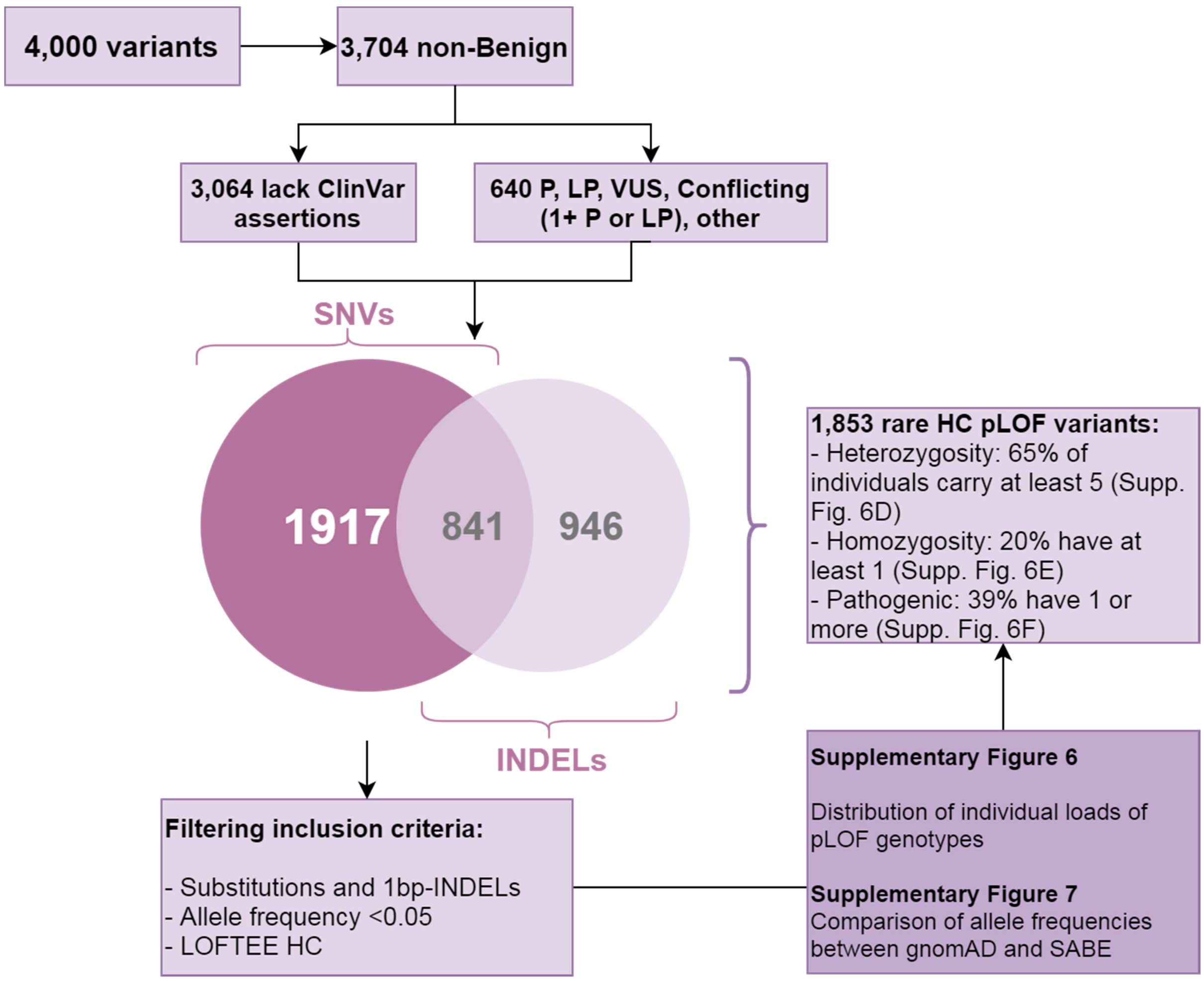

**Supplementary Fig. 5. Filtering strategies for identification of variants of potential loss of function.** Among high-confidence variants, we have identified 5,142 variants within 4,250 OMIM Disease genes, 4,000 of which were classified as potential loss of function (pLOF). A subset of any pLOF non-Benign variants corresponds to 3,704 variants with any ClinVar non Benign assertion (640 variants) plus 3,064 variants that lack any assertions. Substitutions, 1bp-indels, and indels >1bp were analyzed for the distribution of individual loads (Supplementary Fig. 6A, 6B, and 6C) and allele frequencies compared to gnomAD (Supplementary Fig. 7A). Further filtering to remove indels >1bp, common variants and LC-flagged pLOFs yielded 1,854 variants, which individual load distributions were also analyzed (Supplementary Fig. 6D, 6E, and 6F) as well as frequency comparison to gnomAD (Supplementary Fig. 7B).

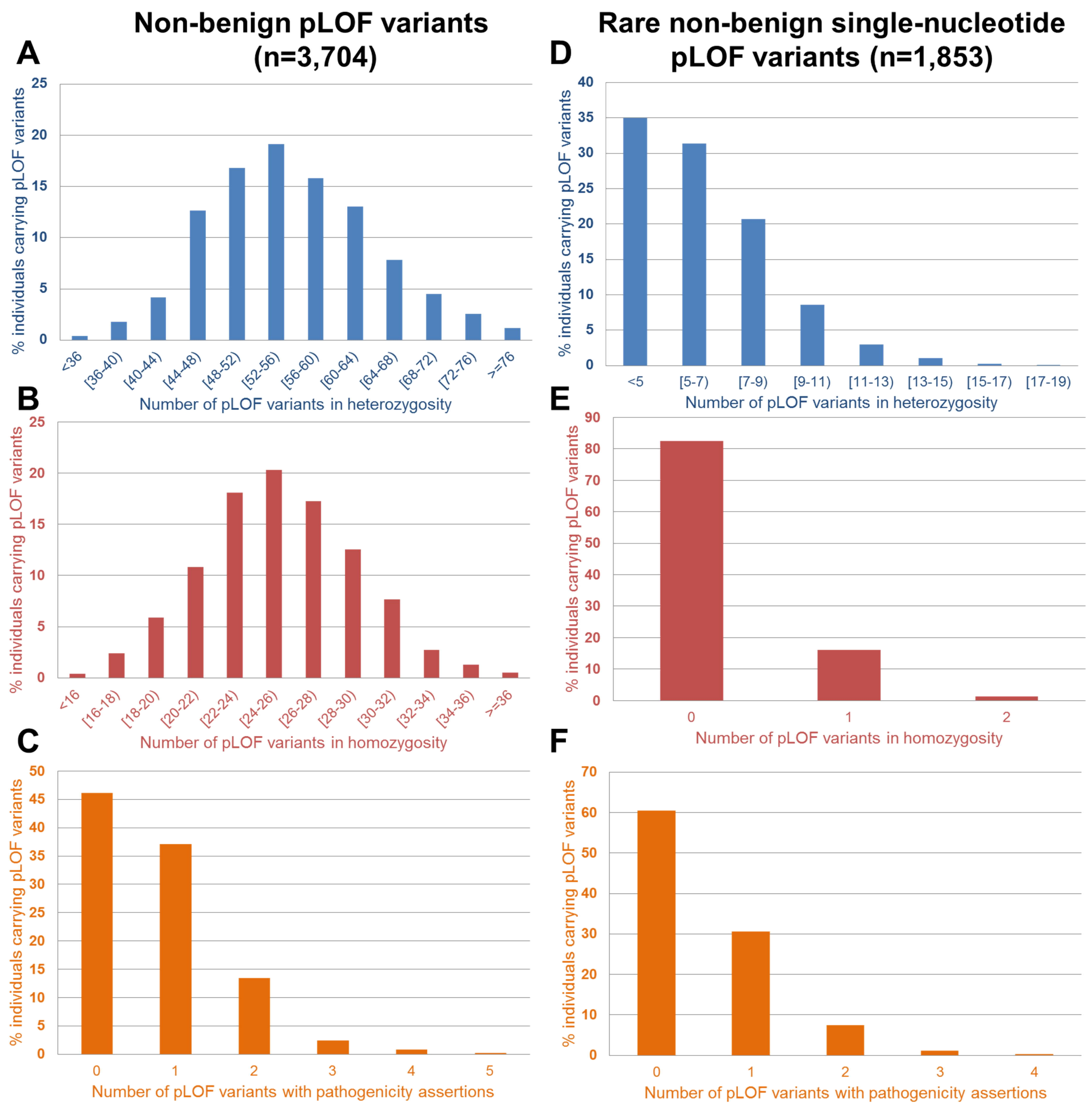

**Supplementary Fig. 6. Distribution of individual loads of potential loss of function (pLOF) variants.** Left panels: a subset of any non-Benign pLOF (ClinVar non-Benign assertions plus variants that lack any assertions) variants. Histogram of individual loads of pLOF variants in **(A)** heterozygosity, **(B)** homozygosity, and **(C)** variants with pathogenic assertions on ClinVar (including Pathogenic, Likely Pathogenic and Conflicting containing one or more pathogenic entries). Right panels: a subset of pLOF variants that are single nucleotide substitutions or 1bp-indels below 5% SABE cohort frequency and flagged as LOFTEE HC. Histogram of individual loads of pLOF variants in **(D)** heterozygosity, **(E)** homozygosity, and **(F)** variants with pathogenic assertions on ClinVar.

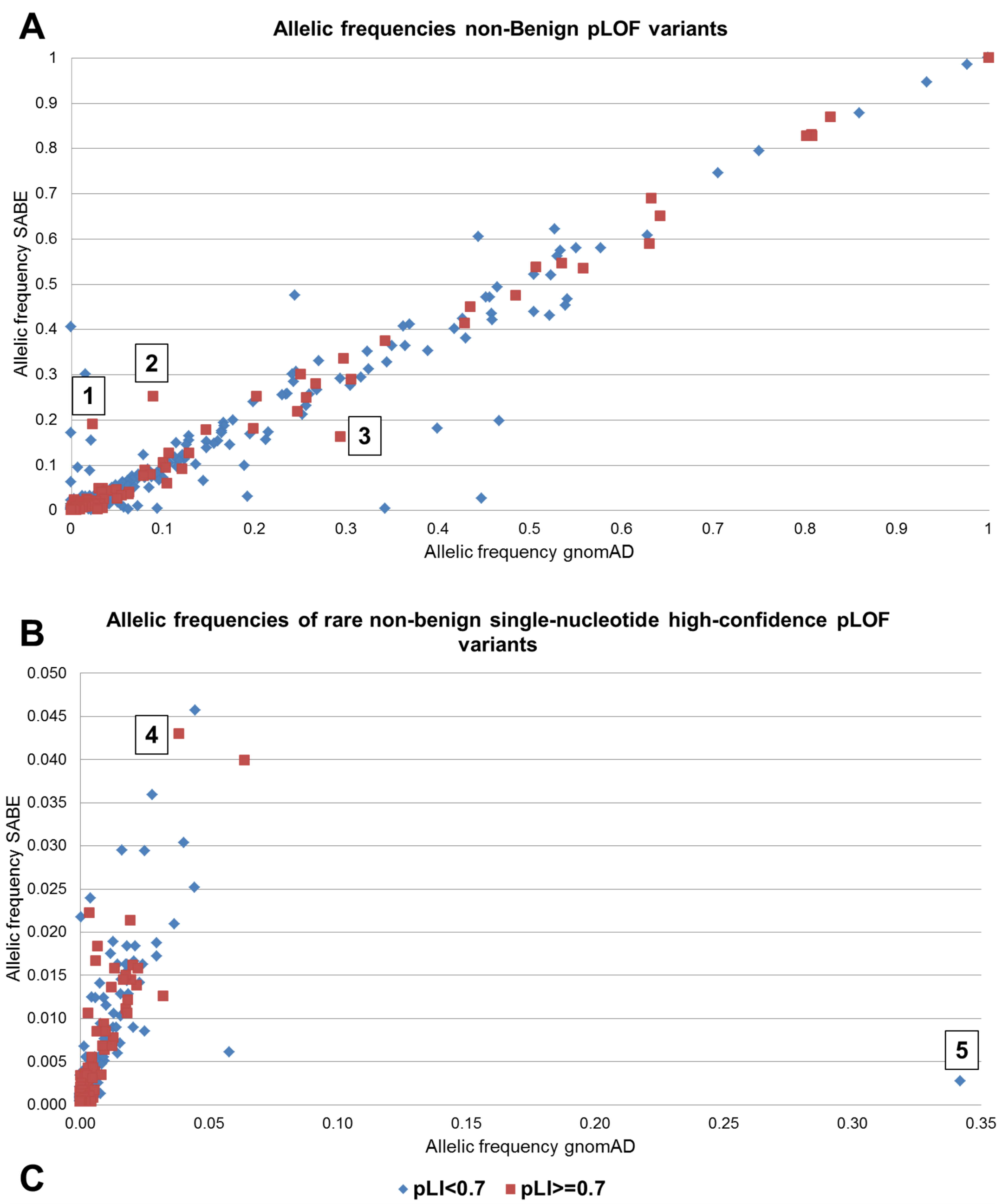

C

| # in Plot | Filter | Chr | Position | Ref | Alt | Gene | pLI | Allele frequencies SABE | gnomAD v2.2 | Cause of disparities |
|---|---|---|---|---|---|---|---|---|---|---|
| 1 | All pLOF Non Benign | 9 | 128694106 | - | TATA | *SET* | 0.96 | 0.1905 | 0.0237 | Multiallelic INDEL with Low Confidence |
| 2 | All pLOF Non Benign | 12 | 70353914 | A | - | *CNOT2* | 1.00 | 0.2516 | 0.0893 | Multiallelic INDEL in poly-A Homopolymer |
| 3 | All pLOF Non Benign | 16 | 89233469 | TGCAACTCAACTCACACTGCGTA | - | *ANKRD11;ZNF778* | 1.00;0 | 0.1619 | 0.2939 | Intergenic, pLI from neighbor gene |
| 4 | Rare HC pLOF SNVs | X | 153481313 | G | - | *BGN; HAUS7* | 0.92; 0.96 | 0.0430 | 0.0385 | pLOF annotation incorrect |
| 5 | Rare HC pLOF SNVs | 3 | 10046724 | G | - | *FANCD2* | 0.00 | 0.0027 | 0.3422 | Low quality site (absence of homozygotes) |

**Supplementary Fig. 7. Comparison of allele frequencies between pLOFs found in SABE and gnomAD (v2.2), within genes of pLI>=0.7 and pLI<0.7. A.** Subset of non-Benign variants provided a comparison of pLOF frequencies of up to 100%. **B.** Rare single nucleotide variants flagged as HC on LOFTEE. **C.** Five examples of deviation were manually verified in gnomAD and explained by context leading to calls or annotations.

### 5. WGS Imputation

**Supplementary Table 13.** Number of SNPs per chromosome in each reference panel

| Chromosome | Number of SNPs in each reference panel | | |
|---|---|---|---|
| | SABE | 1KGP3 | SABE+1KGP3 |
| 1 | 3,996,540 | 6,191,833 | 7,939,598 |
| 2 | 4,406,141 | 6,790,551 | 8,726,263 |
| 3 | 3,690,698 | 5,641,493 | 7,257,393 |
| 4 | 3,596,780 | 5,477,810 | 7,025,900 |
| 5 | 3,325,574 | 5,115,036 | 6,553,929 |
| 6 | 3,174,612 | 4,863,337 | 6,218,835 |
| 7 | 2,945,373 | 4,511,408 | 5,792,582 |
| 8 | 2,898,843 | 4,425,449 | 5,683,312 |
| 9 | 2,204,350 | 3,384,360 | 4,346,771 |
| 10 | 2,514,657 | 3,874,259 | 4,950,281 |
| 11 | 2,509,089 | 3,881,791 | 4,972,826 |
| 12 | 2,421,384 | 3,745,465 | 4,800,039 |
| 13 | 1,806,750 | 2,760,845 | 3,534,231 |
| 14 | 1,644,170 | 2,548,903 | 3,259,739 |
| 15 | 1,484,079 | 2,301,453 | 2,949,517 |
| 16 | 1,655,523 | 2,548,920 | 3,289,287 |
| 17 | 1,427,164 | 2,209,149 | 2,855,082 |
| 18 | 1,432,958 | 2,189,529 | 2,800,626 |
| 19 | 1,106,201 | 1,738,824 | 2,237,376 |
| 20 | 1,180,936 | 1,817,492 | 2,329,578 |
| 21 | 664,678 | 1,045,269 | 1,324,116 |
| 22 | 679,009 | 1,059,079 | 1,357,134 |
| TOTAL | 50,765,509 | 78,229,219 | 100,204,415 |

**Supplementary Table 14.** Comparison between target haplotype phase inferences with different reference haplotypes using the number of imputed SNPs for chromosomes 15, 17, 20, and 22. Target 2.5M EPIGEN

| Imputation Reference Panel | SABE | | 1KGP3 | | SABE+1KGP3 | |
|---|---|---|---|---|---|---|
| Number of variants | Total | info score ≥ 0.8 | Total | info score ≥ 0.8 | Total | info score ≥ 80% |
| Chr 15 | 1,481,369 | 600,332 | 2,297,258 | 799,440 | 2,943,434 | 951,917 |
| Chr 17 | 1,424,402 | 512,055 | 2,204,724 | 738,586 | 2,849,458 | 866,547 |
| Chr 20 | 1,180,618 | 417,420 | 1,816,925 | 615,816 | 2,328,821 | 703,545 |
| Chr 22 | 676,922 | 229,932 | 1,049,542 | 351,164 | 1,345,756 | 402,144 |

**Supplementary Table 15.** Comparison between target haplotype phase inferences with different reference haplotypes using the number of imputed SNPs for chromosomes 15, 17, 20, and 22. Target 2.5M SALVADOR

| Imputation Reference Panel | SABE | | 1KGP3 | | SABE+1KGP3 | |
|---|---|---|---|---|---|---|
| Number of variants | Total | info score ≥ 0.8 | Total | info score ≥ 0.8 | Total | info score ≥ 80% |
| Chr 15 | 1,481,369 | 605,791 | 2,297,258 | 799,308 | 2,943,434 | 921,805 |
| Chr 17 | 1,424,402 | 519,213 | 2,204,724 | 740,926 | 2,849,458 | 851,122 |
| Chr 20 | 1,180,618 | 423,629 | 1,816,925 | 616,333 | 2,328,821 | 691,770 |
| Chr 22 | 676,922 | 234,225 | 1,049,542 | 352,320 | 1,345,756 | 395,895 |

**Supplementary Table 16.** Comparison between target haplotype phase inferences with different reference haplotypes using the number of imputed SNPs for chromosomes 15, 17, 20, and 22. Target 2.5M PELOTAS

| Imputation Reference Panel | SABE | | 1KGP3 | | SABE+1KGP3 | |
|---|---|---|---|---|---|---|
| Number of variants | Total | info score ≥ 0.8 | Total | info score ≥ 0.8 | Total | info score ≥ 80% |
| Chr 15 | 1,481,369 | 594,208 | 2,297,258 | 763,178 | 2,943,434 | 898,979 |
| Chr 17 | 1,424,402 | 509,774 | 2,204,724 | 711,297 | 2,849,458 | 826,780 |
| Chr 20 | 1,180,618 | 414,600 | 1,816,925 | 594,121 | 2,328,821 | 673,256 |
| Chr 22 | 676,922 | 228,107 | 1,049,542 | 339,416 | 1,345,756 | 384,628 |

**Supplementary Table 17.** Comparison between target haplotype phase inferences with different reference haplotypes using the number of imputed SNPs for chromosomes 15, 17, 20, and 22. Target 2.5M BAMBUI

| Imputation Reference Panel | SABE | | 1KGP3 | | SABE+1KGP3 | |
|---|---|---|---|---|---|---|
| Number of variants | Total | info score ≥ 0.8 | Total | info score ≥ 0.8 | Total | info score ≥ 80% |
| Chr 15 | 1,481,369 | 573,646 | 2,297,258 | 692,257 | 2,943,434 | 803,886 |
| Chr 17 | 1,424,402 | 495,561 | 2,204,724 | 648,734 | 2,849,458 | 746,572 |
| Chr 20 | 1,180,618 | 403,534 | 1,816,925 | 541,084 | 2,328,821 | 605,172 |
| Chr 22 | 676,922 | 224,813 | 1,049,542 | 314,774 | 1,345,756 | 354,075 |

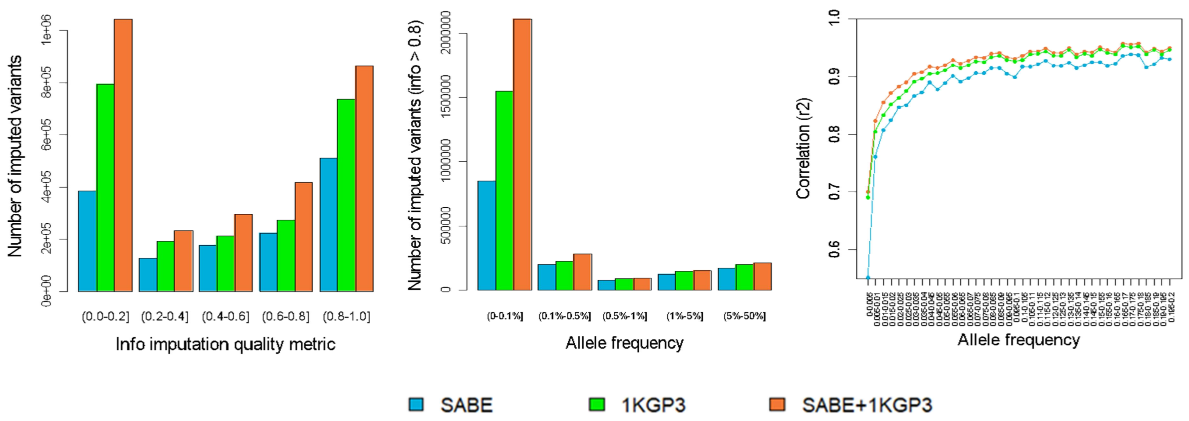

**Supplementary Fig. 8.** Comparison of imputation performance of SABE, 1KGP3, and SABE+1KGP3 reference panels using the Omni 2.5M array data for 6,487 Brazilians from **EPIGEN for chromosome 17** as target panel. **A.** The total number of imputed variants across different classes of the info score quality metric. **B.** The total number of imputed variants with info score ≥ 0.8 across the allele frequency spectrum. **C.** Improvement in imputation accuracy as a function of MAF for the target dataset after imputation (MAF from 0 to 0.2, bin sizes of 0.005).

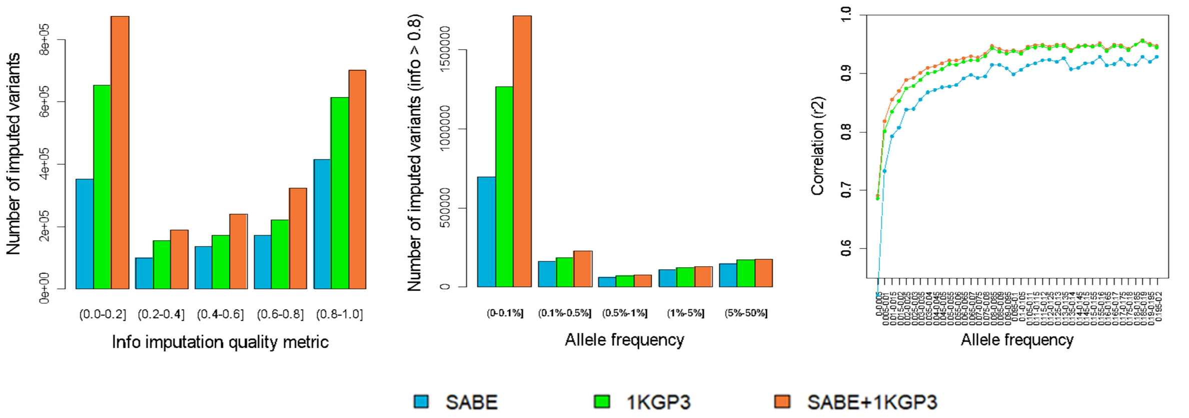

**Supplementary Fig. 9.** Comparison of imputation performance of SABE, 1KGP3, and SABE+1KGP3 reference panels using the Omni 2.5M array data for 6,487 Brazilians from **EPIGEN for chromosome 20** as target panel. **A.** The total number of imputed variants across different classes of the info score quality metric. **B.** The total number of imputed variants with info score ≥ 0.8 across the allele frequency spectrum. **C.** Improvement in imputation accuracy as a function of MAF for the target dataset after imputation (MAF from 0 to 0.2, bin sizes of 0.005).

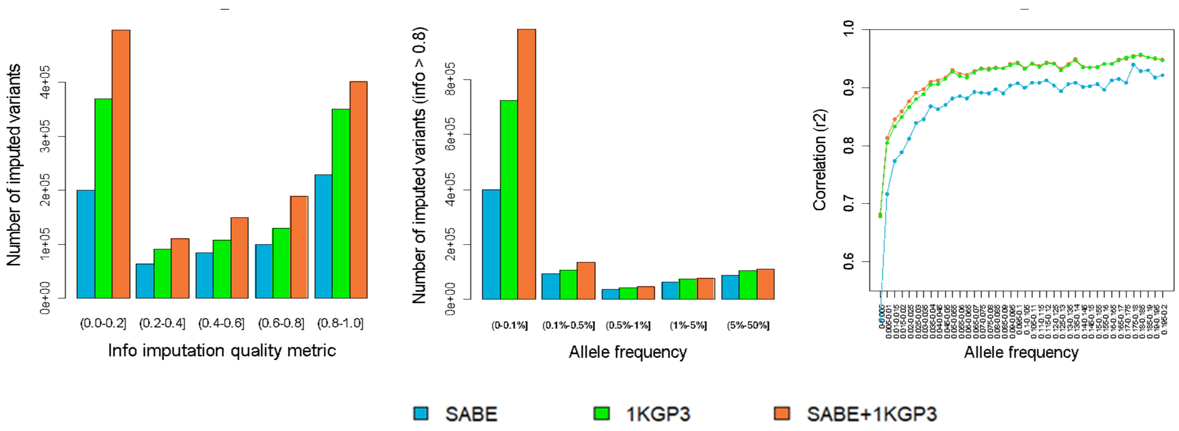

**Supplementary Fig. 10.** Comparison of imputation performance of SABE, 1KGP3, and SABE+1KGP3 reference panels using the Omni 2.5M array data for 6,487 Brazilians from **EPIGEN for chromosome 22** as target panel. **A.** The total number of imputed variants across different classes of the info score quality metric. **B.** The total number of imputed variants with info score ≥ 0.8 across the allele frequency spectrum. **C.** Improvement in imputation accuracy as a function of MAF for the target dataset after imputation (MAF from 0 to 0.2, bin sizes of 0.005).

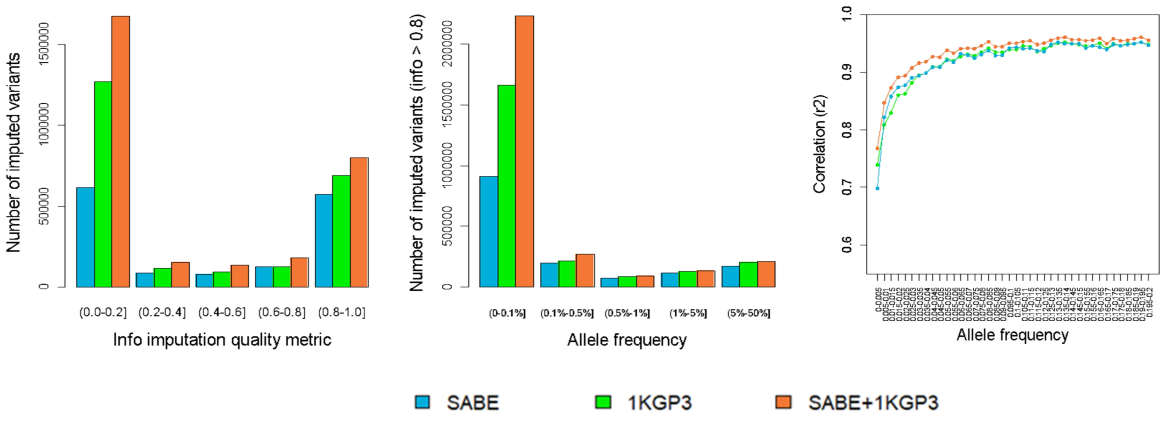

**Supplementary Fig. 11.** Comparison of imputation performance of SABE, 1KGP3, and SABE+1KGP3 reference panels using the Omni 2.5M array data for 6,487 Brazilians from **BAMBUÍ for chromosome 15** as target panel. **A.** The total number of imputed variants across different classes of the info score quality metric. **B.** The total number of imputed variants with info score ≥ 0.8 across the allele frequency spectrum. **C.** Improvement in imputation accuracy as a function of MAF for the target dataset after imputation (MAF from 0 to 0.2, bin sizes of 0.005).

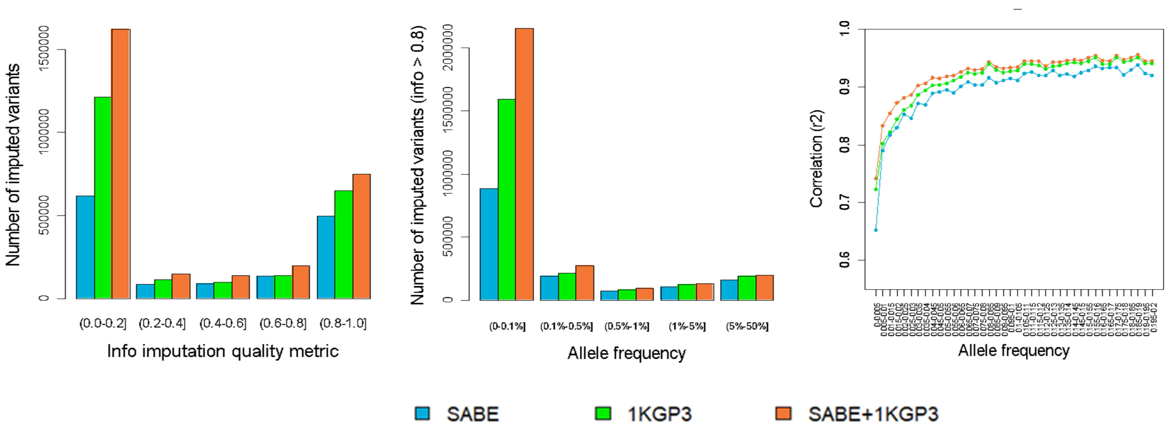

**Supplementary Fig. 12**. Comparison of imputation performance of SABE, 1KGP3, and SABE+1KGP3 reference panels using the Omni 2.5M array data for 6,487 Brazilians from **BAMBUÍ for chromosome 17** as target panel. **A.** The total number of imputed variants across different classes of the info score quality metric. **B.** The total number of imputed variants with info score ≥ 0.8 across the allele frequency spectrum. **C.** Improvement in imputation accuracy as a function of MAF for the target dataset after imputation (MAF from 0 to 0.2, bin sizes of 0.005).

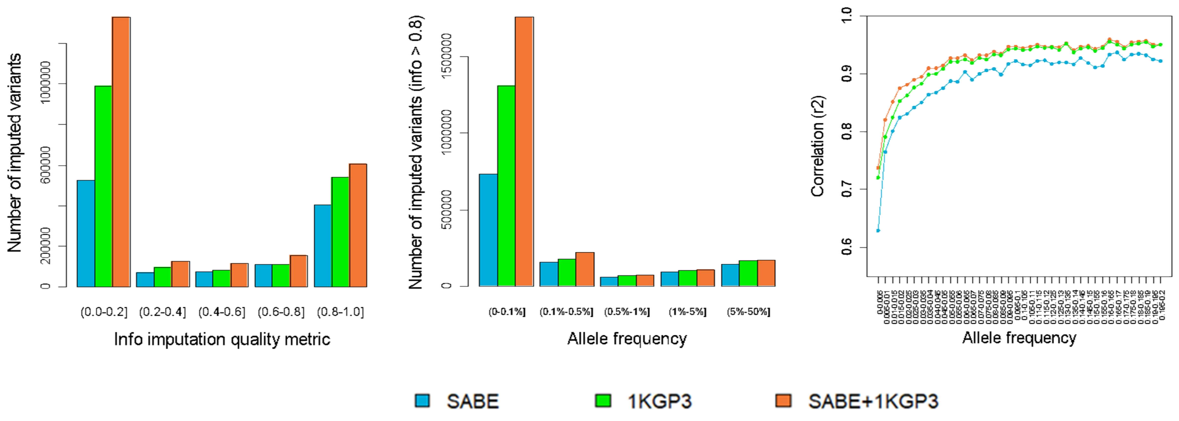

**Supplementary Fig. 13.** Comparison of imputation performance of SABE, 1KGP3, and SABE+1KGP3 reference panels using the Omni 2.5M array data for 6,487 Brazilians from **BAMBUÍ for chromosome 20** as target panel. **A.** The total number of imputed variants across different classes of the info score quality metric. **B.** The total number of imputed variants with info score ≥ 0.8 across the allele frequency spectrum. **C.** Improvement in imputation accuracy as a function of MAF for the target dataset after imputation (MAF from 0 to 0.2, bin sizes of 0.005).

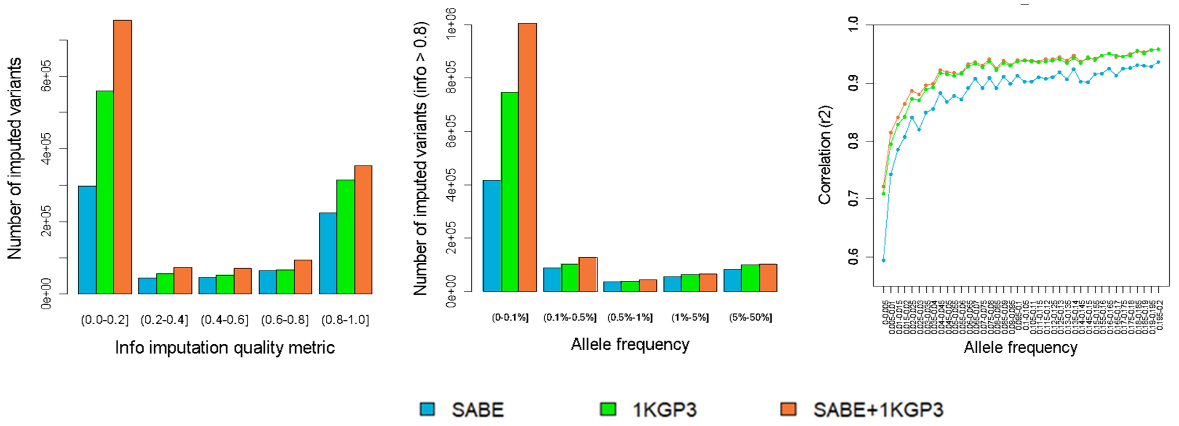

**Supplementary Fig. 14.** Comparison of imputation performance of SABE, 1KGP3, and SABE+1KGP3 reference panels using the Omni 2.5M array data for 6,487 Brazilians from **BAMBUÍ for chromosome 22** as target panel. **A.** The total number of imputed variants across different classes of the info score quality metric. **B.** The total number of imputed variants with info score ≥ 0.8 across the allele frequency spectrum. **C.** Improvement in imputation accuracy as a function of MAF for the target dataset after imputation (MAF from 0 to 0.2, bin sizes of 0.005).

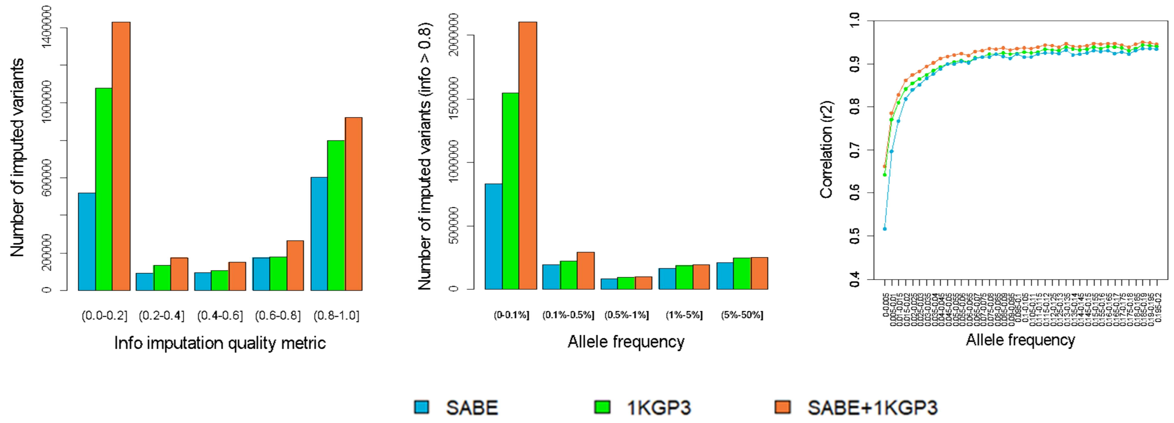

**Supplementary Fig. 15.** Comparison of imputation performance of SABE, 1KGP3, and SABE+1KGP3 reference panels using the Omni 2.5M array data for 6,487 Brazilians from **SALVADOR for chromosome 15** as target panel. **A.** The total number of imputed variants across different classes of the info score quality metric. **B.** The total number of imputed variants with info score ≥ 0.8 across the allele frequency spectrum. **C.** Improvement in imputation accuracy as a function of MAF for the target dataset after imputation (MAF from 0 to 0.2, bin sizes of 0.005).

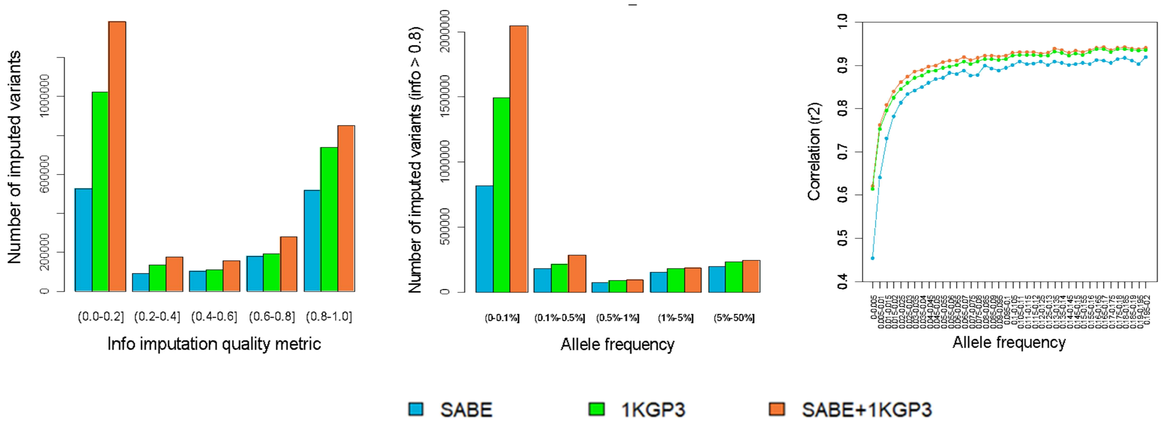

**Supplementary Fig. 16**. Comparison of imputation performance of SABE, 1KGP3, and SABE+1KGP3 reference panels using the Omni 2.5M array data for 6,487 Brazilians from **SALVADOR for chromosome 17** as target panel. **A.** The total number of imputed variants across different classes of the info score quality metric. **B.** The total number of imputed variants with info score ≥ 0.8 across the allele frequency spectrum. **C.** Improvement in imputation accuracy as a function of MAF for the target dataset after imputation (MAF from 0 to 0.2, bin sizes of 0.005).

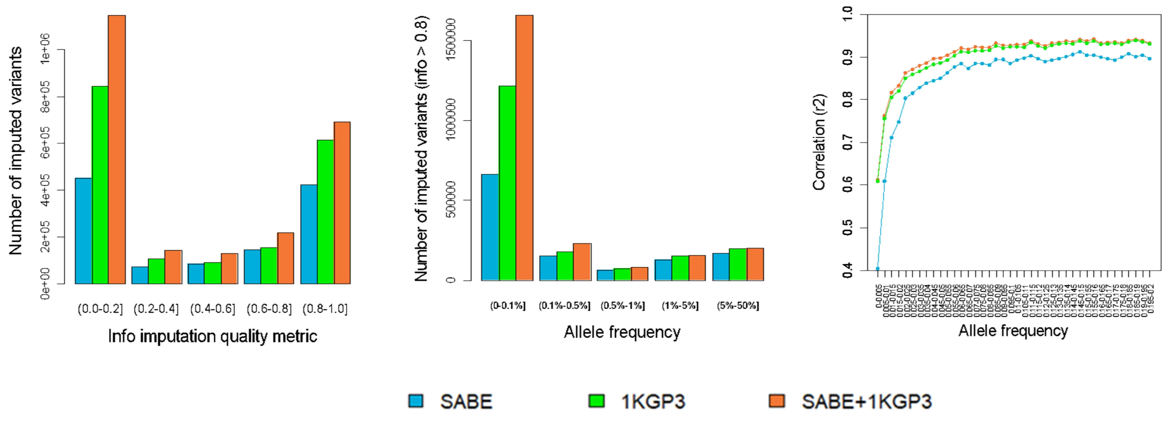

**Supplementary Fig. 17.** Comparison of imputation performance of SABE, 1KGP3, and SABE+1KGP3 reference panels using the Omni 2.5M array data for 6,487 Brazilians from **SALVADOR for chromosome 20** as target panel. **A.** The total number of imputed variants across different classes of the info score quality metric. **B.** The total number of imputed variants with info score ≥ 0.8 across the allele frequency spectrum. **C.** Improvement in imputation accuracy as a function of MAF for the target dataset after imputation (MAF from 0 to 0.2, bin sizes of 0.005).

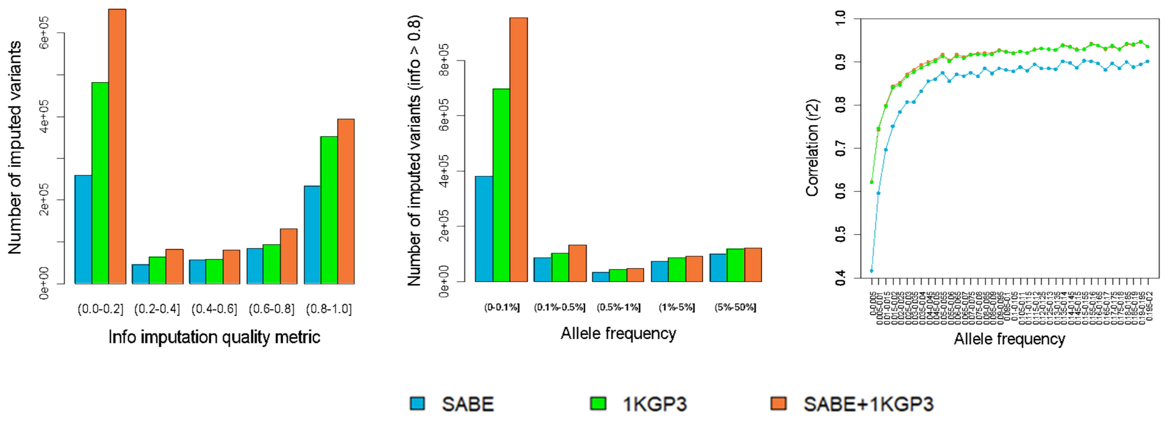

**Supplementary Fig. 18**. Comparison of imputation performance of SABE, 1KGP3, and SABE+1KGP3 reference panels using the Omni 2.5M array data for 6,487 Brazilians from **SALVADOR for chromosome 22** as target panel. **A.** The total number of imputed variants across different classes of the info score quality metric. **B.** The total number of imputed variants with info score ≥ 0.8 across the allele frequency spectrum. **C.** Improvement in imputation accuracy as a function of MAF for the target dataset after imputation (MAF from 0 to 0.2, bin sizes of 0.005).

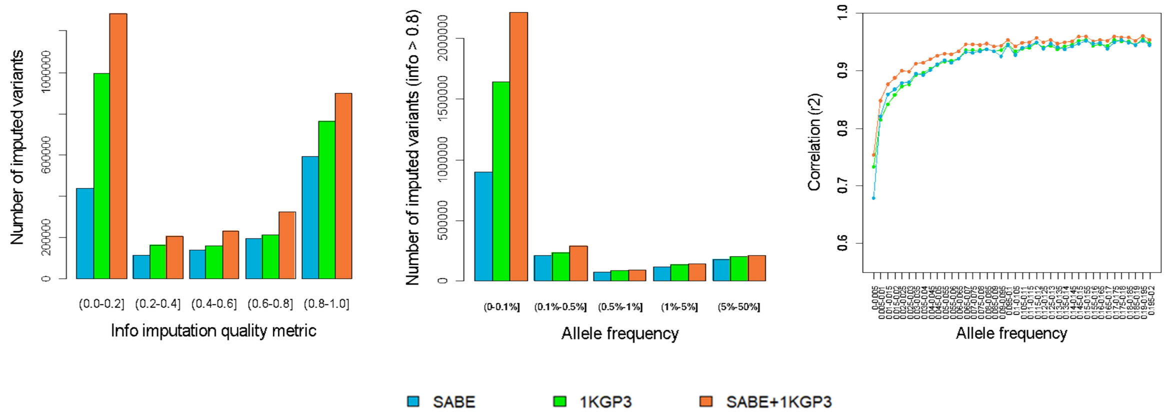

**Supplementary Fig. 19**. Comparison of imputation performance of SABE, 1KGP3, and SABE+1KGP3 reference panels using the Omni 2.5M array data for 6,487 Brazilians from **PELOTAS for chromosome 15** as target panel. **A.** The total number of imputed variants across different classes of the info score quality metric. **B.** The total number of imputed variants with info score ≥ 0.8 across the allele frequency spectrum. **C.** Improvement in imputation accuracy as a function of MAF for the target dataset after imputation (MAF from 0 to 0.2, bin sizes of 0.005).

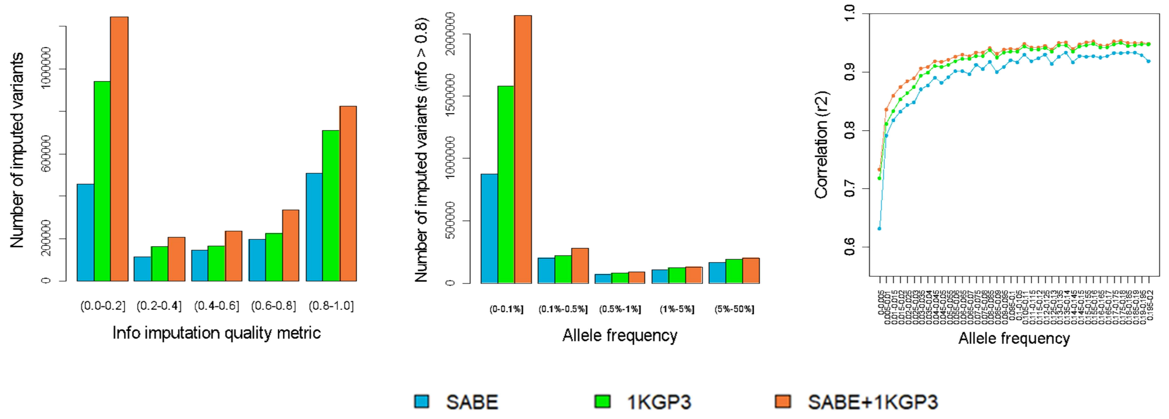

**Supplementary Fig. 20**. Comparison of imputation performance of SABE, 1KGP3, and SABE+1KGP3 reference panels using the Omni 2.5M array data for 6,487 Brazilians from **PELOTAS for chromosome 17** as target panel. **A.** The total number of imputed variants across different classes of the info score quality metric. **B.** The total number of imputed variants with info score ≥ 0.8 across the allele frequency spectrum. **C.** Improvement in imputation accuracy as a function of MAF for the target dataset after imputation (MAF from 0 to 0.2, bin sizes of 0.005).

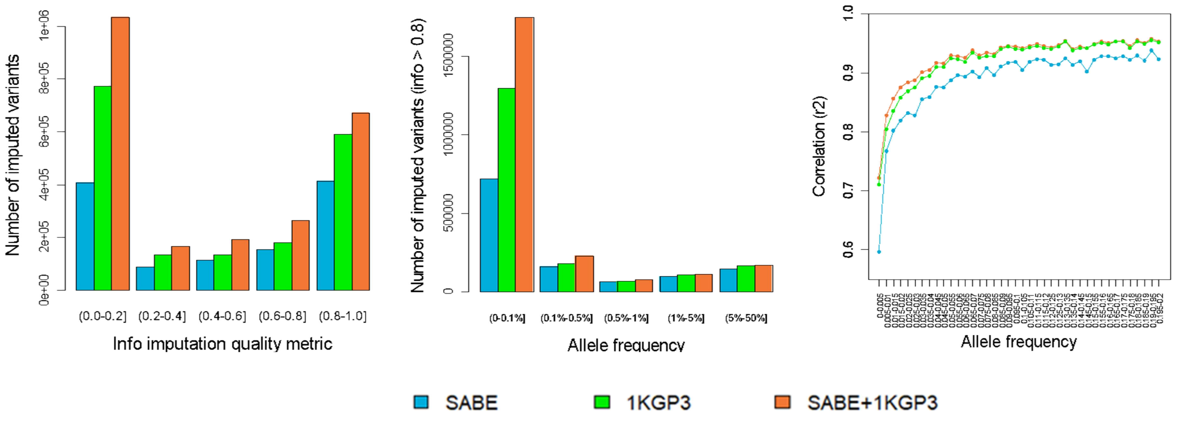

**Supplementary Fig. 21.** Comparison of imputation performance of SABE, 1KGP3, and SABE+1KGP3 reference panels using the Omni 2.5M array data for 6,487 Brazilians from **PELOTAS for chromosome 20** as target panel. **A.** The total number of imputed variants across different classes of the info score quality metric. **B.** The total number of imputed variants with info score ≥ 0.8 across the allele frequency spectrum. **C.** Improvement in imputation accuracy as a function of MAF for the target dataset after imputation (MAF from 0 to 0.2, bin sizes of 0.005).

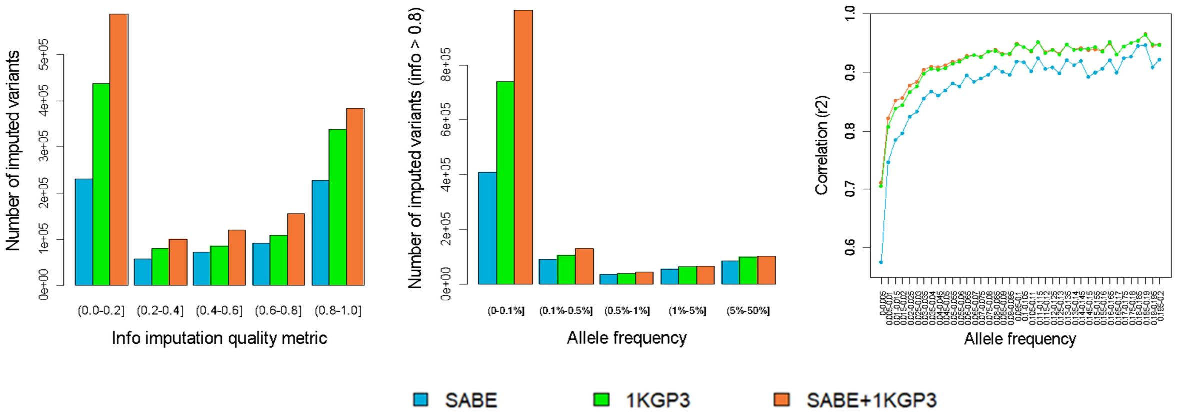

**Supplementary Fig. 22.** Comparison of imputation performance of SABE, 1KGP3, and SABE+1KGP3 reference panels using the Omni 2.5M array data for 6,487 Brazilians from **PELOTAS for chromosome 22** as target panel. **A.** The total number of imputed variants across different classes of the info score quality metric. **B.** The total number of imputed variants with info score ≥ 0.8 across the allele frequency spectrum. **C.** Improvement in imputation accuracy as a function of MAF for the target dataset after imputation (MAF from 0 to 0.2, bin sizes of 0.005).

### 6. HLA

The *hla-mapper* software was designed to optimize read mapping in HLA genes by comparing each read to a database of known HLA sequences and calculating where each read should be mapped or considered ambiguous[10]. This step is essential to get accurate genotype calls in the SNP-level for HLA genes. We used an updated version of this software, version 4, with support to intergenic sequences and faster processing WGS data.

After the map optimization, we used GATK HaplotypeCaller version 4.1.7 to call genotypes in the genome confidence model (GVCF), concatenating all samples together in a VCF file using GenotypeGVCFs. HaplotypeCaller can detect both SNPs and indels. For variant refinement, we noticed that the VQSR-AS approach does not produce reliable results for the HLA region by observing the filtered variants in samples with known HLA alleles. Because of that, we used a different approach for variant refinement and selection for HLA genes. We used a local program (vcfx) that introduces missing alleles in unbalanced genotypes (vcfx checked) and genotypes with a low likelihood (vcfx checkpl), and annotate each variant with some quantitative parameters such as the number of balanced heterozygous variants (when both alleles present similar depth of coverage), number of homozygous samples, the proportion of missing alleles, and others (vcfx evidence). Each variant that has not been approved by the vcfx evidence algorithm was evaluated manually. Each variant was annotated using the dbSNP dataset version 151.

We inferred haplotypes combining two computational strategies. First, we used GATK ReadBackedPhasing (RBP) to detect the phase between close heterozygous variants. The minimal Phase Quality Threshold was set to 500 (25x the default value). This procedure produced phase sets, i.e., blocks of known phases, but unphased among each other. RBP does not consider Multi-allelic variants, indels, and missing alleles. The second step consisted of inferring the full haplotypes using probabilistic models, but considering the phase sets detected by RBP. For that, we used an in-house software named phasex (available upon request), that uses Shapeit4[11] to phase bi-allelic variants considering the RBP's phase sets, in 20 independent runs with different seeds and using a single core per independent run, comparing the results afterward. The independent runs can be parallelized according to the number of cores on the computer. We preserved the haplotypes of all samples in which the same pair of haplotypes was observed in at least 18 runs ($P > 0.9$), passing these haplotypes forward to the next round. Iterations were performed until the number of samples with $P > 0.9$ no longer increased. Then, the haplotypes that have been detected are passed forward to the next step. In this next step, we use Beagle 4.1[12] to infer the final set of haplotypes, now including the multi-allelic variants. The same iteration procedure is used, with 20 independent runs and fixing haplotypes with $P > 0.9$. For each sample, we considered the haplotype with $P > 0.7$ after the last Beagle run. Shapeit4 and Beagle 4.1 also imputed the bi-allelic and multi-allelic missing alleles, which were introduced by the vcfx approach. It should be noted that we removed all singletons before the haplotyping procedure. This step is necessary because singletons are ambiguous by definition and impair haplotyping performance. Singletons were automatically inserted in the final VCF file using a local Perl script as unphased or phased, depending on the singleton's RBP status.

To generate complete sequences and CDS sequences (only exons) for each HLA gene and each sample, we used the vcfx fasta and vcfx transcript functions. We also compared our sequences with the ones described in the IPD-IMGT/HLA Database version 3.4.0 [13] using a local Perl script, identifying whether they were identical or not to known sequences in the database. The CDS sequences were translated into full-length proteins using EMBOSS transeq and named according to the IPD-IMGT/HLA database. Allele, genotype, and haplotype frequencies were calculated by direct counting. Variants were annotated using SNPeff [14].

### 7. HLA Imputation

**Supplementary Table 18.** Number of samples and alleles in each reference panel (1KGP3, SABE and SABE+1KGP3) and the out-of-bag accuracy for the HLA imputation models with 2 fields resolution.

| Locus | Samples in the reference panel | N alleles in the reference panel (2 fields resolution) | Average Out-of-bag Accuracy |
|---|---|---|---|
| | | 1KGP3 | |
| *HLA-A* | 2503 | 82 | 91.33% ± 0.79% |
| *HLA-B* | 2498 | 154 | 86.36% ± 0.87% |
| *HLA-C* | 2503 | 63 | 97.31% ± 0.41% |
| | | SABE | |
| *HLA-A* | 1171 | 68 | 92.47% ± 0.81% |
| *HLA-B* | 1171 | 107 | 85.61% ± 1.12% |
| *HLA-C* | 1171 | 45 | 97.72% ± 0.53% |
| | | SABE + 1KGP3 | |
| *HLA-A* | 3674 | 102 | 90.28% ± 0.54% |
| *HLA-B* | 3669 | 176 | 86.33% ± 0.62% |
| *HLA-C* | 3674 | 74 | 97.58% ± 0.29% |

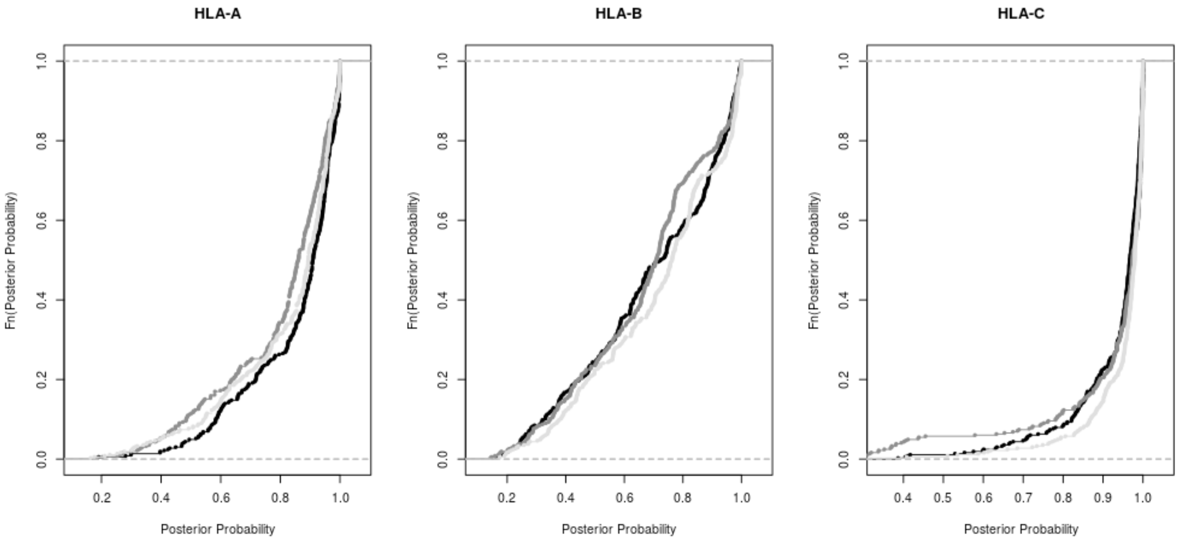

**Supplementary Fig. 23**. Empirical cumulative distribution (ECD) of posterior probabilities for the HLA imputation models: 1KGP3 (black), SABE (dark gray) and SABE+1KGP3 (light gray).

### 8. GWAS

**Supplementary Table 19.** Phenotypes used in GWAS analyses

| Quantitative phenotypes | Total n | Mean value (±stdev) | |
|---|---|---|---|
| Triglycerides (mg/dL) | 1102 | 134.72±87.05 | |
| LDL (mg/dL) | 1102 | 128.61±35.06 | |
| BMI (kg/m2) | 1024 | 28.1±5.24 | |
| **Discrete phenotypes** | | **Cases** | **Controls** |
| | **Total n** | **n (%)** | **n (%)** |
| Diabetes positive history | 1105 | 272 (24.6) | 833 (75.4) |
| Cancer positive history | 1105 | 77 (7) | 1028 (93) |
| Hypertension positive history | 1105 | 758 (68.6) | 347 (31.4) |
| Cognitive decline (≤13 mMMSE)* | 1105 | 152 (13.8) | 953 (86.2) |
| Frailty (≥1 component)** | 1032 | 632 (61.2) | 400 (38.8) |

*mini-Mini Mental State Examination is a reduced version adapted and validated for illiterate individuals[15].
**Frailty is composed of up to 5 phenotypes[16]. Strict sense frail elderly has ≥3 components. Among SABE, there are 110 individuals (10.6%) with 3 or more components.

**Supplementary Table 20.** GWAS hits for cancer. We considered significant, associations with p-values < 10e-09 for whole genomes and with p-values < 5e-8 for array-filtered data.

| Variant ID | Chr:Position (hg38) | Effect allele | Nearest gene: Consequence | EAF SABE1171 | EAF Africans | EAF Europeans | SABE1171-WGS (p-value < e-09)[a] | SABE1171-Array (p-value < 5e-08)[a] |
|---|---|---|---|---|---|---|---|---|
| rs73902781 | 21:36881365 | T | *HLCS*:intronic | 0.018 | 0.056 | 0.0038 | 1.8e-09 (F) | - |

EAF: Effect allele frequency; EAF for Africans and Europeans from gnomAD. [a](A): hit using all individuals, (F): hit using females, (M): hit using males.

**Supplementary Table 21**. GWAS hits for BMI. We considered significant, associations with p-values < 10e-09 for whole genomes and with p-values < 5e-8 for array-filtered data.. SNPs in red have $r^2 > 0.95$ ($0.95 < r^2 < 0.99$).

| Variant ID | Chr:Position (hg38) | Effect allele | Nearest gene: Consequence | EAF SABE1171 | EAF Africans | EAF Europeans | SABE1171-WGS (p-value < e-09)[a] | SABE1171-Array (p-value < 5e-08)[a] |
|---|---|---|---|---|---|---|---|---|
| rs77326660 | 2:106346360 | C | *ILRUNP1/ANAPC1P6*:intergenic | 0.014 | 0.076 | 0.0002 | 8.4e-09 (A) 7.2e-09 (F) | - |
| rs112879838 | 6:19582361 | A | *LOC105374958/LOC105374963*:intergenic | 0.0054 | 0.026 | 0.0002 | 5.3e-09 (A) | - |
| rs187130645 | 6:38222539 | G | *BTBD9*:intronic | 0.0054 | 0.014 | 0.0047 | 5.0e-09 (F) | - |
| rs35322794 | 6:100381956 | T | *LOC100129854/SIM1*: intergenic | 0.1232 | 0.097 | 0.1270 | 7.8e-09 (M) | - |
| rs13209911 | 6:100368601 | T | *LOC100129854/SIM1*: intergenic | 0.1211 | 0.080 | 0.1266 | - | 2.0e-08 (M) |
| rs13201004 | 6:100388908 | A | *IM1*:3 Prime UTR Variant | 0.1214 | 0.095 | 0.1268 | - | 3.2e-08 (M) |
| rs10843058 | 12:28019612 | A | *PTHLH/LOC105369710*:intergenic | 0.0742 | 0.027 | 0.1046 | 6.3e-09 (A) | - |
| rs2619433 | 12:28013124 | G | *PTHLH/LOC105369710*:intergenic | 0.075 | 0.027 | 0.1044 | - | 2.0e-08 (A) |
| rs148632533 | 18:11308150 | T | *PIEZO2/LOC100130329*:intergenic | 0.0144 | 0.001 | 0.0004 | 1.3e-09 (M) | - |

EAF: Effect allele frequency; EAF for Africans and Europeans from gnomAD. [a](A): hit using all individuals, (F): hit using females, (M): hit using males.

**Supplementary Table 22.** GWAS hits for LDL. We considered significant, associations with p-values < 10e-09 for whole genomes and with p-values < 5e-8 for array-filtered data.. SNPs in red have $r^2 > 0.52$ ($0.52 < r^2 < 0.95$). SNPs in blue have $r^2 > 0.35$ ($0.35 < r^2 < 0.64$).

| Variant ID | Chr:Position (hg38) | Effect allele | Nearest gene: Consequence | EAF SABE1171 | EAF Africans | EAF Europeans | SABE1171-WGS (p-value < e-09) | SABE1171-Array (p-value < 5e-08) |
|---|---|---|---|---|---|---|---|---|
| rs114557300 | 1:34295191 | T | *LOC105378639*:Intronic | 0.0099 | 0.051 | 0.0002 | 1.1e-10 (A) | - |
| rs112794427 | 1:34273959 | T | *C1orf94/LOC100420336*: intergenic | 0.009 | 0.049 | 0.0001 | 2.3e-10 (A) | - |
| rs2140686 | 1:34306903 | A | *LOC105378639/LOC105378640*:intergenic | 0.0117 | 0.055 | 0.0001 | 2.3e-09 (A) | 2.3e-09 (A) |
| rs115111275 | 1:34289979 | T | *LOC105378639*:intronic | 0.0135 | 0.054 | 0.0097 | 2.3e-09 (A) | - |
| rs4653019 | 1:34299118 | T | *LOC105378639/LOC105378640*:intergenic | 0.0162 | 0.056 | 0.0097 | 9.9e-09 (A) | - |
| rs1589800 | 1:34305463 | G | *LOC105378639/LOC105378640*:intergenic | 0.0171 | 0.056 | 0.0097 | - | 2.1e-08 (A) |
| rs4131235 | 1:81444514 | T | *ADGRL2*:intronic | 0.0235 | 0.044 | 0.0127 | - | 3.5e-08 (A) |
| rs532144565 | 12:120346670 | G | *MSI1*:intronic | 0.0067 | 0.000 | 0.0014 | 4.4e-09 (F) | - |
| rs113548199 | 12:2136299 | C | *CACNA1C*:intronic *LOC107984131*:intronic | 0.019 | 0.075 | 0.0004 | 1.4e-09 (M) | - |
| rs74807647 | 12:2137484 | T | *CACNA1C*:intronic *LOC107984131*:intronic | 0.0104 | 0.040 | 0.0001 | 3.1e-09 (M) | - |
| rs112586389 | 12:2124147 | C | *CACNA1C*:intronic | 0.0144 | 0.065 | 0.0002 | 5.5e-09 (M) | - |
| rs115676766 | 16:74924982 | T | *WDR59*:intronic | 0.0076 | 0.077 | 0.0001 | 1.2e-09 (A) | - |

EAF: Effect allele frequency; EAF for Africans and Europeans from gnomAD. [a](A): hit using all individuals, (F): hit using females, (M): hit using males.

**Supplementary Table 23.** GWAS hits for triglycerides. We considered significant, associations with p-values < 10e-09 for whole genomes and with p-values < 5e-8 for array-filtered data.

| Variant ID | Chr:Position (hg38) | Effect allele | Nearest gene: Consequence | EAF SABE1171 | EAF Africans | EAF Europeans | SABE1171 -WGS (p-value < e-09) | SABE1171-Array (p-value < 5e-08) |
|---|---|---|---|---|---|---|---|---|
| rs76305592 | 13:20967039 | A | *RPSAP54/LATS2*:intergenic | 0.0108 | 0.043 | 0.0002 | 1.9e-09 (F) | - |
| rs478470 | 20:4414505 | C | *ADRA1D/RPL7AP12*:intergenic | 0.0085 | 0.031 | 0.0000 | 1.9e-09 (A) 1.9e-09 (F) | - |

EAF: Effect allele frequency; EAF for Africans and Europeans from gnomAD. [a](A): hit using all individuals, (F): hit using females, (M): hit using males.

**A**

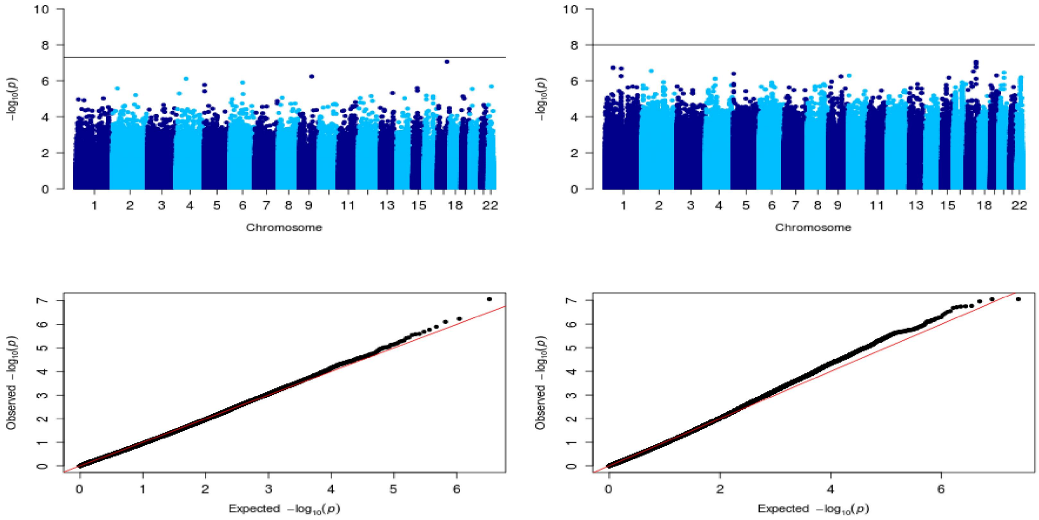

**B**

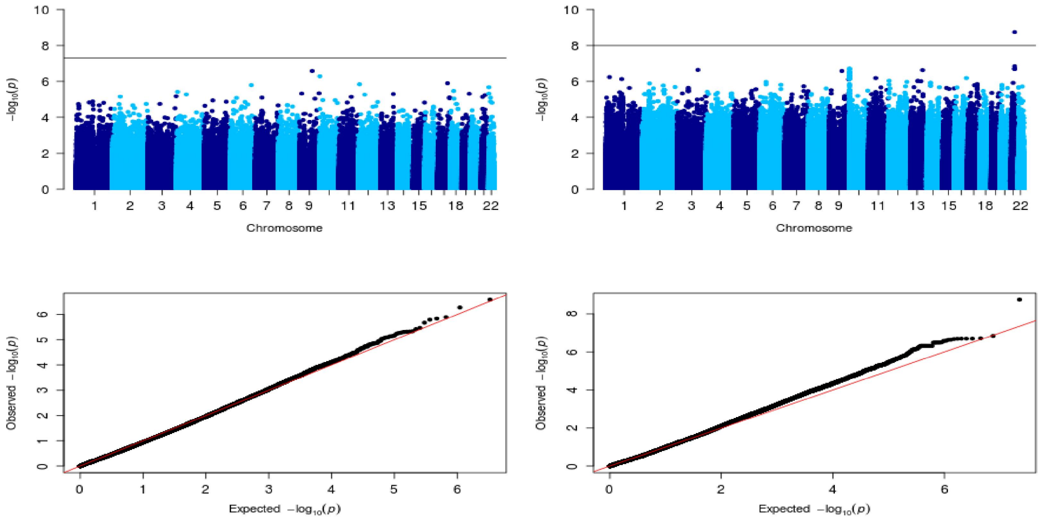

**C**

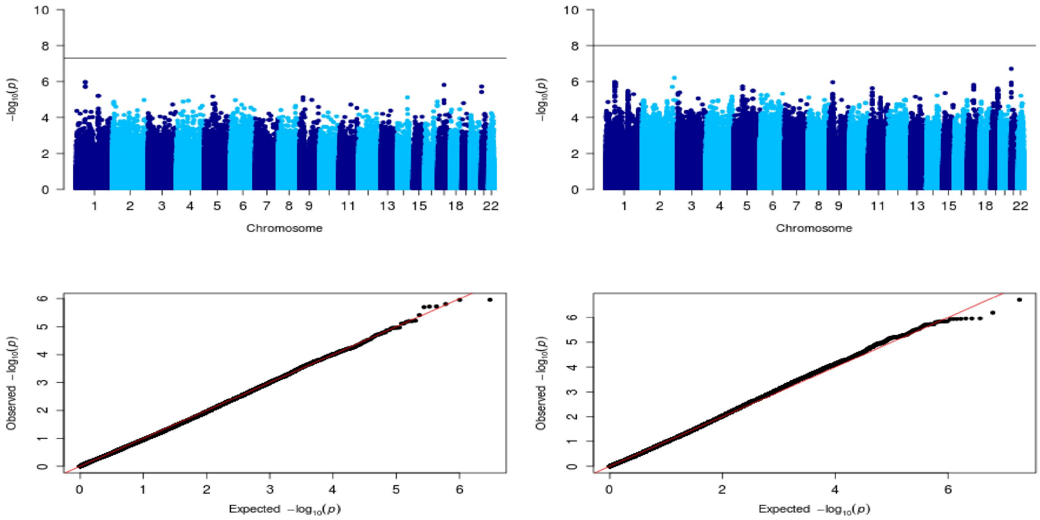

**Supplementary Fig. 24**. Manhattan and qq plots from GWAS analysis for **cancer** using: **A.** all individuals, **B.** females, **C.** males. Left panels: SABE1171-Array, Right panels: SABE1171-WGS. Black line in the Manhattan plots corresponds to the threshold p-value.

**A**

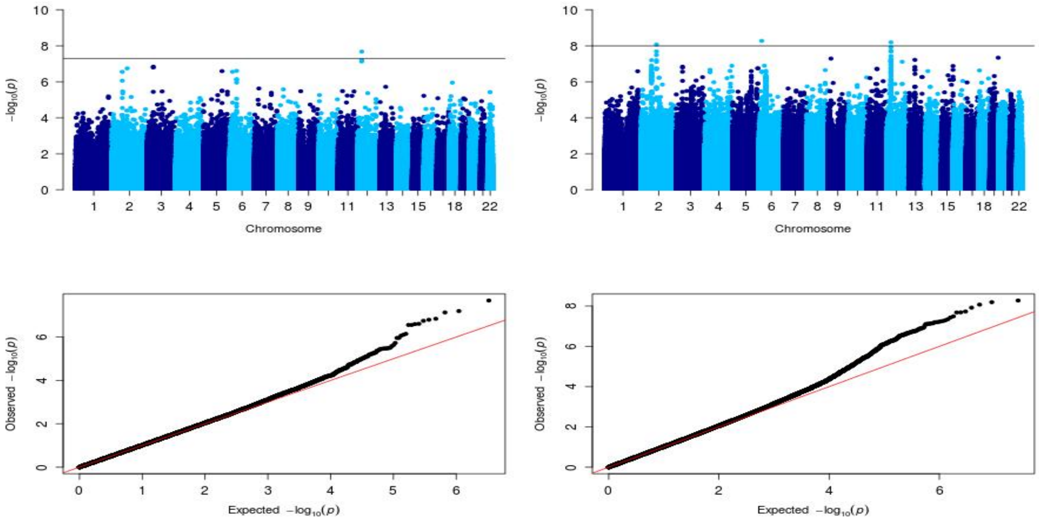
-log10(p)
Chromosome
Observed -log10(p)
Expected -log10(p)

**B**

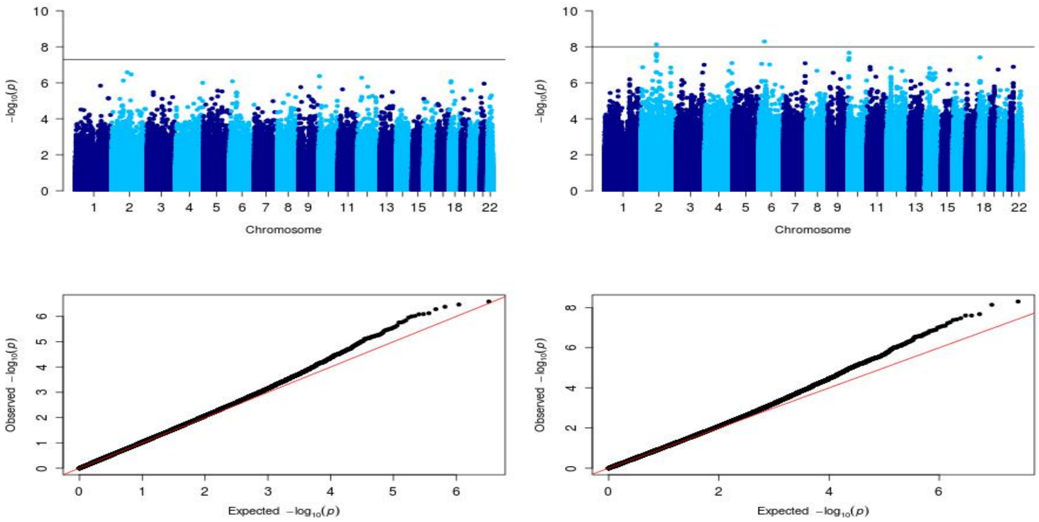
-log10(p)
Chromosome
Observed -log10(p)
Expected -log10(p)

**C**

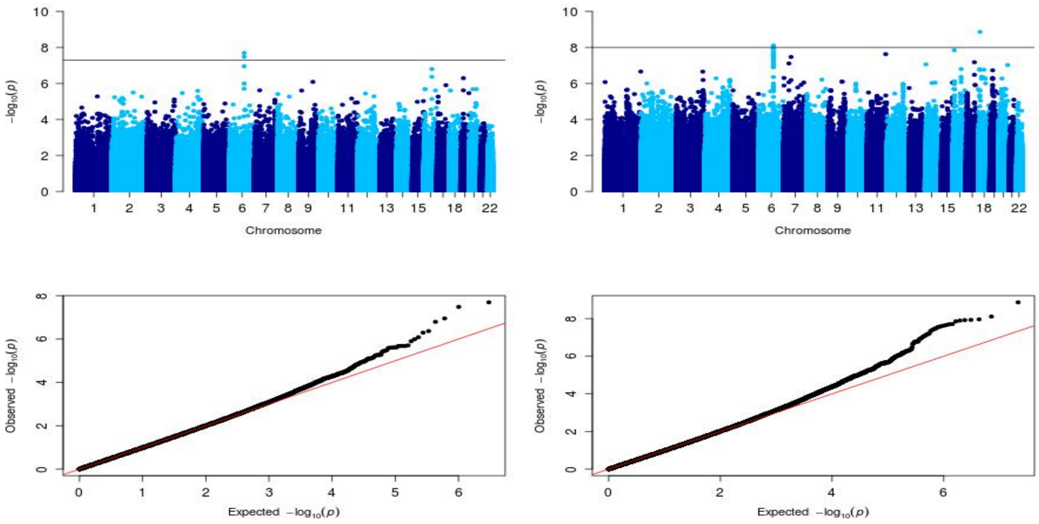
-log10(p)
Chromosome
Observed -log10(p)
Expected -log10(p)

**D**

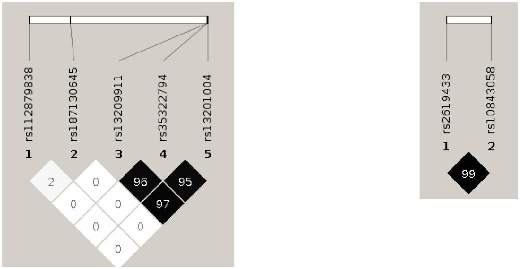

**Supplementary Fig. 25**. Manhattan and qq plots from GWAS analysis for **BMI** using: **A.** all individuals, **B.** females, **C.** males. Left panels: SABE1171-Array, Right panels: SABE1171-WGS. Black line in the Manhattan plots corresponds to the threshold p-value. (D) plot by Haploview.showing $r^2$ among associated SNPs.

A

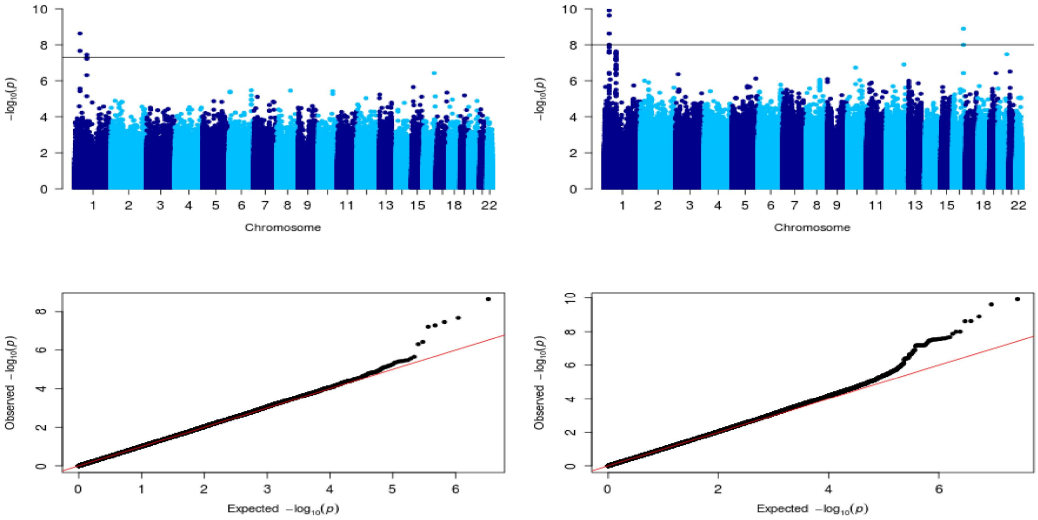
Chromosome
Chromosome
$-\log_{10}(p)$
Observed $-\log_{10}(p)$
Expected $-\log_{10}(p)$

B

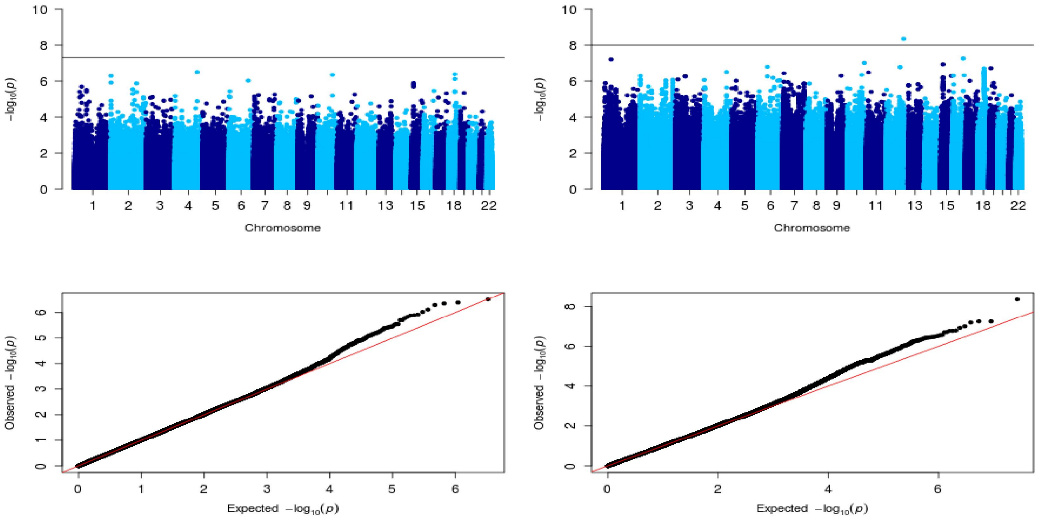
Chromosome
Chromosome
$-\log_{10}(p)$
Observed $-\log_{10}(p)$
Expected $-\log_{10}(p)$

C

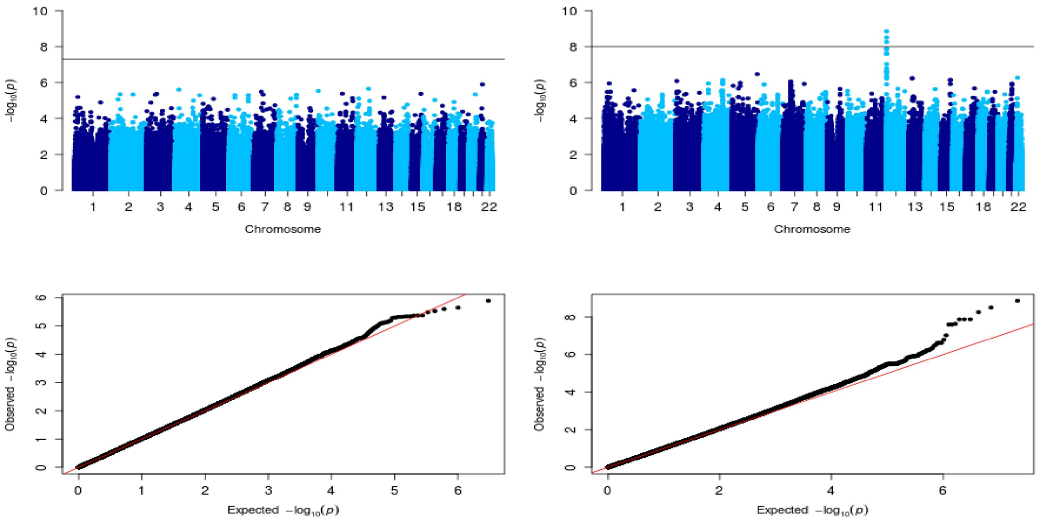
Chromosome
Chromosome
$-\log_{10}(p)$
Observed $-\log_{10}(p)$
Expected $-\log_{10}(p)$

**D**

**Supplementary Fig. 26**. Manhattan and qq plots from GWAS analysis for **LDL** using: **A.** all individuals, **B.** females, **C.** males. Left panels: SABE1171-Array, Right panels: SABE1171-WGS. Black line in the Manhattan plots corresponds to the threshold p-value. (D) plot by Haploview showing $r^2$ among associated SNPs.

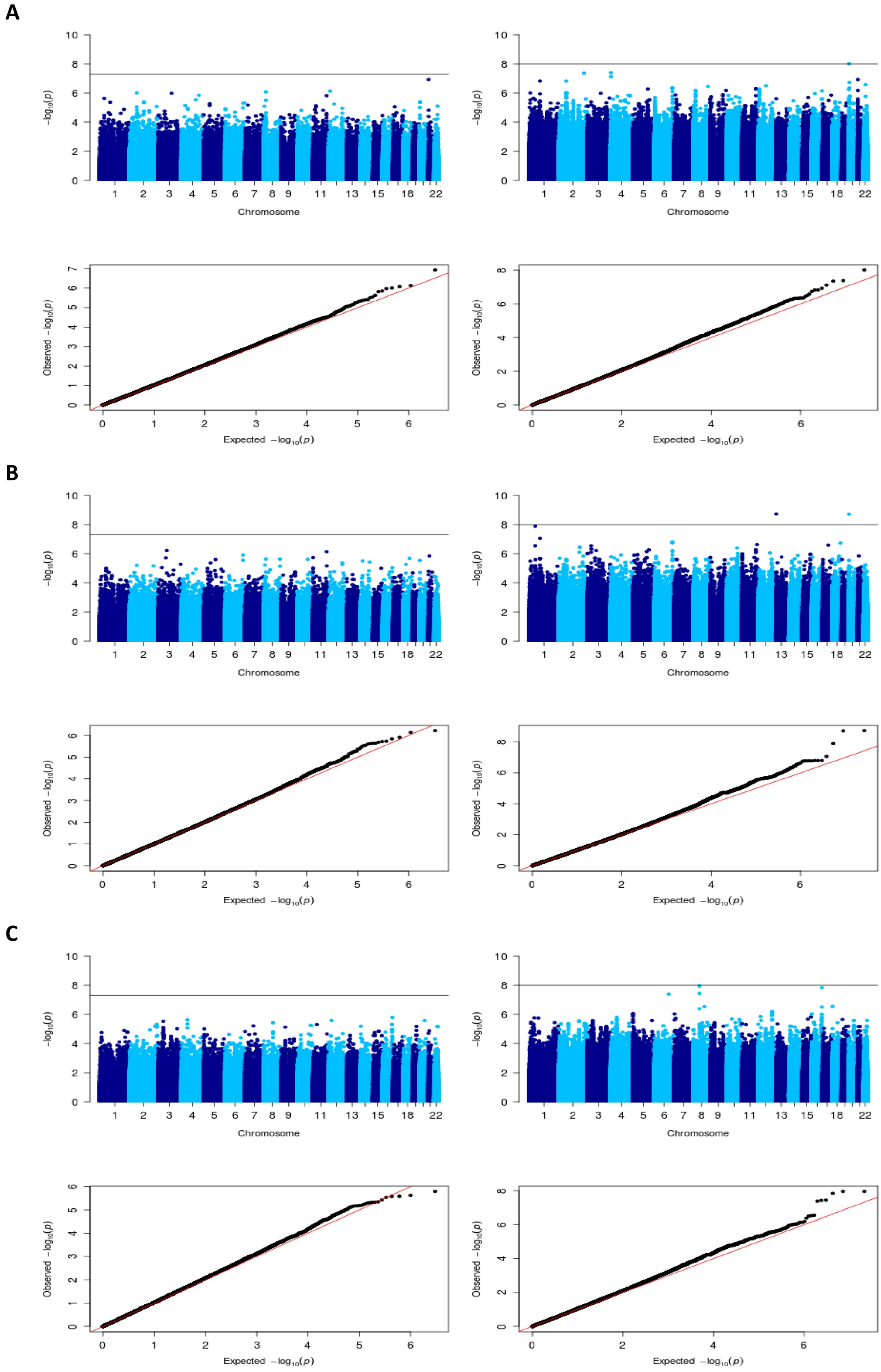

**Supplementary Fig. 27**. Manhattan and qq plots from GWAS analysis for **triglycerides** using: **A.** all individuals, **B.** females, **C.** males. Left panels: SABE1171-Array, Right panels: SABE1171-WGS. Black line in the Manhattan plots corresponds to the threshold p-value.

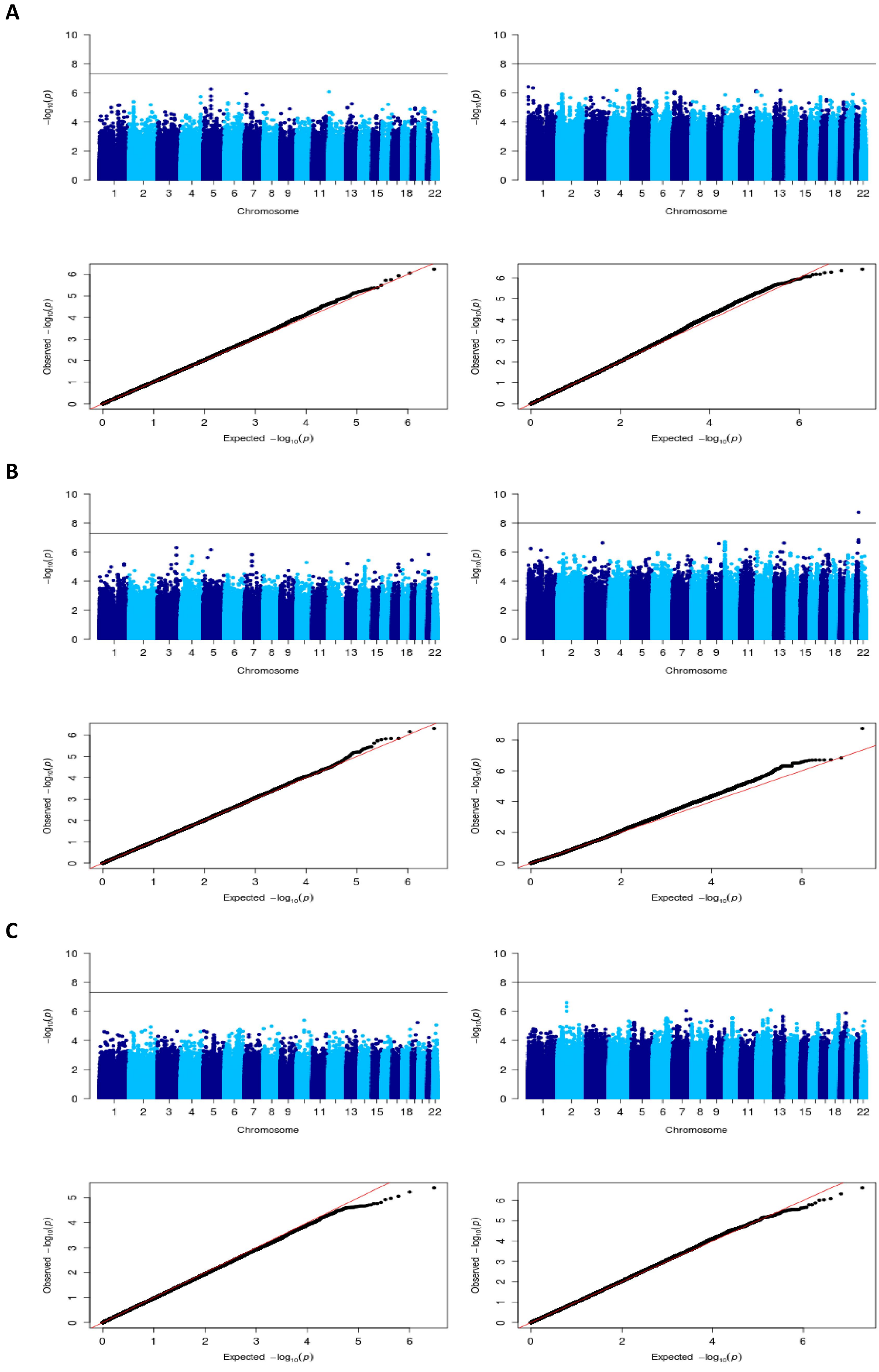

**Supplementary Fig. 28.** Triglycerides Manhattan and qq plots from GWAS analysis for **cognitive** using: **A.** all individuals, **B.** females, **C.** males. Left panels: SABE1171-Array, Right panels: SABE1171-WGS. Black line in the Manhattan plots corresponds to the threshold p-value.

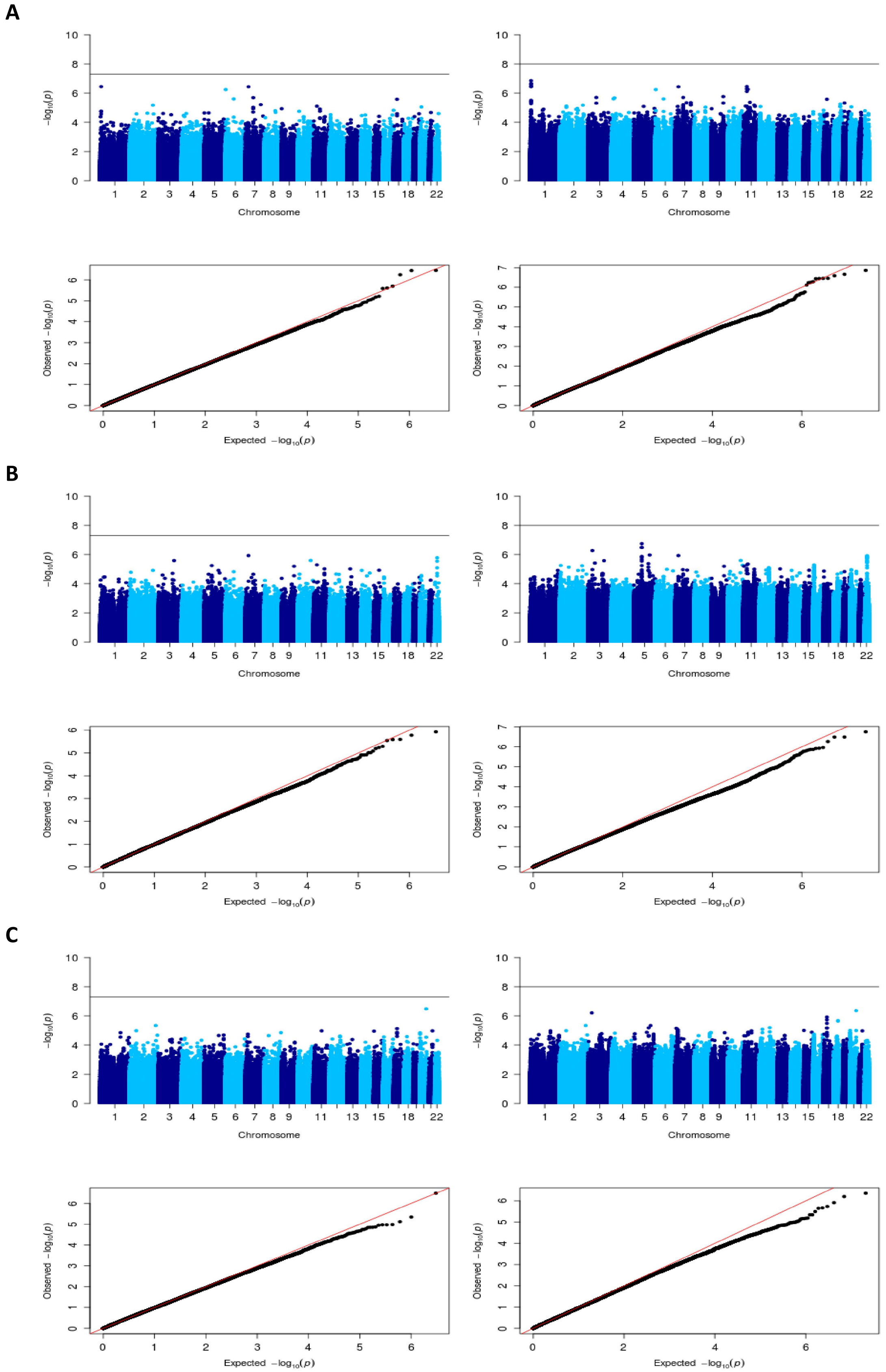

**Supplementary Fig. 29.** Manhattan and qq plots from GWAS analysis for **diabetes** using: **A.** all individuals, **B.** females, **C.** males. Left panels: SABE1171-Array, Right panels: SABE1171-WGS. Black line in the Manhattan plots corresponds to the threshold p-value.

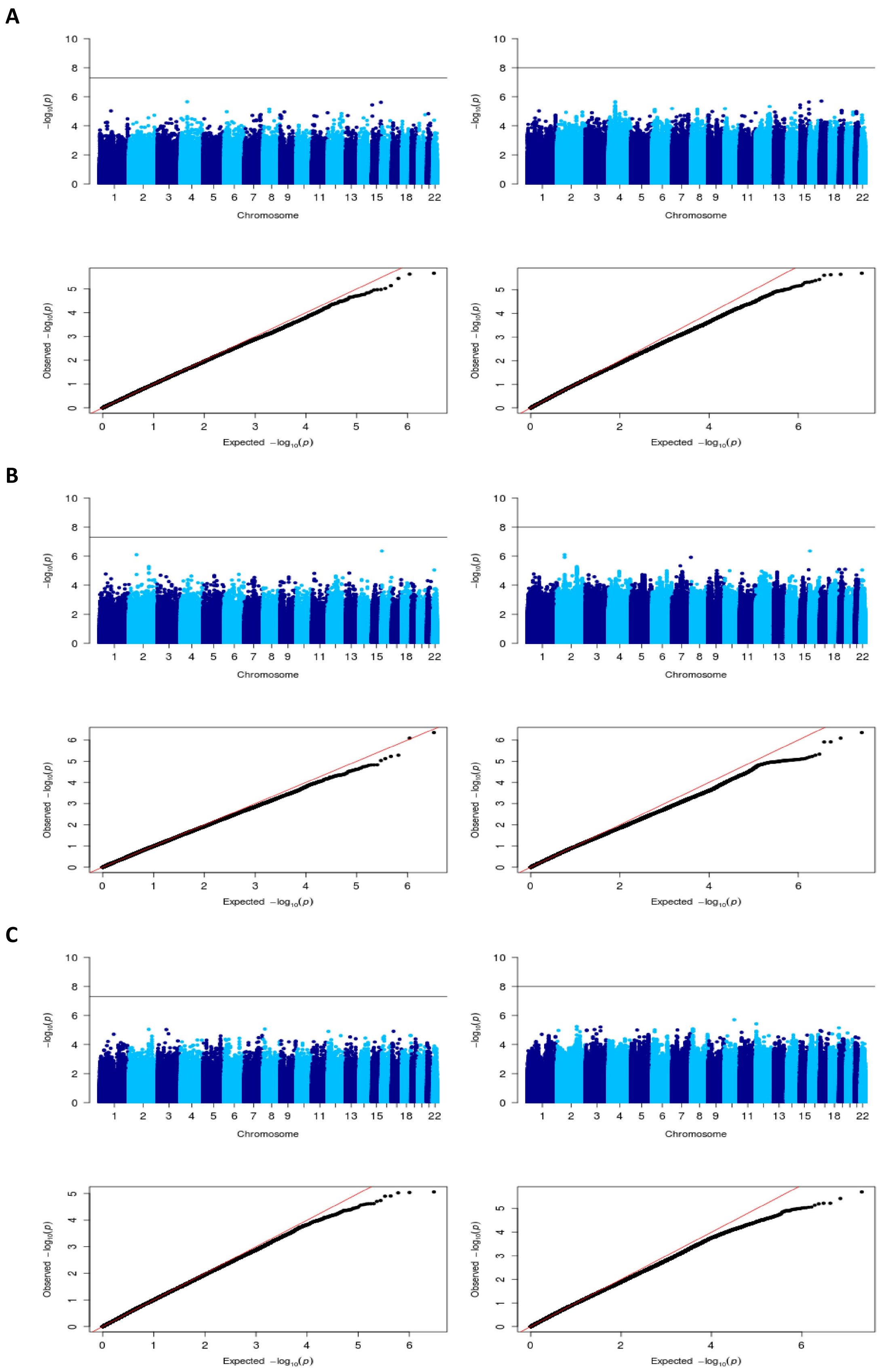

**Supplementary Fig. 30.** Manhattan and qq plots from GWAS analysis for **frailty** using: **A.** all individuals, **B.** females, **C.** males. Left panels: SABE1171-Array, Right panels: SABE1171-WGS. Black line in the Manhattan plots corresponds to the threshold p-value.

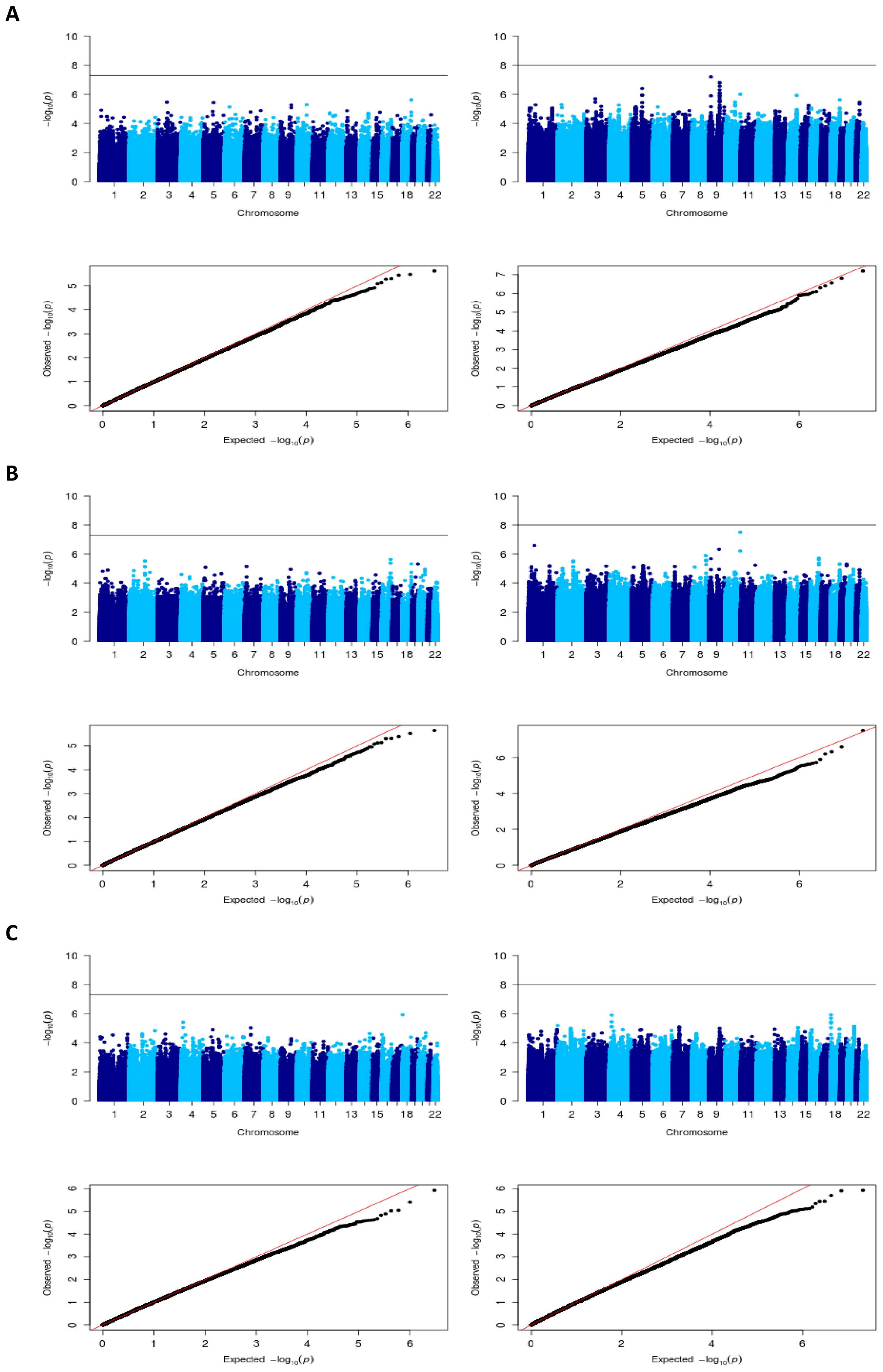

**Supplementary Fig. 31.** Manhattan and qq plots from GWAS analysis for **hypertension** using: **A.** all individuals, **B.** females, **C.** males. Left panels: SABE1171-Array, Right panels: SABE1171-WGS. Black line in the Manhattan plots corresponds to the threshold p-value.
